# On a Reef Far, Far Away: Anthropogenic Impacts Following Extreme Storms Affect Sponge Health and Bacterial Communities

**DOI:** 10.1101/2020.04.27.064568

**Authors:** Amanda Shore, Jordan A. Sims, Michael Grimes, Lauren I. Howe-Kerr, Carsten G.B. Grupstra, Shawn M. Doyle, Lauren Stadler, Jason B. Sylvan, Kathryn E.F. Shamberger, Sarah W. Davies, Lory Z. Santiago-Vázquez, Adrienne M.S. Correa

**Author notes:** **Correspondence:** Amanda Shore.

## Abstract

Terrestrial runoff can negatively impact marine ecosystems through stressors including excess nutrients, freshwater, sediments, and contaminants. Severe storms, which are increasing with global climate change, generate massive inputs of runoff over short timescales (hours to days); such runoff impacted offshore reefs in the northwest Gulf of Mexico (NW GoM) following severe storms in 2016 and 2017. Several weeks after coastal flooding from these events, NW GoM reef corals, sponges, and other benthic invertebrates -185 km offshore experienced mortality (2016 only) and/or sub-lethal stress (both years). To assess the impact of storm-derived runoff on reef filter feeders, we characterized the bacterial communities of two sponges, *Agelas clathrodes* and *Xestospongia muta*, from offshore reefs during periods of sub-lethal stress and no stress over a three-year period (2016-2018). Sponge-associated bacterial communities were altered during both flood years. Additionally, we found evidence of wastewater contamination (based on 16S rRNA gene libraries and quantitative PCR) in offshore sponges following these flood years, but not during the no-flood year. We show that flood events from severe storms have the capacity to reach offshore reef ecosystems and impact resident benthic organisms. Such impacts are most readily detected if baseline data on organismal physiology and associated microbiome composition are available. This highlights the need for molecular and microbial time series of benthic organisms in near- and offshore reef ecosystems, and the continued mitigation of stormwater runoff and climate change impacts.

## INTRODUCTION

Tropical coral reef ecosystems have evolved in the context of nutrient-poor (oligotrophic) waters. Thus, when nutrient-laden (eutrophic) terrestrial runoff mixes with reef-associated waters, this can directly or indirectly stress or kill reef organisms (Knight and Fell, 1987; Kerswell and Jones, 2003; Fabricius, 2005; Humphrey et al., 2008). Terrestrial runoff exposes reef organisms to decreased salinity, and increased levels of turbidity and contaminants (e.g., microbial pathogens, chemical pollutants) (Fabricius, 2005). Terrestrial runoff can also reduce dissolved oxygen levels in reef-associated waters through several mechanisms (Nelson and Altieri, 2019). Terrestrial runoff can constitute a chronic stress in areas with developed coastlines or at river outflows, or an acute stress when floodwaters generated by extreme storms move offshore. Floodwaters originating in urban areas may constitute a greater threat, if they contain high nutrient and contaminant loads due to overflows of wastewater management systems (Chen et al., 2019; Humphrey et al., 2019). Storm-associated runoff is increasingly recognized as a threat to reefs since the intensity and precipitation associated with tropical storms is increasing with climate change (Knutson et al., 2010; Emanuel, 2017).

Although various studies document the impacts of terrestrial runoff on coral reefs, few works specifically address runoff derived from floodwaters; all of these latter works focus on nearshore, shallow water reefs (Ostrander et al., 2008; Lapointe et al., 2010). It has been assumed that offshore or deep (e.g., mesophotic) reefs are unlikely to interact with terrestrial runoff (Szmant, 2002). For example, reefs within the Flower Garden Banks (FGB) National Marine Sanctuary (northwest Gulf of Mexico, Figure 1a) occur ∼185 km offshore in relatively deep water (20-30 m), boast some of the highest coral cover (∼55%) in the wider Caribbean (Johnston et al., 2016), and have been generally presumed to be protected from land-based stressors (Johnston et al., 2016). Low salinity (< 33 parts per thousand) water from terrestrial runoff has been detected within surface waters of the FGB for the past several decades (Dodge and Lang, 1983; Deslarzes and Lugo-Fernandez, 2007; Kealoha, 2019); however, the impact of these waters on reef ecosystem health has not been directly studied.

**Figure 1.**
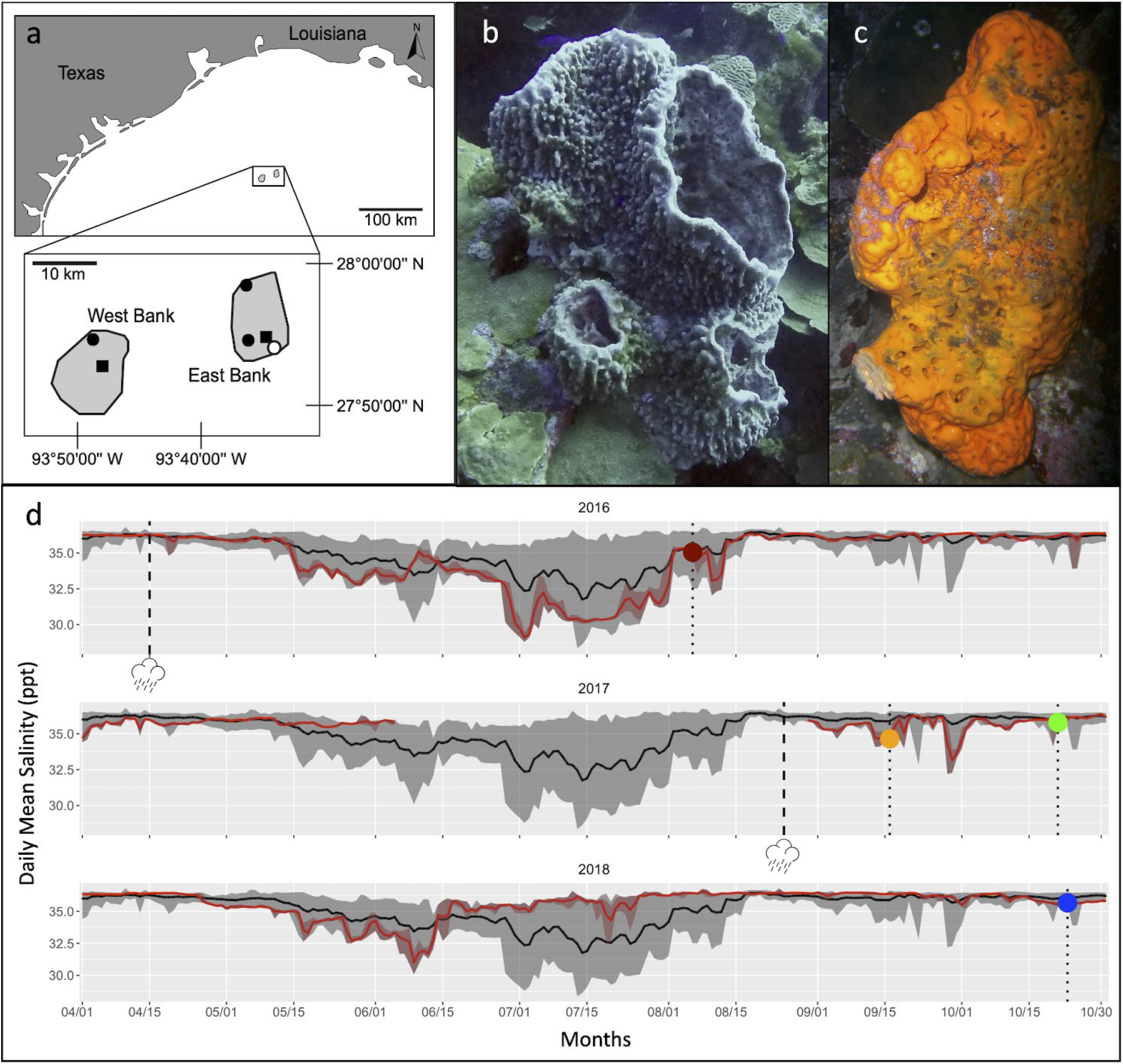
Summary of study site, host taxa and local abiotic conditions associated with this study. (A) Map of Flower Garden Banks National Marine Sanctuary (northwest Gulf of Mexico) with sites of sponge and seawater collection indicated as black shapes. Filled circles represent seawater column sample collection sites; filled squares represent Buoy 2 and Buoy 4 (sponge, seawater column, and near reef seawater collection) and open circle represents location of TABS Buoy V (surface salinity data collection); (B) Representative *Xestospongia muta* sponge; (C) Representative *Agelas clathrodes* sponge; (D) Surface Salinity (ppt) at buoy V spanning the months April through October in which sampling occurred for this study. Black lines represent daily means from 2006–2018 and grey shaded areas encompass minimum and maximum values from 2006–2018. Red lines represent daily means for 2015, 2016, 2017, and 2018, individually. Dashed lines with storm icon represent dates of terrestrial flooding. Dotted lines with symbols represent mean daily values on each sample collection date: dark red circle = 6 August 2016, orange circle = 16 September 2017, green circle = 21 October 2017, and blue circle = 23 October 2018.

Since 2015, the Texas coast has experienced several flooding events related to extreme storms: Memorial Day Flood of 2015; Tax Day Flood of 2016, Hurricane Harvey in 2017, and Hurricane Imelda in 2019. Each of these floods impacted Central Texas nearshore ecosystems, including salt marshes (Congdon et al., 2019; Oakley and Guillen, 2019) and oyster beds (Kiaghadi and Rifai, 2019). It has been assumed that floodwaters would not impact the benthic reef ecosystem at 20-30 m depth; however, three months after Tax Day flooding in 2016, a localized mortality event occurred on a portion of the East Bank (EB) of the FGB. During the mortality event, approximately 82% of corals in a 0.06 km^2^ area experienced partial or full mortality (Johnston et al., 2019), and mortality in many other benthic invertebrates, such as sponges, was also observed. Although data on abiotic conditions on the reef at EB during the 2016 mortality event are not available, measurements from nearby sites suggest that poor water quality from floodwaters moving offshore and low dissolved oxygen levels played a role in the mortality event (Le Hénaff et al., 2019; Kealoha et al., 2020). Then, in late August of 2017, Hurricane Harvey released more than one meter of rainfall over the course of six days in some areas of southeastern Texas (Blake and Zelinsky, 2018). Although surface salinity was slightly depressed near the FGB following Hurricane Harvey, much of the water mass diverted southwest along the coast (Roffer et al., 2018), and no mass mortality was observed on the reef (Wright et al., 2019).

Benthic reef invertebrates, such as sponges, harbor a diversity of microbial symbionts (e.g., bacteria, archaea, protists, fungi and viruses) that contribute to their health and nutrition. Yet, these microbial communities can be disrupted by environmental stress, including terrestrial runoff (Slaby et al., 2019). Previous work has shown the utility of using filter-feeding sponges as monitoring tools for fecal-coliform contamination in near-shore environments (Longo et al., 2010; Maldonado et al., 2010). Therefore, we tested whether changes to microbial symbioses in offshore reef sponges were detectable following storm-derived coastal flood events. We leverage the Tax Day Flood (2016) and Hurricane Harvey (2017) as natural ‘experimental treatments’ applied to two high-microbial-abundance sponge species (*Xestospongia muta* and *Agelas clathrodes*; Figure 1b-c) at the EB and West Bank (WB) of the FGB. Bacterial communities were sampled from sponges and seawater at four time points: August 2016 (two weeks after the localized mortality event at EB), September 2017 (immediately after Hurricane Harvey), October 2017 (one month after Hurricane Harvey), and October 2018 (approximately one year following Hurricane Harvey) (Figure 1d). No flood events occurred in southeast central Texas during 2018, and thus samples from this time point function as an ‘experimental baseline’. We hypothesized that: (1) sponge-associated bacterial communities shift during flood years (relative to the no-flood year) and (2) flood year bacterial communities contain genetic signatures of terrestrial-derived bacteria. Understanding how and when environmental stressors influence sponge-microbe associations, and the subsequent implications for sponge health and function, is important as sponges increase in abundance and in ecological importance on Caribbean reefs (Bell et al., 2013; Pawlik and McMurray, 2020).

## RESULTS

### Reduced surface salinity at the FGB following floods

In the vicinity of the FGB, mean surface salinity is generally variable, ranging between 28.5 ppt and 36 ppt, from early-June through late-August (shaded grey areas in Figure 1d). In early spring and in fall, however, surface salinity is more consistent (∼33-36 ppt). Approximately 30 days after the 2016 Tax Day Flood impacted Texas, a period of depressed mean surface salinity (relative to the mean surface salinity for a thirteen-year period: 2006-2018) was recorded from 16 May 2016 to 12 August 2016 at Texas Automated Buoy System (TABS) Real Time Ocean Observations, Buoy V (located within the sanctuary boundaries of EB, open circle Figure 1a). Average surface salinity was 1.2 ppt lower than the thirteen-year mean during this period; the most significant deviation occurred around 2 July 2016 when mean surface salinity reached a minimum of 29.1 ppt (3.8 ppt below the thirteen-year mean). In 2017, surface salinity remained > 35 ppt until early June. TABS Buoy V data are unavailable for much of June to August 2017, so it is unclear how surface salinity changed in the months preceding Hurricane Harvey. Two abrupt reductions in surface salinity (to 32.6 ppt and 32.7 ppt) were recorded near the FGB during mid-to-late September 2017 following Hurricane Harvey, before surface salinity returned to > 35 ppt in October. In contrast, no significant influence of freshwater was recorded in 2018 and surface salinity remained above 35 ppt after mid-June.

When the 2016 mortality event was discovered on 25 July 2016, recreational divers reported ‘low visibility and green coloration in the water column’ at EB (Johnston et al., 2019); on this date, mean surface salinity had been lower than the thirteen-year mean for 44 days. Collections of live but visually impacted sponges as well as healthy sponges were collected on 6 August 2016, approximately four days after surface salinity had returned to 35 ppt (Figure 1d). Sampling immediately post-Hurricane Harvey on 16 September 2017 occurred when mean surface salinity was 34.6 ppt, or 1.3 ppt lower than the thirteen-year mean (Figure 1d). During sampling that occurred approximately one-month post-hurricane Harvey (on 21 October 2017), mean surface salinity had been > 35 ppt for 19 days prior to sampling (Figure 1d). Samples during the no-flood year were collected on 23 October 2018, which had a mean surface salinity of 35.7 ppt; surface salinity had been > 35 ppt for 94 days (Figure 1d).

### Acute shifts in sponge- and seawater-associated bacterial communities following flood events

Paired-end Illumina MiSeq sequencing of the V4 region of the bacterial 16S rRNA gene yielded 5,350,216 high quality sequences from a total of 139 samples that included healthy and impacted sponges (*A. clathrodes*, n=54; *X. muta*, n=45) as well as seawater (n=40) (Table 1, Supplemental Table S1). After removal of Mitochondrial, Chloroplast, and Unassigned reads, the total pool of bacterial sequences were assigned to 11,828 Amplicon Sequence Variants (ASVs). Samples clustered into 4 distinct groups (Figure 2), which were all significantly different from each other (Analysis of Similarities (ANOSIM): p < 0.001; Supplemental Table S2). These groups were: 1) seawater samples; 2) diseased sponges of both species; 3) visually healthy *A. clathrodes*; and 4) visually healthy *X. muta*. Because each sponge species and the seawater samples had distinct bacterial communities, subsequent analyses were conducted on each group individually. Additionally, within each sponge species and within seawater samples, there were no differences between EB and WB sites (ANOSIM: p > 0.05; Supplemental Table S3); therefore, sites were grouped for subsequent beta-diversity analyses.

**Table 1.**
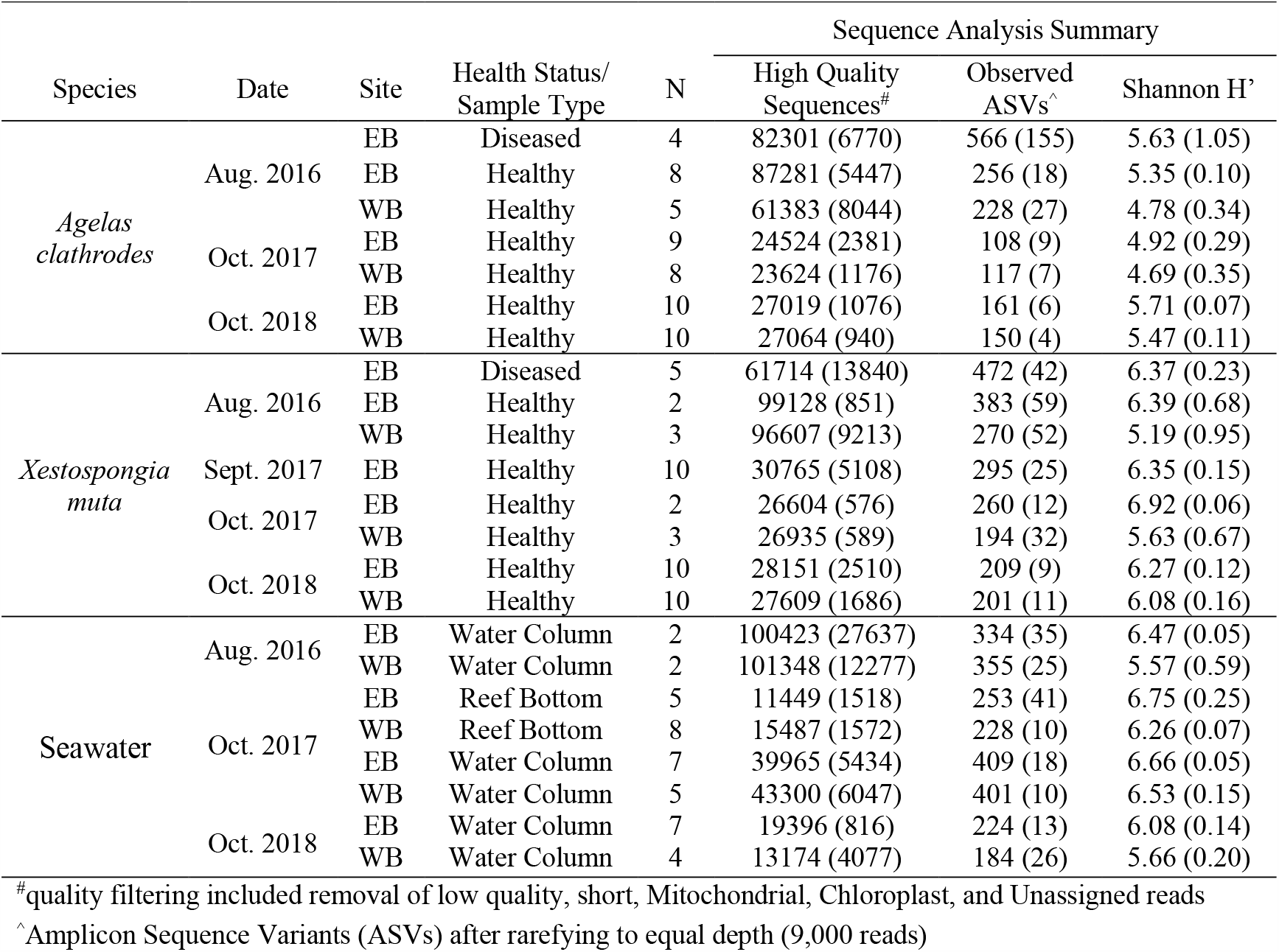
Summary of sample collections from two reef sponge species and seawater at the East Bank (EB) and West Bank (WB) of the Flower Garden Banks National Marine Sanctuary (northwest Gulf of Mexico) and amplicon sequencing results of the V4 region of the 16S rRNA gene from sponge-associated bacterial communities. Richness (Observed ASVs^^^) and diversity (Shannon H’) were calculated from rarefied ASV tables. Data are presented as mean ± (sem).

**Figure 2.**
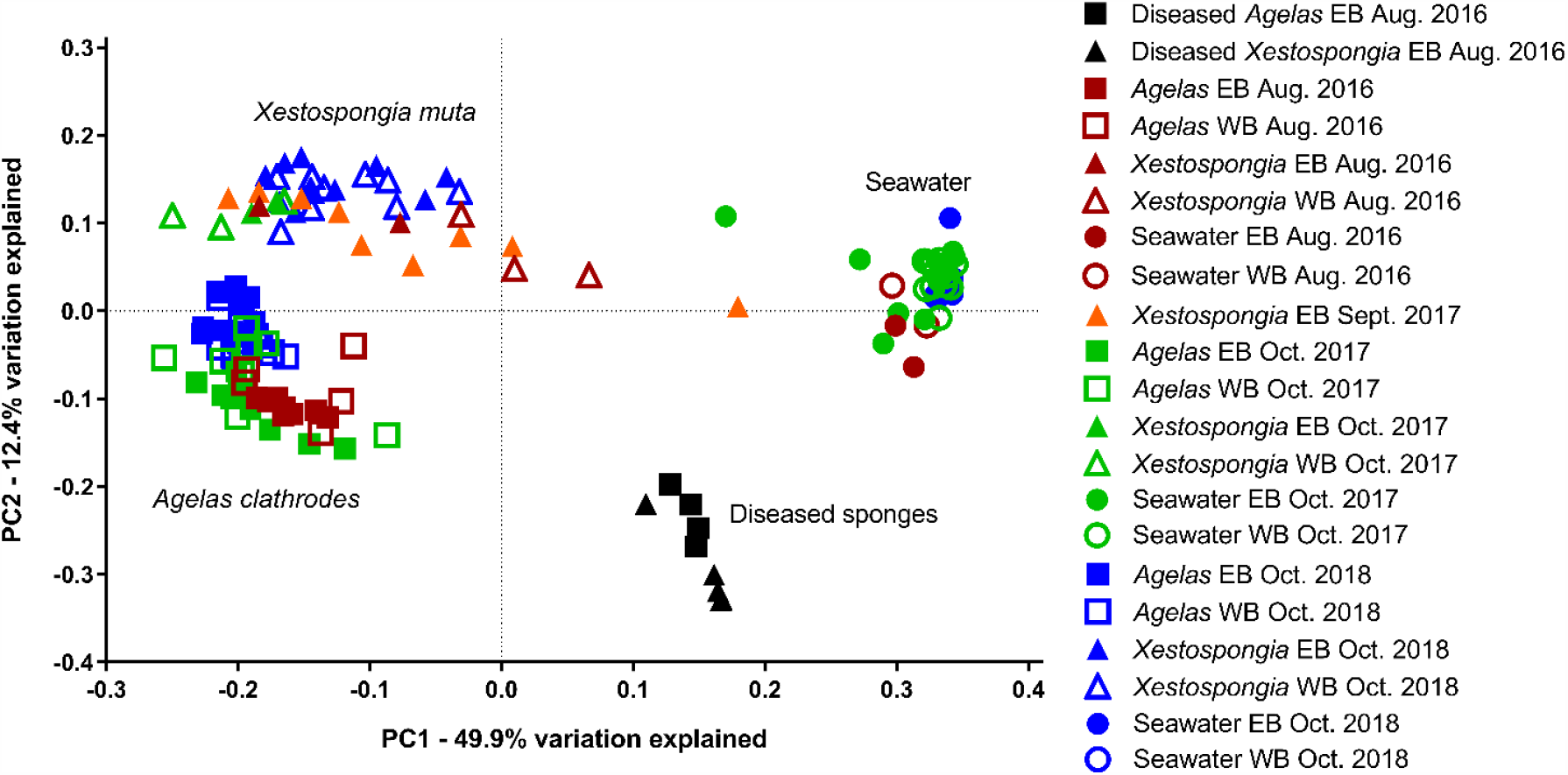
Principle Coordinate Analysis of the weighted UniFrac distance matrix for bacterial communities analyzed in this study. Empty symbols = *Agelas clathrodes* samples; filled symbols = *Xestospongia muta* samples. Black symbols = samples from diseased sponges (August 2016); Red symbols = visually healthy sponge samples (August 2016). Orange symbols = samples collected immediately following Hurricane Harvey (Sept. 2017) from visually healthy sponges; Green symbols = samples collected one month following Hurricane Harvey (Oct. 2017) from visually healthy sponges. Blue symbols = samples collected in a no-flood (baseline) year from visually healthy sponges (Oct. 2018). Circles = samples collected from East Bank (EB); squares = samples collected from West Bank (WB) of the Flower Garden Banks National Marine Sanctuary (northern Gulf of Mexico).

Bacterial communities of *X. muta, A. clathrodes*, and seawater samples were all significantly disrupted during both flood years, relative to the no-flood year. Within *X. muta* samples, bacterial communities in Aug. 2016 and Sept. 2017 shifted in a similar way and Oct. 2017 shifted in another; all three flood-associated dates were significantly different than the no-flood year (Figure 3a; ANOSIM: p < 0.05, Supplemental Table S4). In *A. clathrodes*, bacterial communities associated with flood years shifted in different ways (Figure 3a; ANOSIM: p < 0.05, Supplemental Table S4). In seawater samples, bacterial communities were also distinct in both flood years, and, interestingly, samples taken directly above the reef bottom (‘Reef Water’) differed from samples taken from higher in the water column on the same date (Oct. 2017) (ANOSIM: R = 0.516, p = 0.002; Supplemental Table S4).

**Figure 3.**
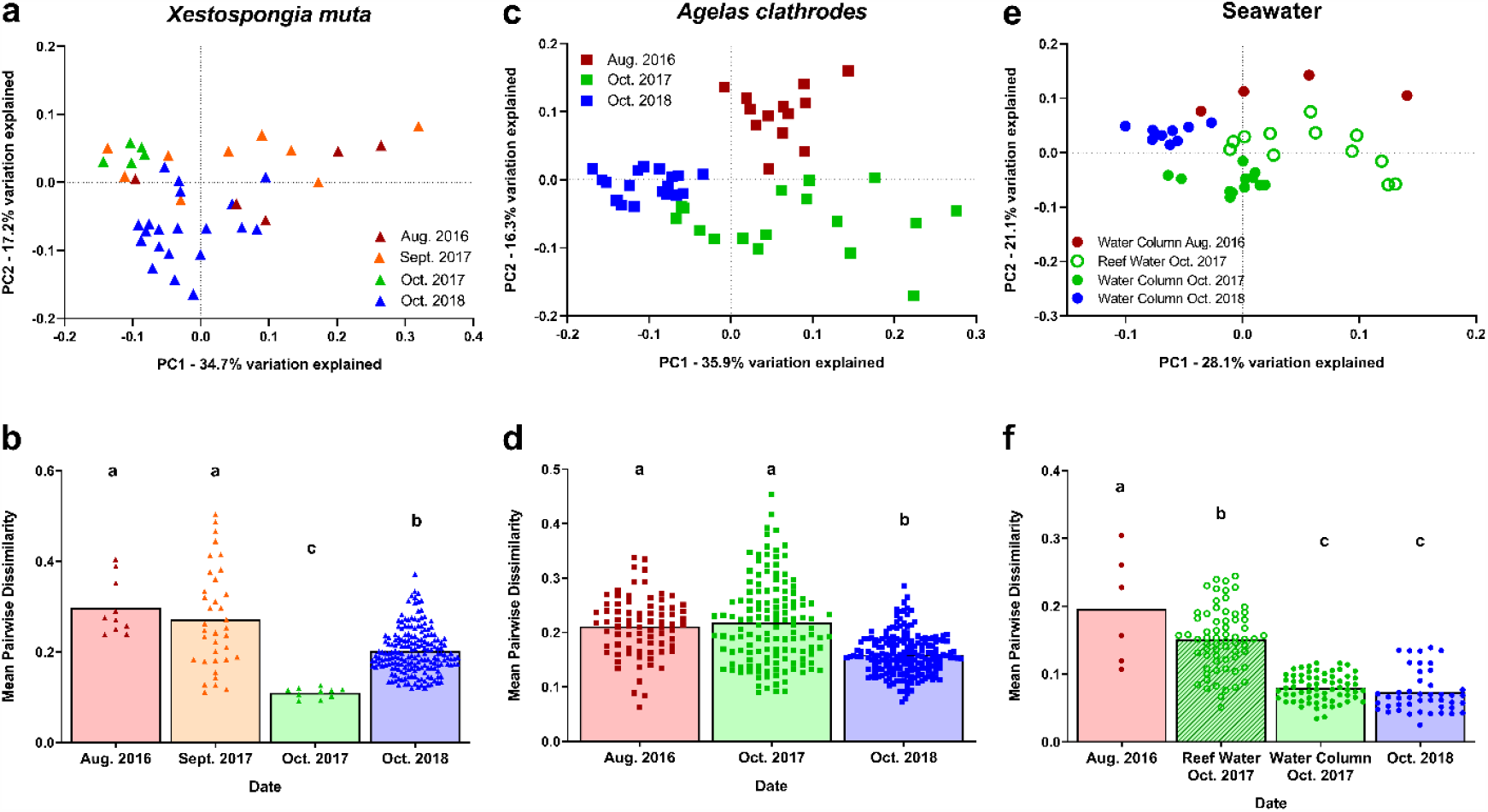
Sponge-associated bacterial communities differed in composition and variability following extreme storm-derived floods. Principle Coordinate Analysis of the weighted UniFrac distance matrices for (A) visually healthy *Xestospongia muta*, (B) visually healthy *Agelas clathrodes*, and (C) seawater bacterial communities. Mean (with individual value dots) pairwise dissimilarity values for (D) *X. muta*, (E) *A. clathrodes*, and (F) seawater bacterial communities. Red = August 2016. Orange = September 2017; Green = October 2017. Blue = October 2018 associated with no flooding stress. Bars represent mean (± sem). Within a species, bars that do not share a letter are significantly different based on ANOVA with Tukey’s multiple comparisons (p < 0.05).

Disruption of bacterial communities during both flood years included significant increases in community dispersion (or variation) as well as significant increases in community (or ASV) richness. Within *X. muta* samples, bacterial communities in Aug. 2016 and Sept. 2017 had significantly higher dispersion compared to the no-flood year, but the variability of bacterial communities in Oct. 2017 was lower than other flood-associated dates as well as the no-flood year (ANOVA with Tukey’s comparisons: p < 0.01, Figure 3d). In *X. muta*, there was a significant difference in ASV richness across Date (GLM: F = 6.00, p = 0.002) and Health State (GLM: F = 11.19, p = 0.002), but not by Site (GLM: F = 2.29 p = 0.139). Communities from Aug. 2016 and Sept. 2017 had higher ASV richness than communities from the no-flood year, and diseased *X. muta* communities had higher ASV richness than visually healthy *X. muta* communities. Bacterial communities in *A. clathrodes* also displayed higher dispersion during both flood years compared to the no-flood year (ANOVA: p < 0.05; Figure 3e). There was also a significant difference in ASV richness across Date (GLM: F = 9.49, p = 0.0001) and Health State (GLM: F = 43.14, p = 0.0001), but not by Site (GLM: F = 0.13 p = 0.724). Communities from both flood years had higher ASV richness than communities from the no-flood year, and affected *A. clathrodes* communities had higher ASV richness than visually healthy *A. clathrodes* communities. Seawater bacterial communities also displayed higher dispersion during both flood years compared to the no-flood year (ANOVA: p < 0.05; Figure 3f). However, in the 2017 flood year, Reef Water, but not Seawater Column samples had higher dispersion compared to the no-flood year. There was a significant difference in ASV richness across Date (GLM: F = 48.03, p = 0.0001) and Sample Type (GLM: F = 72.95, p = 0.0001), but not by Site (GLM: F = 1.32 p = 0.258). Communities from both flood years had higher ASV richness than the no-flood year. In *X. muta, A. clathrodes*, and seawater samples, Shannon Diversity was not impacted by Date, Health Status/Sample Type, or Site (GLM: p > 0.05).

Disruptions to bacterial communities of *X. muta, A. clathrodes*, and seawater samples during flood years are also reflected in significant changes in the relative abundance of several bacterial taxa (as assessed by DESeq2 analysis). *X. muta* bacterial communities during the no-flood year were dominated by Chloroflexi (21.4 ±1.2%), Gammaproteobacteria (14.3 ±1.0%), Acidobacteria (13.1 ±1.6%), and Actinobacteria (10.1 ±1.3%) (Supplemental Figure S1a). In *X. muta*, Flavobacteriaceae, Poribacteria, SAR86 clade, and Cyanobiaceae were enriched in both Aug. 2016 and Sept. 2017, but no bacterial Family was enriched or depleted across all three flood-associated dates (Supplemental Table S5). In *A. clathrodes*, bacterial communities during the no-flood year were also dominated by Chloroflexi (22.7 ±1.2%), Gammaproteobacteria (15.3 ±1.0%), Actinobacteria (15.1 ±1.6%), and Acidobacteria (13.6 ±0.6%) (Supplemental Figure S1b). Numerous (35) bacterial Families were enriched in Aug. 2016 as compared to the no-flood year; in contrast, only 5 Families were enriched in Oct. 2017 (Supplemental Table S5). In *A. clathrodes*, bacterial Families with potential human and marine pathogens (Enterobacteriaceae and Vibrionaceae) were enriched in Aug. 2016, and one Family (Stappiaceae) was enriched in both flooding years (Supplemental Table S5). For both sponge species, seven Families (Cyanobiaceae, Clostridiaceae, Desulfovibrionaceae, Halieaceae, Poribacteria, Vibrionaceae, and Vicinamibacteraceae) were enriched during flood-associated dates. For Aug. 2016, Desulfovibrionaceae was the most enriched bacterial Family in both sponge species. Seawater bacterial communities during the no-flood year were dominated by Alphaproteobacteria (36.9 ±0.9%), Cyanobacteria (31.2 ±2.4%), and Gammaproteobacteria (14.1 ±0.9%) (Supplemental Figure S1c). Many (95) diverse seawater-associated bacterial Families were enriched in each flood year, but only Burkholderiaceae, Pseudomonadaceae, SAR86 clade were enriched in both flood years (Supplemental Table S5). One bacterial Family (Halieaceae) was enriched during flood-associated dates in all three sample types.

Over 322 ASVs (198, 123, and 212 ASVs in *X. muta, A. clathrodes*, and seawater communities, respectively) were significantly associated with flooding events as detected by Indicator Species Analysis (Indicator Value > 0.3, p < 0.05; Supplemental Table S6). Eleven Indicator ASVs were detected in both sponge species in Aug. 2016, and one Indicator ASV was detected in both Aug. 2016 and Sept. 2017 (Table 2). No ASVs were significantly associated with Oct. 2017 in both sponge species. ASVs classified as *Halodesulfovibrio* or *Desulfovibrio* (sulfate-reducing taxa) were associated with the Aug. 2016 flooding event in both sponge species. Additionally, 5 ASVs classified as *Prochlorococcus* and *Synechococcus* (photosynthetic marine Cyanobacteria) were associated with flood years in both sponge species. An increase in one Indicator ASV, classified as *Synechococcus spongarium*, was the largest driver of differences in healthy *X. muta*-associated communities in Aug. 2016 and Sept. 2017 compared to the no-flood year (as assessed by Similarity Percentages (SIMPER) analysis). This *Synechococcus spongarium* ASV was not present in any *A. clathrodes* or seawater samples. Eight ASVs classified as *Prochlorococcus* and *Synechococcus* were also Indicator ASVs associated with flood years in seawater samples (Supplemental Table S6); however, none of these seawater-associated Cyanobacterial Indicator ASVs were present in any sponge samples. No Indicator ASV identified for seawater was also identified for a sponge species.

**Table 2.** Bacterial taxa unique to both *X. muta* and *A. clathrodes*-associated bacterial communities following recent flooding events* according to Indicator Species Analysis. Analysis was conducted on each sponge species individually.

| Date | Amplicon Sequence Variant ID | Taxonomy (Phylum_Class_Family_Genus) | Host | Indicator Value | P-value |
| --- | --- | --- | --- | --- | --- |
| Aug. 2016 | 64805be33440e427c18e31e0d5e6094b | Bacteroidota_Bacteroidia_Saprospiraceae_Phaeodactylibacter | *X. muta* | 0.577 | 0.028 |
|  |  |  | *A. clathrodes* | 0.379 | 0.004 |
|  | ccd8d5e31ad9bfa58066251ea1138225 | Bacteroidota_Bacteroidia_Flavobacteriaceae_NA | *X. muta* | 0.577 | 0.028 |
|  |  |  | *A. clathrodes* | 0.343 | 0.002 |
|  | e09f2d58d292332880fe9e37607469e1 | Desulfobacterota_Desulfovibronia_Desulfovibrionaceae_Halodesulfovibrio | *X. muta* | 0.573 | 0.028 |
|  |  |  | *A. clathrodes* | 0.385 | 0.000 |
|  | e4a5985be454d1e8a3d50a2ab6ea7a8b | Desulfobacterota_Desulfovibronia_Desulfovibrionaceae_Halodesulfovibrio | *X. muta* | 0.525 | 0.003 |
|  |  |  | *A. clathrodes* | 0.386 | 0.000 |
|  | e51947121c608cf15031596f9d0c09eb | Desulfobacterota_Desulfovibronia_Desulfovibrionaceae_NA | *X. muta* | 0.461 | 0.028 |
|  |  |  | *A. clathrodes* | 0.397 | 0.000 |
|  | 0261a50c7edb68abf5e65bb5094ce011 | Desulfobacterota_Desulfovibronia_Desulfovibrionaceae_NA | *X. muta* | 0.426 | 0.028 |
|  |  |  | *A. clathrodes* | 0.524 | 0.000 |
|  | 5546361caf3e42e50fb56d85a11571f2 | Proteobacteria_Alphaproteobacteria_Clade I_Clade Ia | *X. muta* | 0.533 | 0.025 |
|  |  |  | *A. clathrodes* | 0.511 | 0.001 |
|  | 2ef8937fcfef105310d9aac9f65c08ab | Proteobacteria_Gammaproteobacteria_Kangiellaceae_Aliikangiella | *X. muta* | 0.577 | 0.028 |
|  |  |  | *A. clathrodes* | 0.341 | 0.004 |
|  | b47b6e5813cfd4b7690b522531f10aa4 | Proteobacteria_Gammaproteobacteria_Thioglobaceae_SUP05 cluster | *X. muta* | 0.826 | 0.000 |
|  |  |  | *A. clathrodes* | 0.393 | 0.015 |
|  | efa28eb526ec0c3caa54fd35809774e1 | SAR324 clade_Marine group B_NA_NA | *X. muta* | 0.577 | 0.027 |
|  |  |  | *A. clathrodes* | 0.393 | 0.015 |
|  | 9442980515ce7f911bb4cc9c372db4e8<sup>#</sup> | Proteobacteria_Gammaproteobacteria_SAR86_clade_NA | *X. muta* | 0.805 | 0.000 |
|  |  |  | *A. clathrodes* | 0.669 | 0.001 |
| Aug. 2016 and Sept. 2017 | 4bf34e939f3102402abec9deab0319b3 | Proteobacteria_Gammaproteobacteria_Nitrosococcaceae_AqS1 | *X. muta* (Sept. 2017) | 0.489 | 0.036 |
|  |  |  | *A. clathrodes* (Aug. 2016) | 0.382 | 0.016 |

\*No ASVs associated with Oct. 2017 (1 month post-Hurricane Harvey) flooding were present in both *X. muta* and *A. clathrodes*-associated bacterial communities

### Sponge bacterial communities show signs of wastewater contamination after flooding

Thirty bacterial ASVs were classified as Family Enterobacteriaceae and were recovered from most samples of both sponge species. Of these 30 Enterobacteriaceae ASVs, 23 were further classified as *Escherichia coli*. In *X. muta*, bacterial communities had low abundances (< 0.1%) of Enterobacteriaceae (Figure 4a), displaying no significant differences across Date (GLM: H = 0.52, p = 0.674), Site (GLM: F = 1.01, p = 0.321), or Health Status (GLM: F = 0.59, p = 0.445). In contrast, *A. clathrodes* bacterial communities had a significantly higher abundance of Enterobacteriaceae in diseased samples compared to visually healthy samples (GLM: F = 66.11, p = 0.0001) and in Aug. 2016 samples compared to other dates (GLM: F = 5.24, p = 0.009). For example, diseased and visually healthy *A. clathrodes* samples in Aug. 2016 contained 7.91 (± 1.83)% and 1.44 (± 0.64)% Enterobacteriaceae, respectively, whereas in Oct. 2017 and the no-flood year, Enterobacteriaceae abundance was <0.1% (Figure 4b, Supplemental Figure S1d). No Enterobacteriaceae ASVs were detected in any seawater samples, and no Enterobacteriaceae ASVs were identified in Indicator Species Analysis as being significantly associated with either flood year (Supplemental Table S6).

**Figure 4.**
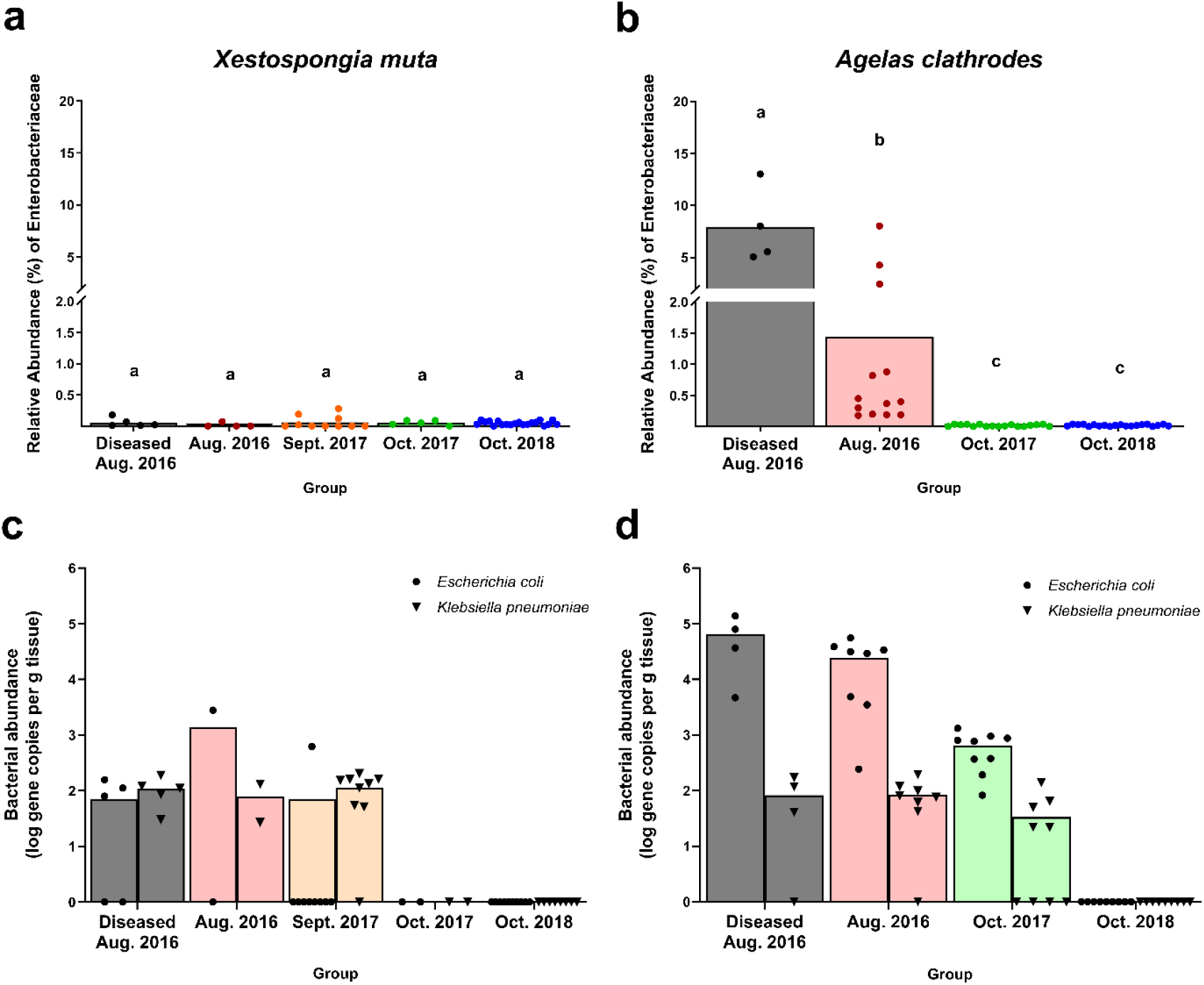
Relative abundance of sequences in the Family Enterobacteriaceae (A, B) and to specific human pathogens (*Escherichia coli* and *Klebsiella pneumoniae)* (C, D) across sites and years. Data in (A, B) are based on Illumina MiSeq amplicon sequencing of the V4 region of the bacterial 16S rRNA gene. Data in (C, D) are quantitative PCR amplification of the bacterial *ybbW* and *phoE* genes for *E. coli* and *K. pneumoniae*, respectively. All data are presented as mean with individual value dots. Black bars = affected sponges in 2016; red bars = healthy sponges in 2017; green bars = healthy sponges in 2017; blue bars = healthy sponges in 2018 (no flood, baseline year). Groups that share a letter (in a, b) are not significantly different based on a Generalized Linear Model with Tukey’s multiple comparisons across groups within a species. ND = not detected.

To test the hypothesis that FGB sponges were exposed to wastewater-derived bacteria from storm generated floodwaters, seawater and sponge samples were screened for seven human pathogens using quantitative PCR. Diseased and visually healthy sponge samples collected in both flood years yielded positive detection for 2 out of 7 human pathogens screened: *Escherichia coli* and *Klebsiella pneumoniae* (Figure 4c,d). No human pathogens were detected in sponges sampled during the no-flood year (Figure 4c,d). In *X. muta, E. coli* abundance was highest in visually healthy samples from Aug. 2016, with a mean of 1.96 ×10^3^ (± 1.40 ×10^3^) gene copies per g tissue, compared to a mean of 8.96 ×10^1^ (± 8.54 ×10^1^) and 6.90 ×10^1^ (± 2.18 ×10^1^) gene copies per g tissue for diseased samples from Aug. 2016 and visually healthy sponges from Sept. 2017, respectively (Figure 4c). However, *E. coli* was detected in only one *X. muta* sample in Sept. 2017. In *X. muta, K. pneumoniae* abundance was similar across groups, averaging 1.08 ×10^2^ (± 5.78 ×10^1^), 7.84 ×10^1^ (± 7.27 ×10^1^), and 1.06 ×10^2^ (± 6.74 ×10^1^) gene copies per g tissue for diseased samples from Aug. 2016, visually healthy sponges from Aug. 2016, and visually healthy sponges from Sept. 2017, respectively (Figure 5c). In *A. clathrodes, E. coli* was more abundant in samples from Aug. 2016, with means of 6.47 ×10^4^ (± 5.76 ×10^4^) and 2.85 ×10^4^ (± 1.79 ×10^4^) gene copies per g tissue for diseased and visually health sponges, respectively, compared to Oct. 2017, with a mean of 6.26 ×10^2^ (± 4.17 ×10^2^) gene copies per g tissue (Figure 4d). In *A. clathrodes, K. pneumoniae* was less abundant (> 2 orders of magnitude difference) compared to *E. coli*, displaying similar abundance across groups where it was detected, averaging 8.22 ×10^1^ (± 7.65 ×10^1^), 8.36 ×10^1^ (± 5.68 ×10^1^), and 4.61 ×10^1^ (± 3.29 ×10^1^) gene copies per g tissue for diseased samples from Aug. 2016, visually healthy sponges from Aug. 2016, and visually healthy sponges from Oct. 2017, respectively (Figure 4d). No seawater samples tested positive for *E. coli* or *K. pneumoniae*. No sponge samples tested positive for *Enterococcus* spp., *Pseudomonas aeruginosa, Salmonella enterica, Serratia marcescens*, and *Staphylococcus aureus* (data not shown).

## DISCUSSION

### Bacterial communities of offshore reef sponges are disrupted following major flooding events

It has been assumed that remote benthic marine ecosystems (>100 km from land) are not significantly affected by terrestrial runoff. Our findings, however, suggest that floodwaters can reach and impact offshore benthic reef organisms. Bacterial communities of both *X. muta* and *A. clathrodes* showed disruptions to their community structure following two flood events in 2016 and 2017, relative to sponges collected during the same season in a no-flood year in 2018. Furthermore, we quantified an increased relative abundance of Enterobacteriaceae and two known human pathogens (*E. coli* and *K. pneumoniae*) in post-flood sponge samples, indicating that bacteria of terrestrial origin may interact with offshore reefs following extreme storm events.

Bacterial communities associated with the two sponge species in this study exhibited some differences in the strength and duration of their response to flood water stress, likely because they harbored distinctive bacterial communities during the no-flood year. Bacterial communities associated with other sponge species have similarly exhibited variation in their resistance to some flood-associated stressors, such as elevated nutrients (Simister et al., 2012; Luter et al., 2014; Baquiran and Conaco, 2018) and/or reduced salinity (Glasl et al., 2018b). Larger shifts in *X. muta* bacterial communities were associated with the 2016 flood year. In contrast, *X. muta* bacterial communities were relatively resilient following the sub-lethal stress associated with the 2017 food year, with Oct. 2017 communities being more similar to the no-flood year. Interestingly, *A. clathrodes* sponges exhibited larger shifts in its microbiota in 2017 compared to 2016. Wright *et al*. (2019) similarly found that interspecific differences were the strongest driver of gene expression changes in two coral species following Hurricane Harvey in 2017. Differences in host resistance/resilience to environmental changes or differences in a hosts’ ability to regulate its bacterial community may explain why *X. muta* showed more ephemeral changes (and more dispersion) in response to the 2017 flood, whereas shifts in *A. clathrodes* bacterial communities lingered.

Bacterial community shifts documented in both sponge species after these floods could be driven by a combination of several mechanisms including: (a) invasion of floodwater-derived bacteria into host tissues, (b) invasion of seawater-derived bacteria into host tissues, and/or (c) shifts in the abundance of sponge-associated bacteria already present in hosts. It is plausible that all three mechanisms contributed, albeit to different degrees, following each flood event. Invasion of floodwater-derived bacteria into host tissues is supported by the presence of human pathogens in sponge-tissue in Aug. 2016. Invasion of seawater-derived bacteria into sponge tissue is supported by the enrichment of *Synechococcus* and *Prochlorococcus*, generally free-living marine Cyanobacteria, in both sponge species. However, sponge-associated and seawater associated bacterial communities were distinct across all dates, with few shared ASVs across all dates. Furthermore, relevant changes to *X. muta* bacterial communities in Aug. 2016 and Sept. 2017 were driven primarily by the relative increase of a resident sponge-associated Cyanobacterium (*Synechococcus spongarium*), which was not present in any seawater samples. This suggests that shifts in the abundance of bacterial taxa already present in sponges drove much of the bacterial community shift documented after these floods. To confirm the mechanism(s) underlying such microbial shifts on reefs, robust manipulative experiments or ‘before flood’ environmental samples of seawater and benthic host tissues would be required.

Changes in environmental parameters during and after flood events potentially drive shifts in the structure and composition of sponge- and seawater-associated bacterial communities by promoting or inhibiting growth of particular bacterial taxa and/or reducing the capacity of the animal host to regulate its microbiome (Zaneveld et al., 2017; Pita et al., 2018). For example, floodwaters over the FGB in 2016 contained higher ammonium concentrations (Kealoha et al., 2020), which may be contributing to enrichment of *Synechococcus* which generally occurs in higher abundance in the more mesotrophic environments near coasts (Partensky et al., 1999). Surface salinity decreased at the FGB following both flood events, but salinity at depth (24 m), reported from instruments at the WB, was unchanged preceding and during the flood event in 2016 (Johnston et al., 2019), suggesting that changes in salinity did not directly impact sponge-associated communities. Measurements for other environmental parameters associated with terrestrial runoff, such as turbidity, nutrient content, and pH, are not available for waters at the collection sites within FGB following these flood events.

### Wastewater contamination of sponges after severe flooding reaches offshore marine ecosystems

The increased abundance of Enterobacteriaceae in reef sponges 185 km offshore in the 2016 flood year, and, particularly, the detection of two fecal coliforms (*E. coli* and *K. pneumoniae*) in both flood years strongly suggest that FGB reefs were exposed to wastewater contamination after severe storms. Although the Family Enterobacteriaceae is ubiquitous in nature, occupying terrestrial, aquatic, and marine habitats, especially the intestine of homeothermic animals (Whitman, 2015), members of this group are not common or abundant members of the bacterial communities of sponges (Moitinho-Silva et al., 2017; Cleary et al., 2019). Other sources of Enterobacteriaceae, such as excreted waste from reef fish and marine mammals (Wallace et al., 2013) could explain the presence of this bacterial family across flood year sponge samples. Although the GoM has high ship traffic, it is unlikely that ship wastewater discharge represents a significant source of fecal coliforms to these reef organisms because it is prohibited for ships to discharge untreated human waste within the boundaries of the FGB National Marine Monument (https://flowergarden.noaa.gov/about/regulations.html). If any of these other potential sources drove the Enterobacteriaceae patterns in this study, then similar detections would be expected in all years. Other members of the Enterobacteriaceae, besides those screened for via qPCR, do not explain the presence of Enterobacteriaceae reads in samples across all sampling dates because most Enterobacteriaceae ASVs were classified as *E. coli*. Given that no human pathogens were detectable from samples during the no-flood year, the most parsimonious explanation for the Enterobacteriaceae detections in this study is that FGB reefs were exposed to land-based human wastewater via terrestrial runoff following the 2016 and 2017 floods.

We expected to find Enterobacteriaceae in seawater samples during one or both flood years; however, no members of this Family were observed in sequencing data or detected via qPCR. The dynamic, transient nature of seawater masses and the ability of sponges to filter feed, and thus, concentrate bacterial cells, may explain why Enterobacteriaceae was observed in sponge samples but not seawater samples. Fine-scale time series sampling of the seawater above FGB in the weeks following each flood event could have further resolved potential interactions among bacterial communities in near-reef seawater and benthic reef organisms. Unfortunately, logistical challenges stemming from the offshore location of the FGB precluded the collection of these samples. Although *X. muta* is larger and thus likely has a higher filtration rate (McMurray et al., 2008; Parra-velandia et al., 2014; Morganti et al., 2019), *A. clathrodes* had a higher frequency of detection of *E. coli* and contained this bacterial taxa at higher abundances. Interspecific differences in the digestion of filtered material (Lynch and Phlips, 2000) may have contributed to differences in the abundance of human pathogens between these two sponge sponges. Regardless, this work demonstrates that sponge species can be effective tools for monitoring wastewater contamination in offshore, as well as nearshore, marine ecosystems, and that the species selected for monitoring requires consideration.

It is unclear whether wastewater-derived bacteria contributed to mortality at EB in July of 2016, but detection of wastewater contamination at FGB (especially in diseased *A. clathrodes* samples) raises the question: do fecal coliforms pose health risks to offshore reefs? Human or animal wastewater contamination is linked to negative impacts on coral communities, particularly based on the input of excess nutrients, but chemical, bacterial, and pharmaceutical contamination are also potential issues (Wear and Thurber, 2015). The wastewater-derived bacteria, *Serratia marcescens*, is a coral pathogen (Sutherland et al., 2010, 2011), and in Hawaii, fecal coliforms colonized coral tissues after exposure, potentially contributing to a major disease outbreak (Beurmann et al., 2018). There is little information on the effect of fecal coliform exposure on marine sponge health, but some sponges may be relatively tolerant, using bacterial cells as a source of nutrition (Chaves-Fonnegra et al., 2007). The surprisingly far reach of contaminated floodwaters observed here underscores the urgent need to understand how floodwaters impact the health and function of offshore reef environments. A key question to address is whether detection of *E. coli* and *K. pneumoniae* represent detection of DNA from dead wastewater-derived bacterial cells or whether living wastewater-derived bacteria are potentially interacting with sponges (and other marine life) following extreme storms. If wastewater-derived bacteria contaminating sponges are metabolically active, then we must determine how long they persist within the sponge microbiome and the extent to which these microbes impact sponge physiology and function. A better understanding of the interactions between benthic reef hosts and terrestrial-derived bacteria will support effective management and protection of offshore reef ecosystems, such as the FGB.

### Comparisons of FGB sponge bacterial communities to those in the wider Caribbean

This study is the first to characterize sponge-associated microbial communities from the northwest Gulf of Mexico (and the FGB), offering the opportunity for comparisons of sponge microbial communities across regions of the GoM and the Caribbean. *X. muta*-associated bacterial communities at the FGB in 2018 (no storm condition) were similar between EB and WB and were dominated by Phyla also commonly reported from *X. muta* in the Florida Keys, Bahamas, and greater Caribbean (i.e. Proteobacteria, Chloroflexi, Cyanobacteria, Poribacteria, and to a lesser extent, Acidobacteria, Actinobacteria, and Crenarchaeota (Schmitt et al., 2012; Olson and Gao, 2013; Montalvo et al., 2014; Fiore et al., 2015; Morrow et al., 2016; Villegas-Plazas et al., 2019). These previous studies report regional differences due to changes in relative abundance of these shared Phyla (Fiore et al., 2015; Morrow et al., 2016). *X. muta* bacterial communities from the FGB may also be regionally distinct, in particular containing a high abundance of Crenarchaeota archaea (∼10%) compared to what has been reported from other regions (<5%) (Fiore et al., 2013). Ammonia oxidizing Archaea, such as Nitrosomopumilaceae, play an important role in nitrogen cycling in *X. muta* (López-Legentil et al., 2010). Ammonia-oxidizing archaea are outcompeted in environments with higher levels of ammonium (Erguder et al., 2009), so the greater abundance of Nitrosomopumilaceae likely reflects the oligotrophic environment of the offshore FGB reefs during no storm conditions.

Bacterial communities of *A. clathrodes* at the FGB contained Phyla (i.e. Proteobacteria, Firmicutes, Actinobacteria, Chloroflexi, Crenarchaeota) also present in other *Agelas spp*. from the Florida Keys, Belize, and Central Amazon Shelf (Olson and Gao, 2013; Deignan et al., 2018; Rua et al., 2018; Gantt et al., 2019). However, higher abundances of Archaea, especially Euryarchaeota and Crenarchaeota, and Firmicutes were found in other *Agelas* spp. (Deignan et al., 2018; Rua et al., 2018). Diseased sponges (both *A. clathrodes* and *X. mut*a) sampled after the 2016 mortality event were dominated by Alphaproteobacteria, especially Rhodobacteraceae. Alphaproteobacteria were also enriched in sponges affected by *Agelas* Wasting Syndrome (Deignan et al., 2018), suggesting that this group of bacteria could play a role in pathogenesis and/or serve as a biomarker of disease risk for FGB sponge communities.

### Mitigating the Impacts of Future Storms on Offshore Reefs

This study demonstrates that floodwaters following extreme storms can reach the vicinity of offshore reefs and may contribute to shifts and increased heterogeneity in the bacterial communities of sponges and other benthic reef organisms. Detection of bacteria typically associated with wastewater within these sponge samples illustrates that marine-terrestrial interactions, and thus, the potential impacts of human land and waste management practices, extend far beyond the shoreline. The GoM is regularly impacted by hurricanes, and thus marine communities in the region have evolved in a disturbance regime that includes bursts of storm-generated terrestrial runoff. However, the ongoing expansion of impermeable surfaces (e.g., concrete, pavement) that reduce water absorption by soil in Texas and other coastal areas, as well as changing extreme storm patterns (e.g., slower moving hurricanes with greater precipitation) are increasing the frequency and intensity of floodwater influx into marine environments.

This study of the potential impacts of the 2016 and 2017 floods was catalyzed because a mortality event affected the East Bank following the Tax Day Flood. We hypothesize that flood waters associated with other recent extreme storm events (e.g., 2015 Memorial Day Flood, flooding generated by Hurricane Imelda in September 2019) in the region likely also caused sub-lethal stress at the FGB. However, targeted sampling of FGB did not occur following these storms. Our findings clearly demonstrate the urgent need for: (1) continued mitigation of stormwater runoff and climate change impacts; and (2) establishment of surface and benthic microbial and water quality time series for near- and offshore reefs using standardized protocols. This latter program will ideally generate baseline data on the gene expression and microbiomes of key benthic reef taxa under normal conditions, providing critical context (Glasl et al., 2018a) in which to detect and mitigate floodwater-derived stress on reefs in order to understand their impact on benthic invertebrate physiology and reef ecosystem functions.

## MATERIALS AND METHODS

### Pelagic water properties during sample collection periods

The Flower Garden Banks National Marine Sanctuary (northwest Gulf of Mexico) is comprised of three banks: East Bank (EB), Stetson Bank and West Bank (WB) (Figure 1a). Surface salinity data, available from the Texas Automated Buoy System (TABS) Real Time Ocean Observations system archive (http://tabs.gerg.tamu.edu) were used to assess the potential of flood waters to reach the vicinity of the FGB reef surface waters following recent severe storm events. To characterize local surface salinity (sensor depth of 2 m) in parts per thousand (ppt) representative for the FGB before, during, and after each sampling period, water property data collected each half hour for April-October for the years 2006-2018 were downloaded for TABS Buoy V (27°53.7960’N, 93°35.8380’W), which is located approximately 3 km from EB and 25 km from WB. Data were filtered to remove all timepoints containing no data and to exclude outlier data (i.e., measurements where salinity abruptly changed to <1 ppt or >70 ppt from 35 ppt). Data were not available (due to instrumentation failure) in 2007 and between the dates of 6 June 2017 through 30 August 2017. The remaining data were then plotted with ggplot2 package version 3.3.2 using code from https://github.com/rachelwright8/TagSeq_FGB_HurricaneHarvey. Data were summarized in Figure 1d as means (black line) and daily ranges (grey shading) for April to October over the thirteen-year period. To assess the lags between continental flooding (associated with an extreme storm) and changes in water quality at the FGB, daily salinity means and range data for April to October of individual years 2016, 2017 and 2018 (red lines and shading, respectively) were overlaid on surface salinity averages for the thirteen-year summary data. Daily means for each sponge sampling date were also calculated and plotted on top of the thirteen-year summary data with colored icons (Figure 1d). For sampling campaigns that spanned more than one day, only the first day of sampling was plotted (the maximum length of a sampling campaign was 5 days).

### Sample Collections

Sponge samples were collected at four timepoints spanning 2016-2018 from two locations within the FGB; Buoy 4 at East Bank (27°52’54.84”, 93°37’41.84”) and Buoy 2 at West Bank (27°54’28.8”, 93°36’0.72”, Figure 1a, Table 1). At all sampling timepoints, fragments were collected from individual *A. clathrodes* and *X. muta* sponges. The same individual sponges were not repeatedly sampled across timepoints due to time constraints in available ship and dive time. In August 2016, samples were collected from ‘diseased sponges’ that were exhibiting progressive tissue loss, as well as from visually healthy sponges. Diseased sponges were sampled at the interface between lesion ad healthy tissue. Representative photos of diseased sponges are presented in Johnston *et al*., (2018). For all other timepoints, samples were collected from visually healthy sponges as diseased sponges were not observed. Samples were clipped from sponge individuals using health status and species-specific cutting shears that were wiped clean between samples. Each sample was placed in an individual sterile bag for transport to the surface. Once topside, each sample was immediately transferred to liquid nitrogen and stored at-20°C until further processing. In total, 109 sponge samples, collected from depths ranging 18-27 m, were analyzed in this study (individual sample metadata provided in Supplemental Table S1).

Seawater samples were collected at three locations over EB (Buoy 4, Grid site 43 and Grid Site 42) and 2 locations over WB (Buoy 2 and Grid Site 23) (Figure 1a, Supplemental Table S1). ‘Grid’ sites refer to sampling sites along a 10 x 10 km grid over the FGB which was established in response to the localized mortality event detected in July 2016 and is described further in Kealoha *et al*., 2020. In Aug. 2016, Oct. 2017, and Oct. 2018, seawater samples were collected from surface waters (< 2m depth) as well as within the water column (10-30 m depth) using a Niskin bottle rosette on a Seabird Electronic (SBE) 25 CTD profiler. Additionally, in Oct. 2017, seawater samples were collected directly above the reef benthos at Buoys 4 and 2 by divers. All CTD collected water samples were sieved with a 100-um nylon net to exclude zooplankton. For each sample, 1000 mL of sieved seawater was then vacuum filtered (≤ 20 cm Hg) through a 47 mm, 0.22 µm Supor PES filter membrane (Pall) and immediately stored at-20°C. After returning to port, samples were transported on dry ice to Texas A&M University and stored at-80°C until DNA extraction.

### Bacterial Community Analysis

DNA was extracted from 250 mg of sponge sample using the Nucleospin Soil DNA extraction kit (Takara Bio) and was submitted to the Baylor College of Medicine’s Alkek Center for Metagenomics & Microbiome Research for high-throughput sequencing. High-throughput sequencing of the V4 hypervariable region of the 16S gene was conducted with 515f: 5’ GTGYCAGCMGCCGCGGTAA 3’ and 806rb: 5’ GGACTACNVGGGTWTCTAAT 3’ primers (Apprill et al., 2015) using the Illumina Mi-Seq (Paired-end 2x 250 read) platform. DNA extraction blanks were not performed during extraction of the sponge samples; however, work on these samples was conducted in a lab in which no human wastewater, nor sponge (or reef) samples had ever been processed previously. Total DNA from seawater samples was extracted from filters using FastDNA Spin kits (MP Biomedical) with a BioSpec Mini-Beadbeater-24. Each sample was amplified in triplicate 25 μL reactions with the following cycling parameters: 95°C for 3min, 30 cycles of 95°C for 45 s, 50°C for 60 s, and 72°C for 90 s, and a final elongation step at 72°C for 10min. All amplifications were performed using the 515F-806R primer pair modified with barcodes and Illumina MiSeq adapters. DNA extraction blanks were performed on seawater samples and did not yield any amplification products. Following amplification, the triplicate products were combined together and run on a 1.5% agarose gel to assess amplification success and relative band intensity. Amplicons were then quantified with the QuantiFluor dsDNA System (Promega), pooled at equimolar concentrations, and purified with an E.Z.N.A. Cycle Pure PCR Clean-Up Kit (Omega Bio-Tek). The purified library was sequenced on the Illumina MiSeq platform (v2 chemistry, 2×250 bp) at the Georgia Genomics Facility (Athens, GA, USA).

Sequence analysis was conducted using QIIME2 v. 2019.10 pipeline (Bolyen et al., 2019). Pair-end, demultiplexed reads were quality filtered, trimmed of poor-quality bases, de-replicated, chimera filtered, pair merged, and identified as amplicon sequence variants (ASVs) using the DADA2 plug-in (Callahan et al., 2016). Taxonomy was assigned by training a naïve-Bayes classifier on the V4 region of the 16S gene in the SILVA version 138 database (Quast et al., 2013) using the feature-classifier plugin (Bokulich et al., 2018). A 90% confidence threshold was used to retain classifications at each level of taxonomy. Low abundance ASVs (< 2 occurrences over all samples) and non-prokaryotic ASVs (i.e., mitochondria, chloroplast, eukaryote, and unknown sequences) were removed. Rarefied ASV tables (rarefied to 9,000 reads per sample) were used to calculate alpha diversity metrics and to conduct beta diversity analyses using weighted UniFrac distance matrices.

### Quantitative PCR for human pathogens associated with Hurricane Harvey-derived floodwaters

Species-specific functional genes were chosen as biomarkers to detect and quantify fecal indicator bacteria (*Escherichia coli* and *Enterococcus spp*.), putative pathogenic bacteria in the Family Enterobacteriaceae (*Klebsiella pneumoniae, Serratia marcescens*, and *Salmonella enterica*), and other putative pathogenic bacteria (*Pseudomonas aeruginosa* and *Staphylococcus aureus*, Supplemental Table S7). These bacterial species were targeted because they were identified in qPCR screening and in high-throughput sequencing of terrestrial floodwater and sediment samples collected immediately following Hurricane Harvey (Yu et al., 2018). Sponge samples were screened for all seven bacterial species. Seawater samples were then screened for the bacterial species found in sponges (*E. coli* and *K. pneumoniae*).

Target gene amplicons were used to establish the standard curve between the threshold cycle (Ct) value and log10 (gene copies) for each pathogenic bacterium individually. To generate amplicons for target gene quantitation, genomic DNA of pure cultures of each bacterial strain was extracted using DNeasy PowerSoil Kit (Qiagen) and target genes were amplified by conventional PCR (50 µL reactions) with EmeraldAmp GT PCR Master Mix (Takara Bio, thermocycler conditions listed in Supplemental Table S7). Five µL of each PCR product was visualized via gel electrophoresis to confirm amplification, and the remaining PCR product was cleaned using GeneJET PCR Purification Kit (ThermoScientific). Amplicon concentration was quantified using a Qubit 3 Fluorometer (Invitrogen) with dsDNA Broad Range Assay (Invitrogen), amplicon quality was assessed using a NanoDrop One (Thermo Scientific), and amplicon sequences were confirmed via Sanger sequencing. Verified amplicons were then serially diluted to create a set of standards of known concentrations, calculated by the Thermo Fisher DNA Copy Number Calculator (https://www.thermofisher.com/us/en/home/brands/thermo-scientific/molecular-biology/molecular-biology-learning-center/molecular-biology-resource-library/thermo-scientific-web-tools/dna-copy-number-calculator.html). Each standard curve was established by linear regression of the threshold cycle (C_T_) values versus the log-normal (10 base) abundance of gene copies (10^8^, 10^7^, 10^6^, 10^5^, 10^4^, 10^3^, 10^2^) from technical triplicates. R^2^ values of least square fit ranged from 0.995-0.999 for all the standard curves. The qPCR amplification efficiency for all the biomarkers ranged from 92-105%, calculated by plotting average C_T_ vs amplicon concentration (copies/µL) on a log_10_ scale. To assess gene copy number in a given sample, C_T_ values of samples were compared to a prepared standard curve that was included in triplicate with each qPCR run. Calculated copy number was normalized to g of wet sponge tissue and normalized to input DNA concentration of each sample as conducted previously (Radax et al., 2012; Moeller et al., 2019). The limit of quantification ranged from 20-100 gene copies.

Quantitative PCR reaction mix consisted of 5 µL 2x Power SYBR Green Master Mix (Applied Biosystems), 1 µL DNA, 1.3 µL of each primer (10 µmol stock), and molecular-grade water, for a final volume of 10 µL. Primer specifications and thermocycler conditions for each pathogen are listed in Supplemental Table S2. All samples were run in triplicate on a QuantStudio 3 Real-Time PCR System (ThermoScientific) and were screened for all seven pathogens (Supplemental Table S7). Negative controls (no template) were run in triplicate for each qPCR experiment to monitor for potential contamination. The temperature profile for SYBR Green qPCR involved 95°C for 10 min, 45 cycles of 95°C for 15 sec and annealing/extension temperature of 54 – 65°C for 1 min. Melt curve analysis was conducted after PCR completion to ensure nonspecific PCR products were not generated. Specificity of each qPCR assay was confirmed by testing for amplification in all pathogen strains used in this study as well as in four environmental isolates (*Vibrio* sp., *Alteromonas* sp., *Pseudoalteromonas* sp., and *Shewanella* sp.) previously cultured from FGB coral. No non-target strains amplified below a threshold cycle (C_T_) of 30. Amplifications were considered positive if all three replicates amplified at a threshold cycle (C_T_) less than 28 and melt curve analysis showed similar patterns to the standard.

### Statistics

A weighted UniFrac distance matrix was used to calculate beta-diversity and to assess within group dispersion in bacterial communities and to construct Principle Coordinates Analysis (PCoA) plots to visualize differences in bacterial community structure between groups using QIIME2. PCoA was conducted for all samples (both sponge species), as well as for healthy samples of each species individually. Pairwise Analysis of Similarities (ANOSIM), conducted with 999 permutations, was used to test for significant differences in bacterial communities among categorical variables including species, health state, site, and collection date. To assess differences in bacterial abundance at the Family level, the unrarefied ASV table for each species was first summarized to Family level using tax_glom in phyloseq (v1.30.0). A negative binomial model was then fitted with the R package DESeq2 (v1.26.0) and Wald tests were used to test for differences in taxon abundance within each comparison of Date versus the ‘baseline’ Date (October 2018). Benjamini-Hochberg FDR tests were used to account for multiple comparisons, and Families with p-values less than 0.05 were identified as significantly differentially abundant. Indicator species analysis was performed to test the association between bacterial community and collection date, using indicspecies (v1.7.9) package in R. Significance was assessed with 9999 permutations, and ASVs with p-values less than 0.05 selected. PRIMER-E v6 (Primer-E Ltd) was used to conduct Similarity Percentages (SIMPER) analysis, which identified ASVs that contributed to differences between bacterial communities across Dates. Differences in ASV richness, Shannon Diversity, relative abundance of bacterial taxa, and quantified bacterial abundance data were analyzed within the three sample types (*X. muta, A. clathrodes*, and Seawater) individually using generalized linear models (GLM) with Site, Health State (healthy vs. affected sponge), Seawater sample type (water column vs. reef water), and Date as categorical predictors. If significant differences were detected, then a Tukey’s HSD post hoc test was performed to examine the differences between the levels of the independent variables. Differences in mean community variability (mean pairwise dissimilarity) across Dates was assessed using ANOVA with Tukey’s pairwise comparisons. All data are represented as mean (± SEM), unless otherwise stated.

## DATA AVAILABILITY

The raw sequence data files are available in the NCBI Sequence Read Archive under accession numbers: SRP248232 for sponge samples, PRJNA509639 for Aug. 2016 seawater samples and PRJNA691373 for Oct. 2017 and Oct. 2018 seawater samples. Data files including Sample Metadata, ASV Table, ASV Taxonomy Assignment, and R script for DESeq and Indicator Species analysis are available on FigShare at https://figshare.com/account/home#/projects/82841.

## AUTHOR CONRTRIBUTIONS

AMSC, JS, LSV, LS, KS, and SD obtained financial support and conceived experimental design, sampling strategy, site selection, sampling methodologies for fieldwork. All authors contributed to dive support, sample collection, and/or sample processing. AS, CGBG, LIHK, and JAS contributed to data analysis. AS, LIHK, JAS, LS, and AMSC contributed to manuscript preparation including figures and tables. All authors reviewed manuscript drafts.

### FUNDING

This work was funded by NSF awards OCE-1800914 to AC, OCE-1800905 to LSV, OCE-1800904 to SD, OCE-1800913 to KEFS and JBS, as well as a Rice University Faculty Initiative award to AC and LS, and a NAS Gulf Research Program Early Career Fellowship and Rice University start-up funds to AC. AS was supported by a Rice Academy Postdoctoral Fellowship and Rice University start-up funds (to AC).

## ACKNOWLEDGEMENTS

We thank Marissa Nuttall, John Embesi and James MacMillan (NOAA), Jake Emmert and Kaitlin Buhler (Moody Gardens), and Ryan Eckert (Florida Atlantic University) for field and logistical support. We are grateful to Emma Hickerson for facilitating collection permits. We also acknowledge the following members of the science party during sampling cruises: Jennifer Drummond, Kelsey Sanders, and Anna Knochel (Rice University). We also acknowledge Rachel Wright for the code used to retrieve and plot TABS Buoy V data. 2016 samples were collected under FGBNMS-2014-001; 2017 and 2018 samples were collected under FGBNMS-2017-012 to AC. August 2016 and September 2017 samples were collected on the R/V Manta, whereas October 2017 and October 2018 samples were collected aboard the R/V Point Sur. We express our sincere gratitude to the crews of these vessels, as well as members of the NOAA FGBNMS office and Moody Gardens Aquarium (Galveston, TX). This manuscript was released as a pre-print at BioRxiv, under DOI: 10.1101/2020.04.27.064568v3 (Shore et al., 2020).

## REFERENCES

Apprill, A., Mcnally, S., Parsons, R., and Weber, L. (2015). Minor revision to V4 region SSU rRNA 806R gene primer greatly increases detection of SAR11 bacterioplankton. Aquat. Microb. Ecol. 75, 129–137. doi:10.3354/ame01753.

Baquiran, J. I. P., and Conaco, C. (2018). Sponge-microbe partnerships are stable under eutrophication pressure from mariculture. Mar. Pollut. Bull. 136, 125–134. doi:10.1016/j.marpolbul.2018.09.011.

Bell, J. J., Davy, S. K., Jones, T., Taylor, M. W., and Webster, N. S. (2013). Could some coral reefs become sponge reefs as our climate changes? Glob. Chang. Biol. 19, 2613–2624. doi:10.1111/gcb.12212.

Beurmann, S., Ushijima, B., Videau, P., Svoboda, C. M., Chatterjee, A., Aeby, G. S., et al. (2018). Dynamics of acute montipora white syndrome: Bacterial communities of healthy and diseased M. capitata colonies during and after a disease outbreak. Microbiol. (United Kingdom) 164, 1240–1253. doi:10.1099/mic.0.000699.

Blake, E. S., and Zelinsky, D. A. (2018). National Hurricane Center Tropical Cyclone Report Hurricane Harvey.

Bokulich, N. A., Kaehler, B. D., Rideout, J. R., Dillon, M., Bolyen, E., Knight, R., et al. (2018). Optimizing taxonomic classification of marker-gene amplicon sequences with QIIME 2’s q2-feature-classifier plugin. Microbiome 6, 1–17. doi:10.1186/s40168-018-0470-z.

Bolyen, E., Rideout, J. R., Dillon, M. R., Bokulich, N. A., Abnet, C. C., Al-Ghalith, G. A., et al. (2019). Reproducible, interactive, scalable and extensible microbiome data science using QIIME 2. Nat. Biotechnol. 37, 852–857. doi:10.1038/s41587-019-0209-9.

Callahan, B. J., McMurdie, P. J., Rosen, M. J., Han, A. W., Johnson, A. J. A., and Holmes, S. P. (2016). DADA2: High-resolution sample inference from Illumina amplicon data. Nat. Methods 13, 581–583. doi:10.1038/nmeth.3869.

Chaves-Fonnegra, A., Zea, S., and Gómez, M. L. (2007). Abundance of the excavating sponge Cliona delitrix in relation to sewage discharge at San Andrés Island, SW Caribbean, Colombia. Bol. Investig. Mar. y Costeras 36, 63–78. doi:10.25268/bimc.invemar.2007.36.0.201.

Chen, S., Lu, Y. H., Dash, P., Das, P., Li, J., Capps, K., et al. (2019). Hurricane pulses: Small watershed exports of dissolved nutrients and organic matter during large storms in the Southeastern USA. Sci. Total Environ. 689, 232–244. doi:10.1016/j.scitotenv.2019.06.351.

Cleary, D. F. R., Swierts, T., Coelho, F. J. R. C., Polónia, A. R. M., Huang, Y. M., Ferreira, M. R. S., et al. (2019). The sponge microbiome within the greater coral reef microbial metacommunity. Nat. Commun. 10, 1–12. doi:10.1038/s41467-019-09537-8.

Congdon, V. M., Bonsell, C., Cuddy, M. R., and Dunton, K. H. (2019). In the wake of a major hurricane: Differential effects on early vs. late successional seagrass species. Limnol. Oceanogr. Lett. 4, 155–163. doi:10.1002/lol2.10112.

Deignan, L. K., Pawlik, J. R., and Erwin, P. M. (2018). Agelas Wasting Syndrome Alters Prokaryotic Symbiont Communities of the Caribbean Brown Tube Sponge, Agelas tubulata. Microb. Ecol. 76, 459–466. doi:10.1007/s00248-017-1135-3.

Deslarzes, K., and Lugo-Fernandez, A. (2007). “Influence of Terrigenous Runoff on Offshore Coral Reefs: An Example from the Flower Garden Banks, Gulf of Mexico,” in Geological Approaches to Coral Reef Ecology, ed. R. B. Aronson (New York, NY: Springer), 126–160. doi:https://doi.org/10.1007/978-0-387-33537-7_5.

Dodge, R. E., and Lang, J. C. (1983). Environmental correlates of hermatypic coral (Montastrea annularis) growth on the East Flower Gardens Bank, northwest Gulf of Mexico. Limnol. Oceanogr. 28, 228–240. doi:10.4319/lo.1983.28.2.0228.

Emanuel, K. (2017). Assessing the present and future probability of Hurricane Harvey’s rainfall. Proc. Natl. Acad. Sci. U. S. A. 114, 12681–12684. doi:10.1073/pnas.1716222114.

Erguder, T. H., Boon, N., Wittebolle, L., Marzorati, M., and Verstraete, W. (2009). Environmental factors shaping the ecological niches of ammonia-oxidizing archaea. FEMS Microbiol. Rev. 33, 855–869. doi:10.1111/j.1574-6976.2009.00179.x.

Fabricius, K. E. (2005). Effects of terrestrial runoff on the ecology of corals and coral reefs: Review and synthesis. Mar. Pollut. Bull. 50, 125–146. doi:10.1016/j.marpolbul.2004.11.028.

Fiore, C. L., Jarett, J. K., and Lesser, M. P. (2013). Symbiotic prokaryotic communities from different populations of the giant barrel sponge, Xestospongia muta. Microbiologyopen 2, 938–952. doi:10.1002/mbo3.135.

Fiore, C. L., Labrie, M., Jarett, J. K., and Lesser, M. P. (2015). Transcriptional activity of the giant barrel sponge, Xestospongia muta Holobiont: Molecular evidence for metabolic interchange. Front. Microbiol. 6. doi:10.3389/fmicb.2015.00364.

Gantt, S. E., McMurray, S. E., Stubler, A. D., Finelli, C. M., Pawlik, J. R., and Erwin, P. M. (2019). Testing the relationship between microbiome composition and flux of carbon and nutrients in Caribbean coral reef sponges. Microbiome 7, 1–13. doi:10.1186/s40168-019-0739-x.

Glasl, B., Bourne, D. G., Frade, P. R., and Webster, N. S. (2018a). Establishing microbial baselines to identify indicators of coral reef health. Microbiol. Aust. 39, 42–46. doi:10.1071/MA18011.

Glasl, B., Smith, C. E., Bourne, D. G., and Webster, N. S. (2018b). Exploring the diversity-stability paradigm using sponge microbial communities. Sci. Rep. 8, 1–9. doi:10.1038/s41598-018-26641-9.

Humphrey, C., Iverson, G., Skibiel, C., Sanderford, C., and Blackmon, J. (2019). Geochemistry of flood waters from the tar river, North Carolina associated with Hurricane Matthew. Resources 8, 1–12. doi:10.3390/resources8010048.

Humphrey, C., Weber, M., Lott, C., Cooper, T., and Fabricius, K. (2008). Effects of suspended sediments, dissolved inorganic nutrients and salinity on fertilisation and embryo development in the coral Acropora millepora (Ehrenberg, 1834). Coral Reefs 27, 837–850. doi:10.1007/s00338-008-0408-1.

Johnston, M. A., Embesi, J. A., Eckert, R. J., Nuttall, M. F., Hickerson, E. L., and Schmahl, G. P. (2016). Persistence of coral assemblages at East and West Flower Garden Banks, Gulf of Mexico. Coral Reefs 35, 821–826. doi:10.1007/s00338-016-1452-x.

Johnston, M. A., Nuttall, M. F., Eckert, R. J., Blakeway, R. D., Sterne, T. K., Hickerson, E. L., et al. (2019). Localized coral reef mortality event at East Flower Garden Bank, Gulf of Mexico. Bull. Mar. Sci. 95, 239–250. doi:10.5343/bms.2018.0057.

Kealoha, A. K. (2019). Coral Reef Ecosystem Health in Response to Climate Change and Environmental Stressors. Available at: https://oaktrust.library.tamu.edu/handle/1969.1/185032 [Accessed March 26, 2020].

Kealoha, A. K., Doyle, S. M., Shamberger, K. E. F., Sylvan, J. B., Hetland, R. D., and DiMarco, S. F. (2020). Localized hypoxia may have caused coral reef mortality at the Flower Garden Banks. Coral Reefs 39, 119–132. doi:10.1007/s00338-019-01883-9.

Kerswell, A. P., and Jones, R. J. (2003). Effects of hypo-osmosis on the coral Stylophora pistillata: nature and cause of ‘low-salinity bleaching.’ Mar. Ecol. Prog. Ser. 253, 145–154. doi:10.3354/meps253145.

Kiaghadi, A., and Rifai, H. S. (2019). Physical, Chemical, and Microbial Quality of Floodwaters in Houston Following Hurricane Harvey. Environ. Sci. Technol. 53, 4832–4840. doi:10.1021/acs.est.9b00792.

Knight, P. A., and Fell, P. E. (1987). Low salinity induces reversible tissue regression in the estuarine sponge Microciona prolifera (Ellis & Solander). J. Exp. Mar. Bio. Ecol. 107, 253–261. doi:10.1016/0022-0981(87)90042-6.

Knutson, T. R., McBride, J. L., Chan, J., Emanuel, K., Holland, G., Landsea, C., et al. (2010). Tropical cyclones and climate change. Nat. Geosci. 3, 157–163. doi:10.1038/ngeo779.

Lapointe, B. E., Langton, R., Bedford, B. J., Potts, A. C., Day, O., and Hu, C. (2010). Land-based nutrient enrichment of the Buccoo Reef Complex and fringing coral reefs of Tobago, West Indies. Mar. Pollut. Bull. 60, 334–343. doi:10.1016/j.marpolbul.2009.10.020.

Le Hénaff, M., Muller-Karger, F. E., Kourafalou, V. H., Otis, D., Johnson, K. A., McEachron, L., et al. (2019). Coral mortality event in the Flower Garden Banks of the Gulf of Mexico in July 2016: Local hypoxia due to cross-shelf transport of coastal flood waters? Cont. Shelf Res. 190, 103988. doi:10.1016/j.csr.2019.103988.

Longo, C., Corriero, G., Licciano, M., and Stabili, L. (2010). Bacterial accumulation by the Demospongiae Hymeniacidon perlevis: A tool for the bioremediation of polluted seawater. Mar. Pollut. Bull. 60, 1182–1187. doi:10.1016/j.marpolbul.2010.03.035.

López-Legentil, S., Erwin, P. M., Pawlik, J. R., and Song, B. (2010). Effects of Sponge Bleaching on Ammonia-Oxidizing Archaea: Distribution and Relative Expression of Ammonia Monooxygenase Genes Associated with the Barrel Sponge Xestospongia muta. Microb. Ecol. 60, 561–571. doi:10.1007/s00248-010-9662-1.

Luter, H. M., Gibb, K., and Webster, N. S. (2014). Eutrophication has no short-term effect on the Cymbastela stipitata holobiont. Front. Microbiol. 5, 216. doi:10.3389/fmicb.2014.00216.

Lynch, T. C., and Phlips, E. J. (2000). Filtration of the bloom-forming cyanobacteria Synechococcus by three sponge species from Florida Bay, U.S.A. Bull. Mar. Sci. 67, 923–936.

Maldonado, M., Zhang, X., Cao, X., Xue, L., Cao, H., and Zhang, W. (2010). Selective feeding by sponges on pathogenic microbes: A reassessment of potential for abatement of microbial pollution. Mar. Ecol. Prog. Ser. 403, 75–89. doi:10.3354/meps08411.

McMurray, S. E., Blum, J. E., and Pawlik, J. R. (2008). Redwood of the reef: Growth and age of the giant barrel sponge Xestospongia muta in the Florida Keys. Mar. Biol. 155, 159–171. doi:10.1007/s00227-008-1014-z.

Moeller, F. U., Webster, N. S., Herbold, C. W., Behnam, F., Domman, D., Albertsen, M., et al. (2019). Characterization of a thaumarchaeal symbiont that drives incomplete nitrification in the tropical sponge Ianthella basta. Environ. Microbiol. 21, 3831–3854. doi:10.1111/1462-2920.14732.

Moitinho-Silva, L., Nielsen, S., Amir, A., Gonzalez, A., Ackermann, G. L., Cerrano, C., et al. (2017). The sponge microbiome project. Gigascience 6, 1–7. doi:10.1093/gigascience/gix077.

Montalvo, N. F., Davis, J., Vicente, J., Pittiglio, R., Ravel, J., and Hill, R. T. (2014). Integration of culture-based and molecular analysis of a complex sponge-associated bacterial community. PLoS One 9, 1–8. doi:10.1371/journal.pone.0090517.

Morganti, T. M., Ribes, M., Yahel, G., and Coma, R. (2019). Size Is the Major Determinant of Pumping Rates in Marine Sponges. Front. Physiol. 10. doi:10.3389/fphys.2019.01474.

Morrow, K. M., Fiore, C. L., and Lesser, M. P. (2016). Environmental drivers of microbial community shifts in the giant barrel sponge, Xestospongia muta, over a shallow to mesophotic depth gradient. Environ. Microbiol. 18, 2025–2038. doi:10.1111/1462-2920.13226.

Nelson, H. R., and Altieri, A. H. (2019). Oxygen: the universal currency on coral reefs. Coral Reefs 38, 177–198. doi:10.1007/s00338-019-01765-0.

Oakley, J. W., and Guillen, G. J. (2019). Impact of Hurricane Harvey on Galveston Bay Saltmarsh Nekton Communities. Estuaries and Coasts. doi:10.1007/s12237-019-00581-7.

Olson, J. B., and Gao, X. (2013). Characterizing the bacterial associates of three Caribbean sponges along a gradient from shallow to mesophotic depths. FEMS Microbiol. Ecol. 85, 74–84. doi:10.1111/1574-6941.12099.

Ostrander, C. E., McManus, M. A., DeCarlo, E. H., and Mackenzie, F. T. (2008). Temporal and spatial variability of freshwater plumes in a semienclosed estuarine-bay system. Estuaries and Coasts 31, 192–203. doi:10.1007/s12237-007-9001-z.

Parra-velandia, F. J., Zea, S., and Soest, R. W. M. V. S. (2014). Reef sponges of the genus Agelas (Porifera: Demospongiae) from the Greater Caribbean. Zootaxa 3794, 301–343. doi:10.11646/zootaxa.3794.3.1.

Partensky, F., Blanchot, J., and Vaulot, D. (1999). Differential distribution and ecology of Prochlorococcus and Synechococcus in oceanic waters: a review. Bull. Oceanogr. Monaco-Numero Spec. 19, 457–476. Available at: http://cat.inist.fr/?aModele=afficheN&cpsidt=1218663.

Pawlik, J. R., and McMurray, S. E. (2020). The Emerging Ecological and Biogeochemical Importance of Sponges on Coral Reefs. Ann. Rev. Mar. Sci. 12, 315–337. doi:10.1146/annurev-marine-010419-010807.

Pita, L., Rix, L., Slaby, B. M., Franke, A., and Hentschel, U. (2018). The sponge holobiont in a changing ocean: from microbes to ecosystems. Microbiome 6, 46. doi:10.1186/s40168-018-0428-1.

Quast, C., Pruesse, E., Yilmaz, P., Gerken, J., Schweer, T., Yarza, P., et al. (2013). The SILVA ribosomal RNA gene database project: Improved data processing and web-based tools. Nucleic Acids Res. 41, 590–596. doi:10.1093/nar/gks1219.

Radax, R., Hoffmann, F., Rapp, H. T., Leininger, S., and Schleper, C. (2012). Ammonia-oxidizing archaea as main drivers of nitrification in cold-water sponges. Environ. Microbiol. 14, 909–923. doi:10.1111/j.1462-2920.2011.02661.x.

Roffer, M. A., Gawlikowski, G., and Upton, M. (2018). Monitoring of the Hurricane Harvey Plume in the Gulf of Mexico. West Melbourne, FL 32904.

Rua, C. P. J., de Oliveira, L. S., Froes, A., Tschoeke, D. A., Soares, A. C., Leomil, L., et al. (2018). Microbial and Functional Biodiversity Patterns in Sponges that Accumulate Bromopyrrole Alkaloids Suggest Horizontal Gene Transfer of Halogenase Genes. Microb. Ecol. 76, 825–838. doi:10.1007/s00248-018-1172-6.

Schmitt, S., Tsai, P., Bell, J., Fromont, J., Ilan, M., Lindquist, N., et al. (2012). Assessing the complex sponge microbiota: Core, variable and species-specific bacterial communities in marine sponges. ISME J. 6, 564–576. doi:10.1038/ismej.2011.116.

Shore, A. N., Sims, J. A., Grimes, M., Howe-Kerr, L. I., Stadler, L., Sylvan, J. B., et al. (2020). On a reef far, far away: Offshore transport of floodwaters following extreme storms impacts sponge health and associated microbial communities. bioRxiv, 2020.04.27.064568. doi:10.1101/2020.04.27.064568.

Simister, R., Taylor, M. W., Tsai, P., and Webster, N. (2012). Sponge-Microbe Associations Survive High Nutrients and Temperatures. PLoS One 7, e52220. doi:10.1371/journal.pone.0052220.

Slaby, B. M., Franke, A., Rix, L., Pita, L., Bayer, K., Jahn, M. T., et al. (2019). “Marine Sponge Holobionts in Health and Disease,” in Symbiotic Microbiomes of Coral Reefs Sponges and Corals (Springer Netherlands), 81–104. doi:10.1007/978-94-024-1612-1_7.

Sutherland, K. P., Porter, J. W., Turner, J. W., Thomas, B. J., Looney, E. E., Luna, T. P., et al. (2010). Human sewage identified as likely source of white pox disease of the threatened Caribbean elkhorn coral, Acropora palmata. Environ. Microbiol. 12, 1122–1131. doi:10.1111/j.1462-2920.2010.02152.x.

Sutherland, K. P., Shaban, S., Joyner, J. L., Porter, J. W., and Lipp, E. K. (2011). Human pathogen shown to cause disease in the threatened eklhorn coral Acropora palmata. PLoS One 6. doi:10.1371/journal.pone.0023468.

Szmant, A. M. (2002). Nutrient enrichment on coral reefs: Is it a major cause of coral reef decline? Estuaries 25, 743–766. doi:10.1007/BF02804903.

Villegas-Plazas, M., Wos-Oxley, M. L., Sanchez, J. A., Pieper, D. H., Thomas, O. P., and Junca, H. (2019). Variations in Microbial Diversity and Metabolite Profiles of the Tropical Marine Sponge Xestospongia muta with Season and Depth. Microb. Ecol. 78, 243–256. doi:10.1007/s00248-018-1285-y.

Wallace, C. C., Yund, P. O., Ford, T. E., Matassa, K. A., and Bass, A. L. (2013). Increase in antimicrobial resistance in bacteria isolated from stranded marine mammals of the Northwest Atlantic. Ecohealth 10, 201–210. doi:10.1007/s10393-013-0842-6.

Wear, S. L., and Thurber, R. V. (2015). Sewage pollution: mitigation is key for coral reef stewardship. Ann. N. Y. Acad. Sci. 1355, 15–30. doi:10.1111/nyas.12785.

Whitman, W. B. (2015). Bergey’s manual of systematics of archaea and bacteria., eds. P. DeVos, J. Chun, S. Dedysh, B. Hedlund, P. Kämpfer, F. Rainey, et al. Hoboken, NJ: Wiley doi:10.1002/9781118960608.

Wright, R. M., Correa, A. M. S., Quigley, L. A., Santiago-Vázquez, L. Z., Shamberger, K. E. F., and Davies, S. W. (2019). Gene Expression of Endangered Coral (Orbicella spp.) in Flower Garden Banks National Marine Sanctuary After Hurricane Harvey. Front. Mar. Sci. 6, 672. doi:10.3389/fmars.2019.00672.

Yu, P., Zaleski, A., Li, Q., He, Y., Mapili, K., Pruden, A., et al. (2018). Elevated Levels of Pathogenic Indicator Bacteria and Antibiotic Resistance Genes after Hurricane Harvey’s Flooding in Houston. Environ. Sci. Technol. Lett. 5, 481–486. doi:10.1021/acs.estlett.8b00329.

Zaneveld, J. R., McMinds, R., and Thurber, R. V. (2017). Stress and stability: Applying the Anna Karenina principle to animal microbiomes. Nat. Microbiol. 2. doi:10.1038/nmicrobiol.2017.121.

