## Supplemental Tables and Figure for "On a Reef Far, Far Away: Anthropogenic Impacts Following Extreme Storms Affect Sponge Health and Bacterial Communities"

**Supplemental Information**

<u>Tables</u>

-Supplemental Table S1. Metadata for all samples collected

-Supplemental Table S2. ANOSIM comparisons of the weighted UniFRAC distance matrix values for sponge bacterial communities by health state

-Supplemental Table S3. ANOSIM comparisons of the weighted UniFRAC distance matrix values for healthy sponge bacterial communities by site

-Supplemental Table S4. ANOSIM comparisons of the weighted UniFRAC distance matrix values for healthy sponge communities by collection date

-Supplemental Table S5. Differentially abundant bacterial Families in healthy sponges and in seawater samples determined by DESeq2

-Supplemental Table S6. Bacterial taxa significantly associated with recent flooding events according to Indicator Species Analysis

-Supplemental Table S7. Bacterial strains, primers, and annealing conditions used for qPCR analysis

<u>Figures</u>

-Supplemental Figure S1. Percent abundance of bacterial Families in a) healthy *Xestospongia muta* b) healthy *Agelas clathrodes*, c) Seawater, and d) diseased sponge samples

.

**Supplemental Table S1.** Sample Metadata

| Sample ID | Species | Collection Date | Site* | Depth (m) | Sample Type |
|---|---|---|---|---|---|
| AAEa61 | *Agelas clathrodes* | 6 August 2016 | East Bank - Buoy 4 | 19.8 - 22.9 | Diseased sponge |
| AAEa62 | *Agelas clathrodes* | 6 August 2016 | East Bank - Buoy 4 | 19.8 - 22.9 | Diseased sponge |
| AAEa63 | *Agelas clathrodes* | 6 August 2016 | East Bank - Buoy 4 | 19.8 - 22.9 | Diseased sponge |
| AAEa64 | *Agelas clathrodes* | 6 August 2016 | East Bank - Buoy 4 | 19.8 - 22.9 | Diseased sponge |
| XAEa61 | *Xestospongia muta* | 6 August 2016 | East Bank - Buoy 4 | 19.8 - 22.9 | Diseased sponge |
| XAEa62 | *Xestospongia muta* | 6 August 2016 | East Bank - Buoy 4 | 19.8 - 22.9 | Diseased sponge |
| XAEa63 | *Xestospongia muta* | 6 August 2016 | East Bank - Buoy 4 | 19.8 - 22.9 | Diseased sponge |
| XAEa64 | *Xestospongia muta* | 6 August 2016 | East Bank - Buoy 4 | 19.8 - 22.9 | Diseased sponge |
| XAEa65 | *Xestospongia muta* | 6 August 2016 | East Bank - Buoy 4 | 19.8 - 22.9 | Diseased sponge |
| AHEa61 | *Agelas clathrodes* | 6 August 2016 | East Bank - Buoy 4 | 19.8 - 22.9 | Visually healthy sponge |
| AHEa62 | *Agelas clathrodes* | 6 August 2016 | East Bank - Buoy 4 | 19.8 - 22.9 | Visually healthy sponge |
| AHEa63 | *Agelas clathrodes* | 6 August 2016 | East Bank - Buoy 4 | 19.8 - 22.9 | Visually healthy sponge |
| AHEa64 | *Agelas clathrodes* | 6 August 2016 | East Bank - Buoy 4 | 19.8 - 22.9 | Visually healthy sponge |
| AHEa65 | *Agelas clathrodes* | 6 August 2016 | East Bank - Buoy 4 | 19.8 - 22.9 | Visually healthy sponge |
| AHEa66 | *Agelas clathrodes* | 6 August 2016 | East Bank - Buoy 4 | 19.8 - 22.9 | Visually healthy sponge |
| AHEa67 | *Agelas clathrodes* | 6 August 2016 | East Bank - Buoy 4 | 19.8 - 22.9 | Visually healthy sponge |
| AHEa68 | *Agelas clathrodes* | 6 August 2016 | East Bank - Buoy 4 | 19.8 - 22.9 | Visually healthy sponge |
| AHWa61 | *Agelas clathrodes* | 6 August 2016 | West Bank - Buoy 2 | 19.8 - 22.9 | Visually healthy sponge |
| AHWa62 | *Agelas clathrodes* | 6 August 2016 | West Bank - Buoy 2 | 19.8 - 22.9 | Visually healthy sponge |
| AHWa63 | *Agelas clathrodes* | 6 August 2016 | West Bank - Buoy 2 | 19.8 - 22.9 | Visually healthy sponge |
| AHWa64 | *Agelas clathrodes* | 6 August 2016 | West Bank - Buoy 2 | 19.8 - 22.9 | Visually healthy sponge |
| AHWa65 | *Agelas clathrodes* | 6 August 2016 | West Bank - Buoy 2 | 19.8 - 22.9 | Visually healthy sponge |
| XHEa61 | *Xestospongia muta* | 6 August 2016 | East Bank - Buoy 4 | 19.8 - 22.9 | Visually healthy sponge |
| XHEa62 | *Xestospongia muta* | 6 August 2016 | East Bank - Buoy 4 | 19.8 - 22.9 | Visually healthy sponge |
| XHWa61 | *Xestospongia muta* | 6 August 2016 | West Bank - Buoy 2 | 19.8 - 22.9 | Visually healthy sponge |
| XHWa62 | *Xestospongia muta* | 6 August 2016 | West Bank - Buoy 2 | 19.8 - 22.9 | Visually healthy sponge |
| XHWa63 | *Xestospongia muta* | 6 August 2016 | West Bank - Buoy 2 | 19.8 - 22.9 | Visually healthy sponge |
| SCGa61 | Seawater | 30 July - 2 August 2016 | West Bank - KG1 | 0.1 | Seawater column, surface |
| SCGa62 | Seawater | 30 July - 2 August 2016 | East Bank - KG3 | 0.1 | Seawater column, surface |
| SCGa63 | Seawater | 30 July - 2 August 2016 | West Bank - KG1 | 30 | Seawater column, deep |
| SCGa64 | Seawater | 30 July - 2 August 2016 | East Bank - KG2 | 30 | Seawater column, deep |
| XHEs70 | *Xestospongia muta* | 16 September 2017 | East Bank - Buoy 4 | 19.8 - 22.9 | Visually healthy sponge |
| XHEs71 | *Xestospongia muta* | 16 September 2017 | East Bank - Buoy 4 | 19.8 - 22.9 | Visually healthy sponge |
| XHEs73 | *Xestospongia muta* | 16 September 2017 | East Bank - Buoy 4 | 19.8 - 22.9 | Visually healthy sponge |
| XHEs74 | *Xestospongia muta* | 16 September 2017 | East Bank - Buoy 4 | 19.8 - 22.9 | Visually healthy sponge |
| XHEs75 | *Xestospongia muta* | 16 September 2017 | East Bank - Buoy 4 | 19.8 - 22.9 | Visually healthy sponge |
| XHEs76 | *Xestospongia muta* | 16 September 2017 | East Bank - Buoy 4 | 19.8 - 22.9 | Visually healthy sponge |
| XHEs77 | *Xestospongia muta* | 16 September 2017 | East Bank - Buoy 4 | 19.8 - 22.9 | Visually healthy sponge |
| XHEs78 | *Xestospongia muta* | 16 September 2017 | East Bank - Buoy 4 | 19.8 - 22.9 | Visually healthy sponge |
| XHEs79 | *Xestospongia muta* | 16 September 2017 | East Bank - Buoy 4 | 19.8 - 22.9 | Visually healthy sponge |
| AHEo71 | *Agelas clathrodes* | 21 October 2017 | East Bank - Buoy 4 | 18.9 | Visually healthy sponge |
| AHEo72 | *Agelas clathrodes* | 21 October 2017 | East Bank - Buoy 4 | 20.1 | Visually healthy sponge |
| AHEo73 | *Agelas clathrodes* | 21 October 2017 | East Bank - Buoy 4 | 19.8 | Visually healthy sponge |
| AHEo74 | *Agelas clathrodes* | 21 October 2017 | East Bank - Buoy 4 | 20.4 | Visually healthy sponge |
| AHEo75 | *Agelas clathrodes* | 21 October 2017 | East Bank - Buoy 4 | 20.7 | Visually healthy sponge |
| AHEo76 | *Agelas clathrodes* | 21 October 2017 | East Bank - Buoy 4 | 21.3 | Visually healthy sponge |
| AHEo77 | *Agelas clathrodes* | 21 October 2017 | East Bank - Buoy 4 | 18.6 | Visually healthy sponge |
| AHEo78 | *Agelas clathrodes* | 21 October 2017 | East Bank - Buoy 4 | 18.3 | Visually healthy sponge |
| AHEo79 | *Agelas clathrodes* | 21 October 2017 | East Bank - Buoy 4 | 19.2 | Visually healthy sponge |
| AHWo71 | *Agelas clathrodes* | 21 - 24 October 2017 | West Bank - Buoy 2 | 24.1 | Visually healthy sponge |
| AHWo72 | *Agelas clathrodes* | 21 - 24 October 2017 | West Bank - Buoy 2 | 24.4 | Visually healthy sponge |
| AHWo73 | *Agelas clathrodes* | 21 - 24 October 2017 | West Bank - Buoy 2 | 25.6 | Visually healthy sponge |

| | | | | | |
|---|---|---|---|---|---|
| AHWo74 | *Agelas clathrodes* | 21 - 24 October 2017 | West Bank - Buoy 2 | 26.8 | Visually healthy sponge |
| AHWo75 | *Agelas clathrodes* | 21 - 24 October 2017 | West Bank - Buoy 2 | 28 | Visually healthy sponge |
| AHWo76 | *Agelas clathrodes* | 21 - 24 October 2017 | West Bank - Buoy 2 | 25.6 | Visually healthy sponge |
| AHWo77 | *Agelas clathrodes* | 21 - 24 October 2017 | West Bank - Buoy 2 | 22.6 | Visually healthy sponge |
| AHWo78 | *Agelas clathrodes* | 21 - 24 October 2017 | West Bank - Buoy 2 | 24.4 | Visually healthy sponge |
| XHEo71 | *Xestospongia muta* | 21 October 2017 | East Bank - Buoy 4 | 20.1 | Visually healthy sponge |
| XHEo72 | *Xestospongia muta* | 21 October 2017 | East Bank - Buoy 4 | 20.7 | Visually healthy sponge |
| XHWo71 | *Xestospongia muta* | 21 October 2017 | West Bank - Buoy 2 | 25.3 | Visually healthy sponge |
| XHWo72 | *Xestospongia muta* | 21 October 2017 | West Bank - Buoy 2 | 22.6 | Visually healthy sponge |
| XHWo73 | *Xestospongia muta* | 21 October 2017 | West Bank - Buoy 2 | 24.4 | Visually healthy sponge |
| SREo71 | Seawater | 20 - 25 October 2017 | East Bank - Buoy 4 | 21.3 | Seawater column, reef bottom |
| SREo72 | Seawater | 20 - 25 October 2017 | East Bank - Buoy 4 | 20.1 | Seawater column, reef bottom |
| SREo73 | Seawater | 20 - 25 October 2017 | East Bank - Buoy 4 | 20.1 | Seawater column, reef bottom |
| SREo74 | Seawater | 20 - 25 October 2017 | East Bank - Buoy 4 | 20.7 | Seawater column, reef bottom |
| SREo75 | Seawater | 20 - 25 October 2017 | East Bank - Buoy 4 | 18.9 | Seawater column, reef bottom |
| SRWo71 | Seawater | 20 - 25 October 2017 | West Bank - Buoy 2 | 26.8 | Seawater column, reef bottom |
| SRWo72 | Seawater | 20 - 25 October 2017 | West Bank - Buoy 2 | 24.4 | Seawater column, reef bottom |
| SRWo73 | Seawater | 20 - 25 October 2017 | West Bank - Buoy 2 | 22.6 | Seawater column, reef bottom |
| SRWo74 | Seawater | 20 - 25 October 2017 | West Bank - Buoy 2 | 24.4 | Seawater column, reef bottom |
| SRWo75 | Seawater | 20 - 25 October 2017 | West Bank - Buoy 2 | 22.6 | Seawater column, reef bottom |
| SRWo76 | Seawater | 20 - 25 October 2017 | West Bank - Buoy 2 | 23.5 | Seawater column, reef bottom |
| SRWo77 | Seawater | 20 - 25 October 2017 | West Bank - Buoy 2 | 25.3 | Seawater column, reef bottom |
| SRWo78 | Seawater | 20 - 25 October 2017 | West Bank - Buoy 2 | 24.1 | Seawater column, reef bottom |
| SCEo71 | Seawater | 20 - 25 October 2017 | East Bank - Buoy 4 | 18.0 | Seawater column deep |
| SCEo72 | Seawater | 20 - 25 October 2017 | East Bank - Buoy 4 | 10.0 | Seawater column, mid-depth |
| SCEo73 | Seawater | 20 - 25 October 2017 | East Bank - Buoy 4 | 3.2 | Seawater column, surface |
| SCWo71 | Seawater | 20 - 25 October 2017 | West Bank - Buoy 2 | 16.0 | Seawater column, deep |
| SCWo72 | Seawater | 20 - 25 October 2017 | West Bank - Buoy 2 | 10.0 | Seawater column, mid-depth |
| SCWo73 | Seawater | 20 - 25 October 2017 | West Bank - Buoy 2 | 3.4 | Seawater column, surface |
| SCGo71 | Seawater | 20 - 25 October 2017 | West Bank - KG1 | 3.6 | Seawater column, surface |
| SCGo72 | Seawater | 20 - 25 October 2017 | East Bank - KG2 | 2.9 | Seawater column, surface |
| SCGo73 | Seawater | 20 - 25 October 2017 | East Bank - KG3 | 5.0 | Seawater column, surface |
| SCGo74 | Seawater | 20 - 25 October 2017 | West Bank - KG1 | 25.7 | Seawater column, deep |
| SCGo75 | Seawater | 20 - 25 October 2017 | East Bank - KG2 | 24.7 | Seawater column, deep |
| SCGo76 | Seawater | 20 - 25 October 2017 | East Bank - KG3 | 24.7 | Seawater column, deep |
| AHEo80 | *Agelas clathrodes* | 25 October 2018 | East Bank - Buoy 4 | 18.3 | Visually healthy sponge |
| AHEo81 | *Agelas clathrodes* | 25 October 2018 | East Bank - Buoy 4 | 18.3 | Visually healthy sponge |
| AHEo82 | *Agelas clathrodes* | 25 October 2018 | East Bank - Buoy 4 | 16.5 | Visually healthy sponge |
| AHEo83 | *Agelas clathrodes* | 25 October 2018 | East Bank - Buoy 4 | 18.6 | Visually healthy sponge |
| AHEo84 | *Agelas clathrodes* | 25 October 2018 | East Bank - Buoy 4 | 19.5 | Visually healthy sponge |
| AHEo85 | *Agelas clathrodes* | 25 October 2018 | East Bank - Buoy 4 | 18.9 | Visually healthy sponge |
| AHEo86 | *Agelas clathrodes* | 25 October 2018 | East Bank - Buoy 4 | 18.6 | Visually healthy sponge |
| AHEo87 | *Agelas clathrodes* | 25 October 2018 | East Bank - Buoy 4 | 19.2 | Visually healthy sponge |
| AHEo88 | *Agelas clathrodes* | 25 October 2018 | East Bank - Buoy 4 | 19.5 | Visually healthy sponge |
| AHEo89 | *Agelas clathrodes* | 25 October 2018 | East Bank - Buoy 4 | 18.9 | Visually healthy sponge |
| AHWo80 | *Agelas clathrodes* | 26 October 2018 | West Bank - Buoy 2 | 24.7 | Visually healthy sponge |
| AHWo81 | *Agelas clathrodes* | 26 October 2018 | West Bank - Buoy 2 | 24.4 | Visually healthy sponge |
| AHWo82 | *Agelas clathrodes* | 26 October 2018 | West Bank - Buoy 2 | 23.2 | Visually healthy sponge |
| AHWo83 | *Agelas clathrodes* | 26 October 2018 | West Bank - Buoy 2 | 22.6 | Visually healthy sponge |
| AHWo84 | *Agelas clathrodes* | 26 October 2018 | West Bank - Buoy 2 | 22.6 | Visually healthy sponge |
| AHWo85 | *Agelas clathrodes* | 26 October 2018 | West Bank - Buoy 2 | 22.9 | Visually healthy sponge |
| AHWo86 | *Agelas clathrodes* | 26 October 2018 | West Bank - Buoy 2 | 21.9 | Visually healthy sponge |
| AHWo87 | *Agelas clathrodes* | 26 October 2018 | West Bank - Buoy 2 | 21.9 | Visually healthy sponge |
| AHWo88 | *Agelas clathrodes* | 27 October 2018 | West Bank - Buoy 2 | 21.6 | Visually healthy sponge |

| | | | | | |
|---|---|---|---|---|---|
| AHWo89 | *Agelas clathrodes* | 28 October 2018 | West Bank - Buoy 2 | NA | Visually healthy sponge |
| XHEo80 | *Xestospongia muta* | 25 October 2018 | East Bank - Buoy 4 | 19.2 | Visually healthy sponge |
| XHEo81 | *Xestospongia muta* | 25 October 2018 | East Bank - Buoy 4 | 19.8 | Visually healthy sponge |
| XHEo82 | *Xestospongia muta* | 25 October 2018 | East Bank - Buoy 4 | 18.9 | Visually healthy sponge |
| XHEo83 | *Xestospongia muta* | 26 October 2018 | East Bank - Buoy 4 | 19.2 | Visually healthy sponge |
| XHEo84 | *Xestospongia muta* | 26 October 2018 | East Bank - Buoy 4 | 18.9 | Visually healthy sponge |
| XHEo85 | *Xestospongia muta* | 26 October 2018 | East Bank - Buoy 4 | 19.5 | Visually healthy sponge |
| XHEo86 | *Xestospongia muta* | 26 October 2018 | East Bank - Buoy 4 | 18.6 | Visually healthy sponge |
| XHEo87 | *Xestospongia muta* | 26 October 2018 | East Bank - Buoy 4 | 19.5 | Visually healthy sponge |
| XHEo88 | *Xestospongia muta* | 26 October 2018 | East Bank - Buoy 4 | 19.2 | Visually healthy sponge |
| XHEo89 | *Xestospongia muta* | 26 October 2018 | East Bank - Buoy 4 | 20.1 | Visually healthy sponge |
| XHWo80 | *Xestospongia muta* | 27 October 2018 | West Bank - Buoy 2 | 23.5 | Visually healthy sponge |
| XHWo81 | *Xestospongia muta* | 27 October 2018 | West Bank - Buoy 2 | 24.4 | Visually healthy sponge |
| XHWo82 | *Xestospongia muta* | 27 October 2018 | West Bank - Buoy 2 | 23.8 | Visually healthy sponge |
| XHWo83 | *Xestospongia muta* | 27 October 2018 | West Bank - Buoy 2 | 22.6 | Visually healthy sponge |
| XHWo84 | *Xestospongia muta* | 27 October 2018 | West Bank - Buoy 2 | 22.6 | Visually healthy sponge |
| XHWo85 | *Xestospongia muta* | 27 October 2018 | West Bank - Buoy 2 | 23.2 | Visually healthy sponge |
| XHWo86 | *Xestospongia muta* | 27 October 2018 | West Bank - Buoy 2 | 23.2 | Visually healthy sponge |
| XHWo87 | *Xestospongia muta* | 27 October 2018 | West Bank - Buoy 2 | 24.1 | Visually healthy sponge |
| XHWo88 | *Xestospongia muta* | 27 October 2018 | West Bank - Buoy 2 | 22.2 | Visually healthy sponge |
| XHWo89 | *Xestospongia muta* | 28 October 2018 | West Bank - Buoy 2 | NA | Visually healthy sponge |
| SCEo81 | Seawater | 22 - 27 October 2018 | East Bank - Buoy 4 | 17.5 | Seawater column, deep |
| SCEo82 | Seawater | 22 - 27 October 2018 | East Bank - Buoy 4 | 10.0 | Seawater column, mid-depth |
| SCEo83 | Seawater | 22 - 27 October 2018 | East Bank - Buoy 4 | 3.5 | Seawater column, surface |
| SCWo81 | Seawater | 22 - 27 October 2018 | West Bank - Buoy 2 | 19.0 | Seawater column, deep |
| SCWo82 | Seawater | 22 - 27 October 2018 | West Bank - Buoy 2 | 3.5 | Seawater column, surface |
| SCWo83 | Seawater | 22 - 27 October 2018 | West Bank - Buoy 2 | 10.0 | Seawater column, mid-depth |
| SCGo81 | Seawater | 22 - 27 October 2018 | East Bank - KG2 | 4 | Seawater column, surface |
| SCGo82 | Seawater | 22 - 27 October 2018 | East Bank - KG3 | 3.5 | Seawater column, surface |
| SCGo83 | Seawater | 22 - 27 October 2018 | West Bank - KG1 | 25.0 | Seawater column, deep |
| SCGo84 | Seawater | 22 - 27 October 2018 | East Bank - KG2 | 24.5 | Seawater column, deep |
| SCGo85 | Seawater | 22 - 27 October 2018 | East Bank - KG3 | 32.5 | Seawater column, deep |

**Supplemental Table S2**. ANOSIM comparisons (999 permutations) of the weighted UniFRAC distance matrix values for all sampled sponge and seawater communities based on health status and sponge species. Significant differences are shown in **bold**.

| Comparison | Global R | P value |
|---|---|---|
| **Healthy *Agelas* vs. Healthy *Xestospongia*** | **0.748** | **0.001** |
| **Healthy *Agelas* vs. Diseased *Agelas*** | **0.999** | **0.001** |
| **Healthy *Xestospongia* vs. Diseased *Xestospongia*** | **0.988** | **0.001** |
| **Healthy *Agelas* vs. Seawater** | **0.999** | **0.001** |
| **Healthy *Xestospongia* vs. Seawater** | **0.968** | **0.001** |
| Diseased *Agelas* vs. Diseased *Xestospongia* | 0.193 | 0.101 |
| **Seawater vs. Diseased *Agelas*** | **0.999** | **0.001** |
| **Seawater vs. Diseased *Xestospongia*** | **0.999** | **0.001** |

**Supplemental Table S3**. ANOSIM comparisons (999 permutations) of the weighted UniFRAC distance matrix values for healthy sponge communities by site - East Bank (EB) vs. West Bank (WB). No significant differences were detected.

| | Comparison | Global R | P value |
|---|---|---|---|
| Visually healthy *Agelas clathrodes* | All dates | 0.002 | 0.373 |
| | Aug. 2016 | 0.159 | 0.121 |
| | Oct. 2017 | 0.066 | 0.178 |
| | Oct. 2018 | 0.084 | 0.121 |
| Visually healthy *Xestospongia muta* | All dates | 0.057 | 0.099 |
| | Aug. 2016 | 0.250 | 0.326 |
| | Oct. 2017 | 0.166 | 0.269 |
| | Oct. 2018 | 0.048 | 0.269 |
| Seawater | All dates | 0.011 | 0.564 |
| | Aug. 2016 | 0.250 | 0.357 |
| | Oct. 2017 | 0.384 | 0.078 |
| | Oct. 2018 | 0.007 | 0.416 |

**Supplemental Table S4**. ANOSIM comparisons of the weighted UniFRAC distance matrix values for healthy sponge and seawater communities. Significant differences are shown in **bold**.

| | Comparison | Global R | P value |
|---|---|---|---|
| Visually healthy *Agelas clathrodes* | **All Dates** | **0.602** | **0.001** |
| | **Aug. 2016 vs. Oct. 2017** | **0.416** | **0.001** |
| | **Aug. 2016 vs. Oct. 2018** | **0.823** | **0.001** |
| | **Oct. 2017 vs. Oct. 2018** | **0.555** | **0.001** |
| Visually healthy *Xestospongia muta* | **All Dates** | **0.495** | **0.001** |
| | Aug. 2016 vs. Sept. 2017 | 0.131 | 0.163 |
| | **Aug. 2016 vs. Oct. 2017** | **0.544** | **0.016** |
| | **Aug. 2016 vs. Oct. 2018** | **0.653** | **0.003** |
| | Sept. 2017 vs. Oct. 2017 | 0.205 | 0.088 |
| | **Sept. 2017 vs. Oct. 2018** | **0.478** | **0.003** |
| | **Oct. 2017 vs. Oct. 2018** | **0.594** | **0.006** |
| Seawater | **All Dates** | **0.569** | **0.001** |
| | **Water Column Aug. 2016 vs. Water Column Oct. 2017** | **0.919** | **0.002** |
| | **Water Column Aug. 2016 vs. Reef Water Oct. 2017** | **0.657** | **0.004** |
| | **Water Column Aug. 2016 vs. Water Column Oct. 2018** | **0.615** | **0.009** |
| | **Water Column Oct. 2017 vs. Reef Water Oct. 2017** | **0.516** | **0.002** |
| | **Water Column Oct. 2017 vs. Water Column Oct. 2018** | **0.807** | **0.002** |
| | **Reef Water Oct. 2017 vs. Water Column Oct. 2018** | **0.387** | **0.002** |

**Supplemental Table S5.** Bacterial Families within healthy sponge and seawater samples differing significantly ($p < 0.05$) in abundance between collection dates as determined by DESeq2. A positive Log2 Fold Change indicates higher abundance in the first comparison.

| Comparison | Base Mean | Log2 Fold Change | lfcSE | Stat | Adjusted p-value | Phylum_Family |
|---|---|---|---|---|---|---|
| *X. muta*: Aug. 2016 vs. Oct. 2018 | 9.556 | 8.542 | 1.759 | 4.856 | 0.000 | Desulfobacterota_ Desulfovibrionaceae |
| *X. muta*: Aug. 2016 vs. Oct. 2018 | 12.148 | 6.564 | 2.191 | 2.996 | 0.022 | Patescibacteria_ Saccharimonadales |
| *X. muta*: Aug. 2016 vs. Oct. 2018 | 2.120 | 5.920 | 1.954 | 3.030 | 0.022 | Proteobacteria_ Vibrionaceae |
| *X. muta*: Aug. 2016 vs. Oct. 2018 | 7.730 | 5.328 | 1.520 | 3.505 | 0.006 | Proteobacteria_ SAR86_clade |
| *X. muta*: Aug. 2016 vs. Oct. 2018 | 4.661 | 4.994 | 1.466 | 3.408 | 0.008 | Proteobacteria_ AEGEAN-169_marine_group |
| *X. muta*: Aug. 2016 vs. Oct. 2018 | 75.413 | 3.802 | 1.275 | 2.982 | 0.022 | Bacteroidota_ Cyclobacteriaceae |
| *X. muta*: Aug. 2016 vs. Oct. 2018 | 3.890 | 3.759 | 1.124 | 3.344 | 0.009 | Bacteroidota_ Flavobacteriaceae |
| *X. muta*: Aug. 2016 vs. Oct. 2018 | 101.585 | 2.836 | 0.536 | 5.296 | 0.000 | Poribacteria_ Poribacteria |
| *X. muta*: Aug. 2016 vs. Oct. 2018 | 3376.908 | 2.750 | 0.752 | 3.655 | 0.004 | Cyanobacteria_ Cyanobiaceae |
| *X. muta*: Aug. 2016 vs. Oct. 2018 | 264.087 | 2.649 | 0.679 | 3.899 | 0.002 | Bdellovibrionota_ Bdellovibrionaceae |
| *X. muta*: Aug. 2016 vs. Oct. 2018 | 586.922 | 2.489 | 0.543 | 4.585 | 0.000 | Proteobacteria_ UBA10353_marine_group |
| *X. muta*: Aug. 2016 vs. Oct. 2018 | 1013.988 | -1.719 | 0.569 | -3.019 | 0.022 | Chloroflexi_ Caldilineaceae |
| *X. muta*: Aug. 2016 vs. Oct. 2018 | 3.009 | -5.790 | 1.920 | -3.015 | 0.022 | Actinobacteriota_ Bifidobacteriaceae |
| *X. muta*: Aug. 2016 vs. Oct. 2018 | 4.902 | -6.419 | 1.434 | -4.475 | 0.000 | Bacteroidota_ Prevotellaceae |
| *X. muta*: Aug. 2016 vs. Oct. 2018 | 8.018 | -7.348 | 1.091 | -6.733 | 0.000 | Firmicutes_ Enterococcaceae |
| *X. muta*: Aug. 2016 vs. Oct. 2018 | 45.406 | -7.435 | 0.888 | -8.369 | 0.000 | Bacteroidota_ Bacteroidaceae |
| *X. muta*: Sept. 2017 vs. Oct. 2018 | 31.839 | 8.893 | 2.639 | 3.370 | 0.006 | Bacteroidota_ Muribaculaceae |
| *X. muta*: Sept. 2017 vs. Oct. 2018 | 4.144 | 6.500 | 2.212 | 2.939 | 0.022 | Spirochaetota_ Leptospiraceae |
| *X. muta*: Sept. 2017 vs. Oct. 2018 | 3.437 | 6.074 | 1.449 | 4.193 | 0.001 | Proteobacteria_ Halieaceae |
| *X. muta*: Sept. 2017 vs. Oct. 2018 | 4.071 | 5.811 | 2.031 | 2.861 | 0.025 | Firmicutes_ Clostridiaceae |
| *X. muta*: Sept. 2017 vs. Oct. 2018 | 2.669 | 5.432 | 1.282 | 4.237 | 0.001 | Verrucomicrobiota_ Puniceicoccaceae |
| *X. muta*: Sept. 2017 vs. Oct. 2018 | 2.365 | 5.001 | 1.840 | 2.718 | 0.035 | Acidobacteriota_ Vicinamibacteraceae |
| *X. muta*: Sept. 2017 vs. Oct. 2018 | 9.653 | 4.695 | 1.560 | 3.010 | 0.019 | Verrucomicrobiota_ Akkermansiaceae |
| *X. muta*: Sept. 2017 vs. Oct. 2018 | 7.730 | 3.549 | 1.214 | 2.924 | 0.022 | Proteobacteria_ SAR86_clade |
| *X. muta*: Sept. 2017 vs. Oct. 2018 | 3.890 | 3.244 | 0.915 | 3.547 | 0.004 | Bacteroidota_ Flavobacteriaceae |
| *X. muta*: Sept. 2017 vs. Oct. 2018 | 101.585 | 1.471 | 0.424 | 3.467 | 0.005 | Poribacteria_ Poribacteria |
| *X. muta*: Sept. 2017 vs. Oct. 2018 | 436.043 | -1.109 | 0.427 | -2.595 | 0.048 | Acidobacteriota_ PAUC26f |
| *X. muta*: Sept. 2017 vs. Oct. 2018 | 1831.591 | -1.255 | 0.486 | -2.583 | 0.048 | Chloroflexi_ TK17 |
| *X. muta*: Sept. 2017 vs. Oct. 2018 | 271.245 | -1.413 | 0.414 | -3.411 | 0.005 | AncK6_ AncK6 |
| *X. muta*: Sept. 2017 vs. Oct. 2018 | 231.427 | -1.419 | 0.502 | -2.829 | 0.026 | Proteobacteria_ Puniceispirillales_Incertae_Sedis |
| *X. muta*: Sept. 2017 vs. Oct. 2018 | 1194.602 | -1.453 | 0.379 | -3.832 | 0.002 | Gemmatimonadota_ BD2-11_terrestrial_group |
| *X. muta*: Sept. 2017 vs. Oct. 2018 | 45.406 | -1.554 | 0.431 | -3.603 | 0.004 | Bacteroidota_ Bacteroidaceae |
| *X. muta*: Sept. 2017 vs. Oct. 2018 | 586.922 | -1.567 | 0.428 | -3.659 | 0.003 | Proteobacteria_ UBA10353_marine_group |

| | | | | | | |
|---|---|---|---|---|---|---|
| *X. muta*: Sept. 2017 vs. Oct. 2018 | 745.857 | -1.595 | 0.435 | -3.670 | 0.003 | Proteobacteria_ Rhodobacteraceae |
| *X. muta*: Sept. 2017 vs. Oct. 2018 | 1312.880 | -2.251 | 0.452 | -4.974 | 0.000 | Crenarchaeota_ Nitrosopumilaceae |
| *X. muta*: Sept. 2017 vs. Oct. 2018 | 423.423 | -2.554 | 0.562 | -4.542 | 0.000 | Proteobacteria_ JTB23 |
| *X. muta*: Oct. 2017 vs. Oct. 2018 | 744.323 | 3.809 | 0.729 | 5.224 | 0.000 | Chloroflexi_ TK10 |
| *X. muta*: Oct. 2017 vs. Oct. 2018 | 1831.591 | 2.668 | 0.624 | 4.275 | 0.000 | Chloroflexi_ TK17 |
| *X. muta*: Oct. 2017 vs. Oct. 2018 | 99.454 | 2.624 | 0.682 | 3.847 | 0.002 | Chloroflexi_ JG30-KF-CM66 |
| *X. muta*: Oct. 2017 vs. Oct. 2018 | 1324.597 | 2.558 | 0.597 | 4.285 | 0.000 | Chloroflexi_ A4b |
| *X. muta*: Oct. 2017 vs. Oct. 2018 | 1013.988 | 1.631 | 0.569 | 2.867 | 0.035 | Chloroflexi_ Caldilineaceae |
| *X. muta*: Oct. 2017 vs. Oct. 2018 | 1008.358 | 1.554 | 0.574 | 2.705 | 0.045 | Acidobacteriota_ Thermoanaerobaculaceae |
| *X. muta*: Oct. 2017 vs. Oct. 2018 | 316.250 | -1.382 | 0.499 | -2.768 | 0.040 | Dadabacteria_ Dadabacteriales |
| *X. muta*: Oct. 2017 vs. Oct. 2018 | 735.940 | -1.522 | 0.548 | -2.778 | 0.040 | Bacteroidota_ Rhodothermaceae |
| *X. muta*: Oct. 2017 vs. Oct. 2018 | 350.653 | -1.892 | 0.543 | -3.486 | 0.005 | Proteobacteria_ Woeseiaceae |
| *X. muta*: Oct. 2017 vs. Oct. 2018 | 1312.880 | -2.495 | 0.582 | -4.289 | 0.000 | Crenarchaeota_ Nitrosopumilaceae |
| *X. muta*: Oct. 2017 vs. Oct. 2018 | 423.423 | -2.921 | 0.720 | -4.054 | 0.001 | Proteobacteria_ JTB23 |
| *X. muta*: Oct. 2017 vs. Oct. 2018 | 421.632 | -3.034 | 0.807 | -3.760 | 0.002 | Proteobacteria_ Pseudohongiellaceae |
| *X. muta*: Oct. 2017 vs. Oct. 2018 | 75.413 | -5.384 | 1.527 | -3.525 | 0.004 | Bacteroidota_ Cyclobacteriaceae |
| *X. muta*: Oct. 2017 vs. Oct. 2018 | 12.128 | -17.128 | 4.665 | -3.672 | 0.003 | Proteobacteria_ Methyloligellaceae |
| *A. clathrodes*: Aug. 2016 vs. Oct. 2018 | 309.031 | 9.330 | 0.848 | 11.008 | 0.000 | Desulfobacterota_ Desulfovibrionaceae |
| *A. clathrodes*: Aug. 2016 vs. Oct. 2018 | 22.553 | 8.838 | 0.713 | 12.396 | 0.000 | Porifera_ Demospongiae |
| *A. clathrodes*: Aug. 2016 vs. Oct. 2018 | 11.707 | 7.868 | 1.019 | 7.719 | 0.000 | Cyanobacteria_ Gastranaerophilales |
| *A. clathrodes*: Aug. 2016 vs. Oct. 2018 | 19.536 | 7.520 | 1.045 | 7.199 | 0.000 | Proteobacteria_ Stappiaceae |
| *A. clathrodes*: Aug. 2016 vs. Oct. 2018 | 19.574 | 7.334 | 0.983 | 7.461 | 0.000 | Firmicutes_ Clostridiaceae |
| *A. clathrodes*: Aug. 2016 vs. Oct. 2018 | 7.164 | 7.183 | 1.356 | 5.296 | 0.000 | Proteobacteria_ Beijerinckiaceae |
| *A. clathrodes*: Aug. 2016 vs. Oct. 2018 | 6.627 | 7.070 | 2.817 | 2.510 | 0.033 | Bacteroidota_ Lentimicrobiaceae |
| *A. clathrodes*: Aug. 2016 vs. Oct. 2018 | 4.454 | 6.464 | 1.123 | 5.756 | 0.000 | Firmicutes_ Peptostreptococcales-Tissierellales |
| *A. clathrodes*: Aug. 2016 vs. Oct. 2018 | 3.811 | 6.274 | 1.193 | 5.260 | 0.000 | Verrucomicrobiota_ P.palmC41 |
| *A. clathrodes*: Aug. 2016 vs. Oct. 2018 | 3.389 | 6.104 | 1.361 | 4.486 | 0.000 | Desulfobacterota_ Desulfolunaceae |
| *A. clathrodes*: Aug. 2016 vs. Oct. 2018 | 2.395 | 5.602 | 1.479 | 3.789 | 0.001 | Proteobacteria_ Thioglobaceae |
| *A. clathrodes*: Aug. 2016 vs. Oct. 2018 | 4.525 | 5.557 | 1.036 | 5.365 | 0.000 | Proteobacteria_ Sphingomonadaceae |
| *A. clathrodes*: Aug. 2016 vs. Oct. 2018 | 2.841 | 5.427 | 1.490 | 3.643 | 0.001 | Bdellovibrionota_ 0319-6G20 |
| *A. clathrodes*: Aug. 2016 vs. Oct. 2018 | 1.584 | 5.008 | 1.713 | 2.923 | 0.012 | Proteobacteria_ Kangiellaceae |
| *A. clathrodes*: Aug. 2016 vs. Oct. 2018 | 1.561 | 4.985 | 1.112 | 4.482 | 0.000 | Proteobacteria_ MBMPE27 |
| *A. clathrodes*: Aug. 2016 vs. Oct. 2018 | 1.973 | 4.872 | 1.440 | 3.385 | 0.003 | Proteobacteria_ Rhizobiaceae |
| *A. clathrodes*: Aug. 2016 vs. Oct. 2018 | 2.144 | 4.869 | 1.238 | 3.933 | 0.000 | Proteobacteria_ UBA10353_marine_group |
| *A. clathrodes*: Aug. 2016 vs. Oct. 2018 | 125.312 | 4.804 | 0.610 | 7.870 | 0.000 | Proteobacteria_ Enterobacteriaceae |
| *A. clathrodes*: Aug. 2016 vs. Oct. 2018 | 1.296 | 4.718 | 2.013 | 2.343 | 0.047 | Bacteroidota_ Prolixibacteraceae |
| *A. clathrodes*: Aug. 2016 vs. Oct. 2018 | 1.259 | 4.676 | 1.779 | 2.629 | 0.025 | Planctomycetota_ AKAU3564_sediment_group |
| *A. clathrodes*: Aug. 2016 vs. Oct. 2018 | 240.177 | 4.481 | 0.917 | 4.885 | 0.000 | Proteobacteria_ Endozoicomonadaceae |
| *A. clathrodes*: Aug. 2016 vs. Oct. 2018 | 2.463 | 4.460 | 1.363 | 3.271 | 0.004 | Proteobacteria_ Halieaceae |
| *A. clathrodes*: Aug. 2016 vs. Oct. 2018 | 0.838 | 4.090 | 1.480 | 2.763 | 0.018 | Campilobacterota_ Arcobacteraceae |

| | | | | | | |
|---|---|---|---|---|---|---|
| *A. clathrodes*: Aug. 2016 vs. Oct. 2018 | 1.004 | 4.066 | 1.676 | 2.426 | 0.039 | Bacteroidota_ Amoebophilaceae |
| *A. clathrodes*: Aug. 2016 vs. Oct. 2018 | 2.436 | 3.606 | 1.279 | 2.820 | 0.016 | Bacteroidota_ Saprospiraceae |
| *A. clathrodes*: Aug. 2016 vs. Oct. 2018 | 3.748 | 3.232 | 1.267 | 2.550 | 0.030 | Firmicutes_ Peptostreptococcaceae |
| *A. clathrodes*: Aug. 2016 vs. Oct. 2018 | 4.377 | 3.136 | 1.002 | 3.130 | 0.006 | Proteobacteria_ Vibrionaceae |
| *A. clathrodes*: Aug. 2016 vs. Oct. 2018 | 1.850 | 3.071 | 0.860 | 3.572 | 0.001 | SAR324 clade_Marine group B |
| *A. clathrodes*: Aug. 2016 vs. Oct. 2018 | 21.572 | 2.682 | 0.575 | 4.666 | 0.000 | Proteobacteria_ Coxiellaceae |
| *A. clathrodes*: Aug. 2016 vs. Oct. 2018 | 5.892 | 2.125 | 0.828 | 2.566 | 0.029 | Proteobacteria_ Clade_II |
| *A. clathrodes*: Aug. 2016 vs. Oct. 2018 | 3.478 | 2.112 | 0.855 | 2.471 | 0.035 | Proteobacteria_ Ectothiorhodospiraceae |
| *A. clathrodes*: Aug. 2016 vs. Oct. 2018 | 15.669 | 2.050 | 0.559 | 3.671 | 0.001 | Proteobacteria_ Clade_I |
| *A. clathrodes*: Aug. 2016 vs. Oct. 2018 | 58.177 | 1.612 | 0.405 | 3.979 | 0.000 | Poribacteria_ Poribacteria |
| *A. clathrodes*: Aug. 2016 vs. Oct. 2018 | 8.131 | 1.262 | 0.468 | 2.697 | 0.021 | Proteobacteria_ AEGEAN-169_marine_group |
| *A. clathrodes*: Aug. 2016 vs. Oct. 2018 | 199.036 | 1.081 | 0.435 | 2.486 | 0.034 | Proteobacteria_ Rhodobacteraceae |
| *A. clathrodes*: Aug. 2016 vs. Oct. 2018 | 764.128 | -1.073 | 0.390 | -2.749 | 0.019 | Acidobacteriota_ Subgroup_9 |
| *A. clathrodes*: Aug. 2016 vs. Oct. 2018 | 123.395 | -1.221 | 0.460 | -2.655 | 0.023 | PAUC34f_ PAUC34f |
| *A. clathrodes*: Aug. 2016 vs. Oct. 2018 | 971.763 | -1.343 | 0.309 | -4.345 | 0.000 | Bdellovibrionota_ Bdellovibrionaceae |
| *A. clathrodes*: Aug. 2016 vs. Oct. 2018 | 828.027 | -1.568 | 0.472 | -3.324 | 0.003 | Acidobacteriota_ PAUC26f |
| *A. clathrodes*: Aug. 2016 vs. Oct. 2018 | 118.711 | -1.606 | 0.681 | -2.358 | 0.046 | Patescibacteria_ Candidatus_Spechtbacteria |
| *A. clathrodes*: Aug. 2016 vs. Oct. 2018 | 110.478 | -1.818 | 0.406 | -4.484 | 0.000 | Proteobacteria_ HOC36 |
| *A. clathrodes*: Aug. 2016 vs. Oct. 2018 | 73.933 | -2.043 | 0.386 | -5.295 | 0.000 | Chloroflexi_ Caldilineaceae |
| *A. clathrodes*: Aug. 2016 vs. Oct. 2018 | 87.269 | -2.715 | 0.640 | -4.246 | 0.000 | Proteobacteria_ Arenicellaceae |
| *A. clathrodes*: Aug. 2016 vs. Oct. 2018 | 459.748 | -3.131 | 0.402 | -7.791 | 0.000 | Dadabacteria_ Dadabacteriales |
| *A. clathrodes*: Aug. 2016 vs. Oct. 2018 | 1.193 | -3.899 | 1.352 | -2.885 | 0.013 | Actinobacteriota_ Coriobacteriaceae |
| *A. clathrodes*: Aug. 2016 vs. Oct. 2018 | 38.565 | -4.155 | 0.812 | -5.118 | 0.000 | Nanoarchaeota_ Woesearchaeales |
| *A. clathrodes*: Aug. 2016 vs. Oct. 2018 | 1.032 | -4.731 | 1.599 | -2.958 | 0.011 | Campilobacterota_ Helicobacteraceae |
| *A. clathrodes*: Aug. 2016 vs. Oct. 2018 | 1.668 | -4.793 | 1.244 | -3.852 | 0.001 | Firmicutes_ Veillonellaceae |
| *A. clathrodes*: Aug. 2016 vs. Oct. 2018 | 3.356 | -5.428 | 1.467 | -3.700 | 0.001 | Actinobacteriota_ Bifidobacteriaceae |
| *A. clathrodes*: Aug. 2016 vs. Oct. 2018 | 3.375 | -5.491 | 1.231 | -4.461 | 0.000 | Proteobacteria_ Sutterellaceae |
| *A. clathrodes*: Aug. 2016 vs. Oct. 2018 | 6.784 | -6.130 | 0.930 | -6.591 | 0.000 | Bacteroidota_ Prevotellaceae |
| *A. clathrodes*: Aug. 2016 vs. Oct. 2018 | 51.498 | -6.513 | 0.472 | -13.789 | 0.000 | Bacteroidota_ Bacteroidaceae |
| *A. clathrodes*: Aug. 2016 vs. Oct. 2018 | 7.619 | -6.655 | 0.931 | -7.146 | 0.000 | Firmicutes_ Enterococcaceae |
| *A. clathrodes*: Oct. 2017 vs. Oct. 2018 | 19.536 | 4.595 | 0.997 | 4.610 | 0.000 | Proteobacteria_ Stappiaceae |
| *A. clathrodes*: Oct. 2017 vs. Oct. 2018 | 3497.324 | 2.911 | 0.394 | 7.385 | 0.000 | Acidobacteriota_ Vicinamibacteraceae |
| *A. clathrodes*: Oct. 2017 vs. Oct. 2018 | 63.891 | 2.863 | 0.518 | 5.529 | 0.000 | Planctomycetota_ Pirellulaceae |
| *A. clathrodes*: Oct. 2017 vs. Oct. 2018 | 324.440 | 1.648 | 0.358 | 4.609 | 0.000 | Chloroflexi_ JG30-KF-CM66 |
| *A. clathrodes*: Oct. 2017 vs. Oct. 2018 | 5070.997 | 1.541 | 0.376 | 4.093 | 0.000 | Actinobacteriota_ Microtrichaceae |
| *A. clathrodes*: Oct. 2017 vs. Oct. 2018 | 139.306 | 1.138 | 0.419 | 2.713 | 0.031 | Cyanobacteria_ Cyanobiaceae |
| *A. clathrodes*: Oct. 2017 vs. Oct. 2018 | 58.177 | -1.131 | 0.391 | -2.895 | 0.019 | Poribacteria_ Poribacteria |
| *A. clathrodes*: Oct. 2017 vs. Oct. 2018 | 971.763 | -1.160 | 0.287 | -4.045 | 0.000 | Bdellovibrionota_ Bdellovibrionaceae |
| *A. clathrodes*: Oct. 2017 vs. Oct. 2018 | 828.027 | -1.273 | 0.437 | -2.914 | 0.019 | Acidobacteriota_ PAUC26f |
| *A. clathrodes*: Oct. 2017 vs. Oct. 2018 | 123.395 | -1.483 | 0.429 | -3.456 | 0.004 | PAUC34f_ PAUC34f |

| | | | | | | |
|---|---|---|---|---|---|---|
| *A. clathrodes*: Oct. 2017 vs. Oct. 2018 | 387.437 | -1.668 | 0.413 | -4.040 | 0.000 | Nitrospirota_ Nitrospiraceae |
| *A. clathrodes*: Oct. 2017 vs. Oct. 2018 | 9.009 | -1.772 | 0.616 | -2.875 | 0.020 | Spirochaetota_ Leptospiraceae |
| *A. clathrodes*: Oct. 2017 vs. Oct. 2018 | 87.269 | -1.882 | 0.595 | -3.164 | 0.010 | Proteobacteria_ Arenicellaceae |
| *A. clathrodes*: Oct. 2017 vs. Oct. 2018 | 73.933 | -2.142 | 0.366 | -5.860 | 0.000 | Chloroflexi_ Caldilineaceae |
| *A. clathrodes*: Oct. 2017 vs. Oct. 2018 | 3.478 | -2.793 | 0.934 | -2.991 | 0.016 | Proteobacteria_ Ectothiorhodospiraceae |
| *A. clathrodes*: Oct. 2017 vs. Oct. 2018 | 459.748 | -2.887 | 0.374 | -7.721 | 0.000 | Dadabacteria_ Dadabacteriales |
| *A. clathrodes*: Oct. 2017 vs. Oct. 2018 | 5.366 | -2.985 | 0.737 | -4.049 | 0.000 | Actinobacteriota_ Actinomarinaceae |
| *A. clathrodes*: Oct. 2017 vs. Oct. 2018 | 1.440 | -2.995 | 0.990 | -3.026 | 0.015 | Proteobacteria_ Clade_IV |
| *A. clathrodes*: Oct. 2017 vs. Oct. 2018 | 14.297 | -3.049 | 0.550 | -5.540 | 0.000 | Proteobacteria_ SAR86_clade |
| *A. clathrodes*: Oct. 2017 vs. Oct. 2018 | 38.565 | -3.362 | 0.762 | -4.409 | 0.000 | Nanoarchaeota_ Woesearchaeales |
| *A. clathrodes*: Oct. 2017 vs. Oct. 2018 | 17.357 | -3.418 | 0.501 | -6.820 | 0.000 | Bacteroidota_ Flavobacteriaceae |
| *A. clathrodes*: Oct. 2017 vs. Oct. 2018 | 5.892 | -3.539 | 0.913 | -3.877 | 0.001 | Proteobacteria_ Clade_II |
| *A. clathrodes*: Oct. 2017 vs. Oct. 2018 | 8.131 | -4.234 | 0.657 | -6.447 | 0.000 | Proteobacteria_ AEGEAN-169_marine_group |
| *A. clathrodes*: Oct. 2017 vs. Oct. 2018 | 15.669 | -4.311 | 0.715 | -6.032 | 0.000 | Proteobacteria_ Clade_I |
| *A. clathrodes*: Oct. 2017 vs. Oct. 2018 | 24.841 | -6.160 | 1.282 | -4.807 | 0.000 | Nanoarchaeota_ SCGC_AAA011-D5 |
| Seawater: Aug. 2016 vs. Oct. 2018 | 22.204 | 19.242 | 2.660 | 7.233 | 0.000 | Proteobacteria_ Burkholderiaceae |
| Seawater: Aug. 2016 vs. Oct. 2018 | 11.752 | 8.276 | 1.335 | 6.198 | 0.000 | Bacteroidota_ Balneolaceae |
| Seawater: Aug. 2016 vs. Oct. 2018 | 10.880 | 4.904 | 1.305 | 3.757 | 0.001 | Verrucomicrobiota_ Kiritimatiellaceae |
| Seawater: Aug. 2016 vs. Oct. 2018 | 4.309 | 4.231 | 1.361 | 3.110 | 0.008 | Bacteroidota_ Saprospiraceae |
| Seawater: Aug. 2016 vs. Oct. 2018 | 236.318 | 3.866 | 1.069 | 3.615 | 0.002 | Proteobacteria_ Pseudomonadaceae |
| Seawater: Aug. 2016 vs. Oct. 2018 | 4.822 | 3.745 | 0.962 | 3.892 | 0.001 | Nanoarchaeota_ Woesearchaeales |
| Seawater: Aug. 2016 vs. Oct. 2018 | 27.381 | 3.662 | 1.163 | 3.148 | 0.007 | Proteobacteria_ PS1_clade |
| Seawater: Aug. 2016 vs. Oct. 2018 | 105.108 | 3.497 | 0.547 | 6.395 | 0.000 | Proteobacteria_ Clade_III |
| Seawater: Aug. 2016 vs. Oct. 2018 | 37.091 | 3.087 | 0.626 | 4.934 | 0.000 | Proteobacteria_ Porticoccaceae |
| Seawater: Aug. 2016 vs. Oct. 2018 | 72.446 | 3.033 | 0.620 | 4.893 | 0.000 | Proteobacteria_ Litoricolaceae |
| Seawater: Aug. 2016 vs. Oct. 2018 | 99.230 | 2.621 | 0.894 | 2.930 | 0.014 | Proteobacteria_ Sphingomonadaceae |
| Seawater: Aug. 2016 vs. Oct. 2018 | 23.010 | 2.341 | 0.940 | 2.492 | 0.042 | Proteobacteria_ Parvibaculaceae |
| Seawater: Aug. 2016 vs. Oct. 2018 | 50.663 | 2.198 | 0.486 | 4.521 | 0.000 | Bacteroidota_ Cryomorphaceae |
| Seawater: Aug. 2016 vs. Oct. 2018 | 64.932 | 2.005 | 0.435 | 4.608 | 0.000 | Proteobacteria_ MWH-UniP1_aquatic_group |
| Seawater: Aug. 2016 vs. Oct. 2018 | 773.802 | 1.661 | 0.343 | 4.845 | 0.000 | Actinobacteriota_ Actinomarinaceae |
| Seawater: Aug. 2016 vs. Oct. 2018 | 628.467 | 1.611 | 0.299 | 5.386 | 0.000 | Proteobacteria_ SAR116_clade |
| Seawater: Aug. 2016 vs. Oct. 2018 | 113.247 | 1.595 | 0.367 | 4.347 | 0.000 | Verrucomicrobiota_ Puniceicoccaceae |
| Seawater: Aug. 2016 vs. Oct. 2018 | 1037.397 | 1.503 | 0.304 | 4.936 | 0.000 | Proteobacteria_ AEGEAN-169_marine_group |
| Seawater: Aug. 2016 vs. Oct. 2018 | 673.981 | 1.407 | 0.288 | 4.884 | 0.000 | Bacteroidota_ Flavobacteriaceae |
| Seawater: Aug. 2016 vs. Oct. 2018 | 2036.281 | 1.330 | 0.289 | 4.602 | 0.000 | Proteobacteria_ SAR86_clade |
| Seawater: Aug. 2016 vs. Oct. 2018 | 26.126 | 1.285 | 0.408 | 3.151 | 0.007 | Bacteroidota_ NS7_marine_group |
| Seawater: Aug. 2016 vs. Oct. 2018 | 190.611 | 1.105 | 0.329 | 3.362 | 0.004 | Proteobacteria_ Halieaceae |
| Seawater: Aug. 2016 vs. Oct. 2018 | 43.330 | 1.102 | 0.412 | 2.676 | 0.029 | Planctomycetota_ Phycisphaeraceae |
| Seawater: Aug. 2016 vs. Oct. 2018 | 649.623 | 0.613 | 0.244 | 2.512 | 0.042 | Proteobacteria_ Rhodobacteraceae |
| Seawater: Aug. 2016 vs. Oct. 2018 | 157.819 | -0.946 | 0.276 | -3.428 | 0.003 | Proteobacteria_ Pseudohongiellaceae |

| | | | | | | |
|---|---|---|---|---|---|---|
| Seawater: Aug. 2016 vs. Oct. 2018 | 108.200 | -0.954 | 0.372 | -2.566 | 0.037 | Planctomycetota_ Pirellulaceae |
| Seawater: Aug. 2016 vs. Oct. 2018 | 86.848 | -1.095 | 0.439 | -2.495 | 0.042 | Actinobacteriota_ Microtrichaceae |
| Seawater: Aug. 2016 vs. Oct. 2018 | 28.258 | -1.322 | 0.495 | -2.671 | 0.029 | Proteobacteria_ Nitrosomonadaceae |
| Seawater: Aug. 2016 vs. Oct. 2018 | 17.546 | -1.464 | 0.552 | -2.652 | 0.029 | Planctomycetota_ Pla3_lineage |
| Seawater: Aug. 2016 vs. Oct. 2018 | 133.510 | -1.565 | 0.327 | -4.789 | 0.000 | Proteobacteria_ OM182_clade |
| Seawater: Aug. 2016 vs. Oct. 2018 | 115.980 | -1.789 | 0.407 | -4.393 | 0.000 | Dadabacteria_ Dadabacteriales |
| Seawater: Aug. 2016 vs. Oct. 2018 | 226.003 | -2.061 | 0.428 | -4.820 | 0.000 | Chloroflexi_ SAR202_clade |
| Seawater: Aug. 2016 vs. Oct. 2018 | 92.985 | -2.069 | 0.490 | -4.220 | 0.000 | Proteobacteria_ OCS116_clade |
| Seawater: Aug. 2016 vs. Oct. 2018 | 74.891 | -2.187 | 0.334 | -6.552 | 0.000 | Verrucomicrobiota_ Pedosphaeraceae |
| Seawater: Aug. 2016 vs. Oct. 2018 | 29.663 | -2.768 | 0.951 | -2.910 | 0.014 | Proteobacteria_ UBA10353_marine_group |
| Seawater: Aug. 2016 vs. Oct. 2018 | 86.442 | -3.018 | 0.693 | -4.353 | 0.000 | Proteobacteria_ HOC36 |
| Seawater: Aug. 2016 vs. Oct. 2018 | 19.411 | -4.529 | 1.456 | -3.110 | 0.008 | Proteobacteria_ Nitrosococcaceae |
| Seawater: Aug. 2016 vs. Oct. 2018 | 144.855 | -4.727 | 0.401 | -11.780 | 0.000 | Verrucomicrobiota_ DEV007 |
| Seawater: Aug. 2016 vs. Oct. 2018 | 74.929 | -5.150 | 1.309 | -3.935 | 0.001 | Proteobacteria_ Pseudoalteromonadaceae |
| Seawater: Aug. 2016 vs. Oct. 2018 | 115.602 | -5.322 | 0.428 | -12.429 | 0.000 | Verrucomicrobiota_ Arctic97B-4_marine_group |
| Seawater: Aug. 2016 vs. Oct. 2018 | 8.152 | -5.908 | 1.765 | -3.348 | 0.004 | Planctomycetota_ Gimesiaceae |
| Seawater: Aug. 2016 vs. Oct. 2018 | 12.945 | -6.708 | 1.347 | -4.980 | 0.000 | Planctomycetota_ Rubinisphaeraceae |
| Seawater: Aug. 2016 vs. Oct. 2018 | 137.674 | -7.707 | 1.854 | -4.158 | 0.000 | Proteobacteria_ Saccharospirillaceae |
| Seawater: Aug. 2016 vs. Oct. 2018 | 13.166 | -9.317 | 2.787 | -3.343 | 0.004 | Proteobacteria_ Colwelliaceae |
| Seawater: Oct. 2017 vs. Oct. 2018 | 22.204 | 17.944 | 1.953 | 9.186 | 0.000 | Proteobacteria_ Burkholderiaceae |
| Seawater: Oct. 2017 vs. Oct. 2018 | 20.739 | 14.968 | 3.046 | 4.914 | 0.000 | Bacteroidota_ Chitinophagaceae |
| Seawater: Oct. 2017 vs. Oct. 2018 | 32.122 | 4.320 | 1.059 | 4.078 | 0.001 | Verrucomicrobiota_ Rubritaleaceae |
| Seawater: Oct. 2017 vs. Oct. 2018 | 236.318 | 4.054 | 0.799 | 5.076 | 0.000 | Proteobacteria_ Pseudomonadaceae |
| Seawater: Oct. 2017 vs. Oct. 2018 | 4.822 | 4.014 | 0.809 | 4.964 | 0.000 | Nanoarchaeota_ Woesearchaeales |
| Seawater: Oct. 2017 vs. Oct. 2018 | 11.752 | 3.960 | 1.056 | 3.751 | 0.002 | Bacteroidota_ Balneolaceae |
| Seawater: Oct. 2017 vs. Oct. 2018 | 4.160 | 3.949 | 1.010 | 3.910 | 0.001 | Desulfobacterota_ Bradymonadales |
| Seawater: Oct. 2017 vs. Oct. 2018 | 99.230 | 2.788 | 0.649 | 4.295 | 0.000 | Proteobacteria_ Sphingomonadaceae |
| Seawater: Oct. 2017 vs. Oct. 2018 | 10.880 | 2.768 | 0.985 | 2.810 | 0.039 | Verrucomicrobiota_ Kiritimatiellaceae |
| Seawater: Oct. 2017 vs. Oct. 2018 | 65.490 | 2.002 | 0.715 | 2.800 | 0.039 | Crenarchaeota_ Nitrosopumilaceae |
| Seawater: Oct. 2017 vs. Oct. 2018 | 311.983 | 0.942 | 0.348 | 2.708 | 0.049 | Thermoplasmatota_ Marine_Group_II |
| Seawater: Oct. 2017 vs. Oct. 2018 | 1394.069 | -0.906 | 0.241 | -3.755 | 0.002 | Proteobacteria_ Clade_II |
| Seawater: Oct. 2017 vs. Oct. 2018 | 3348.637 | -1.193 | 0.220 | -5.421 | 0.000 | Proteobacteria_ Clade_I |
| Seawater: Oct. 2017 vs. Oct. 2018 | 143.387 | -1.825 | 0.617 | -2.957 | 0.027 | Proteobacteria_ Vibrionaceae |
| Seawater: Oct. 2017 vs. Oct. 2018 | 13.166 | -8.856 | 1.947 | -4.550 | 0.000 | Proteobacteria_ Colwelliaceae |
| Seawater: Oct. 2017 vs. Oct. 2018 | 66.304 | -14.304 | 3.903 | -3.665 | 0.002 | Cyanobacteria_ Obscuribacteraceae |
| Seawater: Reef Oct. 2017 vs. Oct. 2018 | 66.304 | 24.353 | 3.788 | 6.428 | 0.000 | Cyanobacteria_ Obscuribacteraceae |
| Seawater: Reef Oct. 2017 vs. Oct. 2018 | 20.739 | 22.334 | 2.950 | 7.569 | 0.000 | Bacteroidota_ Chitinophagaceae |
| Seawater: Reef Oct. 2017 vs. Oct. 2018 | 9.306 | 21.897 | 3.789 | 5.779 | 0.000 | Bacteroidota_ Cytophagaceae |
| Seawater: Reef Oct. 2017 vs. Oct. 2018 | 22.204 | 21.866 | 1.916 | 11.415 | 0.000 | Proteobacteria_ Burkholderiaceae |
| Seawater: Reef Oct. 2017 vs. Oct. 2018 | 41.297 | 8.594 | 1.799 | 4.777 | 0.000 | Proteobacteria_ Caulobacteraceae |

| | | | | | | |
|---|---|---|---|---|---|---|
| Seawater: Reef Oct. 2017 vs. Oct. 2018 | 236.318 | 7.780 | 0.785 | 9.907 | 0.000 | Proteobacteria_ Pseudomonadaceae |
| Seawater: Reef Oct. 2017 vs. Oct. 2018 | 14.235 | 7.054 | 2.136 | 3.302 | 0.004 | Proteobacteria_ Kordiimonadaceae |
| Seawater: Reef Oct. 2017 vs. Oct. 2018 | 13.908 | 7.016 | 1.950 | 3.598 | 0.002 | Proteobacteria_ Beijerinckiaceae |
| Seawater: Reef Oct. 2017 vs. Oct. 2018 | 6.511 | 5.646 | 1.522 | 3.710 | 0.001 | Proteobacteria_ Endozoicomonadaceae |
| Seawater: Reef Oct. 2017 vs. Oct. 2018 | 5.192 | 5.399 | 1.222 | 4.418 | 0.000 | Proteobacteria_ Moraxellaceae |
| Seawater: Reef Oct. 2017 vs. Oct. 2018 | 8.434 | 5.254 | 2.075 | 2.532 | 0.037 | Proteobacteria_ Comamonadaceae |
| Seawater: Reef Oct. 2017 vs. Oct. 2018 | 4.822 | 3.631 | 0.819 | 4.431 | 0.000 | Nanoarchaeota_ Woesearchaeales |
| Seawater: Reef Oct. 2017 vs. Oct. 2018 | 99.230 | 3.418 | 0.638 | 5.355 | 0.000 | Proteobacteria_ Sphingomonadaceae |
| Seawater: Reef Oct. 2017 vs. Oct. 2018 | 8.399 | 3.327 | 1.250 | 2.663 | 0.027 | Proteobacteria_ Cellvibrionaceae |
| Seawater: Reef Oct. 2017 vs. Oct. 2018 | 65.490 | 2.080 | 0.703 | 2.957 | 0.013 | Crenarchaeota_ Nitrosopumilaceae |
| Seawater: Reef Oct. 2017 vs. Oct. 2018 | 278.308 | 1.735 | 0.626 | 2.772 | 0.020 | Proteobacteria_ Alteromonadaceae |
| Seawater: Reef Oct. 2017 vs. Oct. 2018 | 84.241 | 0.969 | 0.320 | 3.032 | 0.010 | Bacteroidota_ Cyclobacteriaceae |
| Seawater: Reef Oct. 2017 vs. Oct. 2018 | 199.180 | 0.704 | 0.208 | 3.384 | 0.003 | Proteobacteria_ KI89A_clade |
| Seawater: Reef Oct. 2017 vs. Oct. 2018 | 190.611 | -0.680 | 0.237 | -2.877 | 0.015 | Proteobacteria_ Halieaceae |
| Seawater: Reef Oct. 2017 vs. Oct. 2018 | 773.802 | -0.705 | 0.242 | -2.909 | 0.014 | Actinobacteriota_ Actinomarinaceae |
| Seawater: Reef Oct. 2017 vs. Oct. 2018 | 146.913 | -0.722 | 0.249 | -2.906 | 0.014 | Proteobacteria_ Ectothiorhodospiraceae |
| Seawater: Reef Oct. 2017 vs. Oct. 2018 | 228.696 | -0.844 | 0.245 | -3.445 | 0.003 | Proteobacteria_ Clade_IV |
| Seawater: Reef Oct. 2017 vs. Oct. 2018 | 74.891 | -0.855 | 0.237 | -3.604 | 0.002 | Verrucomicrobiota_ Pedosphaeraceae |
| Seawater: Reef Oct. 2017 vs. Oct. 2018 | 3348.637 | -1.046 | 0.216 | -4.842 | 0.000 | Proteobacteria_ Clade_I |
| Seawater: Reef Oct. 2017 vs. Oct. 2018 | 34.925 | -1.111 | 0.326 | -3.411 | 0.003 | Proteobacteria_ Thiotrichaceae |
| Seawater: Reef Oct. 2017 vs. Oct. 2018 | 220.633 | -1.201 | 0.233 | -5.152 | 0.000 | SAR324_clade_Marine group B |
| Seawater: Reef Oct. 2017 vs. Oct. 2018 | 17.546 | -1.264 | 0.408 | -3.099 | 0.009 | Planctomycetota_ Pla3_lineage |
| Seawater: Reef Oct. 2017 vs. Oct. 2018 | 39.138 | -1.303 | 0.504 | -2.584 | 0.033 | Desulfobacterota_ PB19 |
| Seawater: Reef Oct. 2017 vs. Oct. 2018 | 6901.427 | -1.316 | 0.293 | -4.497 | 0.000 | Cyanobacteria_ Cyanobiaceae |
| Seawater: Reef Oct. 2017 vs. Oct. 2018 | 57.481 | -1.520 | 0.362 | -4.206 | 0.000 | Proteobacteria_ Coxiellaceae |
| Seawater: Reef Oct. 2017 vs. Oct. 2018 | 9.987 | -2.242 | 0.539 | -4.160 | 0.000 | Actinobacteriota_ PeM15 |
| Seawater: Reef Oct. 2017 vs. Oct. 2018 | 2.962 | -3.478 | 1.026 | -3.390 | 0.003 | Proteobacteria_ Rickettsiaceae |
| Seawater: Reef Oct. 2017 vs. Oct. 2018 | 8.562 | -3.715 | 0.718 | -5.173 | 0.000 | WPS-2_ WPS-2 |
| Seawater: Reef Oct. 2017 vs. Oct. 2018 | 13.166 | -6.548 | 1.901 | -3.444 | 0.003 | Proteobacteria_ Colwelliaceae |
| Seawater: Reef Oct. 2017 vs. Oct. 2018 | 10.540 | -23.105 | 1.604 | -14.400 | 0.000 | Proteobacteria_ Thioglobaceae |

**Supplemental Table S6.** Bacterial taxa significantly associated with recent flooding events according to Indicator Species Analysis. Analysis was conducted on each sample type (*Xestospongia muta*, *Agelas clathrodes*, and seawater) individually.

| Amplicon Sequence Variant | Group | Phylum_Class_Family_Genus | Indicator Value | P value |
|---|---|---|---|---|
| 1acd92584c994e912f4e31130344f3fb | *X. muta*: Aug. 2016 | Actinobacteriota_ Acidimicrobiia_ Actinomarinaceae_ Candidatus_Actinomarina | 0.541 | 0.0112 |
| 9f44ce941b1f8eb567cee4b400c43356 | *X. muta*: Aug. 2016 | Actinobacteriota_ Acidimicrobiia_ Microtrichaceae_ Sva0996 marine group | 0.463 | 0.0343 |
| a6bb187d647af488dfd1ec39db511f4d | *X. muta*: Aug. 2016 | Actinobacteriota_ Acidimicrobiia_ uncultured_ uncultured | 0.496 | 0.0283 |
| 59185fd96fab2c505f36dcbd50c52906 | *X. muta*: Aug. 2016 | Actinobacteriota_ Acidimicrobiia_ uncultured_ uncultured | 0.496 | 0.0029 |
| ccd8d5e31ad9bfa58066251ea1138225 | *X. muta*: Aug. 2016 | Bacteroidota_ Bacteroidia_ Flavobacteriaceae_unclassified | 0.577 | 0.0277 |
| 64805be33440e427c18e31e0d5e6094b | *X. muta*: Aug. 2016 | Bacteroidota_ Bacteroidia_ Saprospiraceae_ Phaeodactylibacter | 0.577 | 0.0277 |
| 8279d4ca0b7ad8d79bfd2f42749efd25 | *X. muta*: Aug. 2016 | Bdellovibrionota_ Bdellovibrionia_ Bacteriovoracaceae_ Peredibacter | 0.448 | 0.0458 |
| e7ae46f8a3f261e09db98662c33d66ef | *X. muta*: Aug. 2016 | Bdellovibrionota_ Bdellovibrionia_ Bdellovibrionaceae_ Bdellovibrio | 0.577 | 0.0277 |
| 94c6b9401c804746ce724ae2678419a9 | *X. muta*: Aug. 2016 | Bdellovibrionota_ Bdellovibrionia_ Bdellovibrionaceae_ Bdellovibrio | 0.545 | 0.0284 |
| e7a93cbcaecd06373372a13718cf957f | *X. muta*: Aug. 2016 | Bdellovibrionota_ Bdellovibrionia_ Bdellovibrionaceae_ Bdellovibrio | 0.54 | 0.0063 |
| 4c49be929ea5defb5a681abdb3ef13c5 | *X. muta*: Aug. 2016 | Bdellovibrionota_ Bdellovibrionia_ Bdellovibrionaceae_ Bdellovibrio | 0.536 | 0.0133 |
| 58f20f50d118fab8b8c501d59f0eb40d | *X. muta*: Aug. 2016 | Bdellovibrionota_ Bdellovibrionia_ Bdellovibrionaceae_ Bdellovibrio | 0.509 | 0.0165 |
| 26f9913a22bb3c8927c40399d23c23b3 | *X. muta*: Aug. 2016 | Bdellovibrionota_ Bdellovibrionia_ Bdellovibrionaceae_ Bdellovibrio | 0.499 | 0.0262 |
| 3cd16664a28ac80726958a35bf176dfe | *X. muta*: Aug. 2016 | Bdellovibrionota_ Bdellovibrionia_ Bdellovibrionaceae_ Bdellovibrio | 0.488 | 0.0284 |
| 5ebff532b435d2db1adc35f15dfc4ea8 | *X. muta*: Aug. 2016 | Bdellovibrionota_ Bdellovibrionia_ Bdellovibrionaceae_ Bdellovibrio | 0.473 | 0.0038 |
| e976f07cf2723e5b7dc5640e27705210 | *X. muta*: Aug. 2016 | Bdellovibrionota_ Bdellovibrionia_ Bdellovibrionaceae_ Bdellovibrio | 0.472 | 0.0493 |
| 0adc5a9aa8b36ed37177f9ad6f1846db | *X. muta*: Aug. 2016 | Bdellovibrionota_ Bdellovibrionia_ Bdellovibrionaceae_ Bdellovibrio | 0.467 | 0.0333 |
| 7a18ffa05f11051d0f02e39fa6de5709 | *X. muta*: Aug. 2016 | Bdellovibrionota_ Oligoflexia_ uncultured_ uncultured | 0.525 | 0.0009 |
| 2990fe8b204d3cdc8f107ef1a8f9661c | *X. muta*: Aug. 2016 | Bdellovibrionota_ Oligoflexia_ uncultured_ uncultured | 0.474 | 0.0381 |
| 00a0baccdfc8873dd173a84e8b670e01 | *X. muta*: Aug. 2016 | Bdellovibrionota_ Oligoflexia_ uncultured_ uncultured | 0.424 | 0.0337 |
| 58f380b3f2a399f6c26ba2c5b5c1b61e | *X. muta*: Aug. 2016 | Cyanobacteria_ Cyanobacteriia_ Cyanobiaceae_ Prochlorococcus_MIT9313 | 0.548 | 0.0248 |
| cf53a56718dec263aa9ce290bdd15404 | *X. muta*: Aug. 2016 | Cyanobacteria_ Cyanobacteriia_ Cyanobiaceae_ Synechococcus | 0.57 | 0.0036 |
| f7d1d099f965377b0d20486ca77b677e | *X. muta*: Aug. 2016 | Cyanobacteria_ Cyanobacteriia_ Cyanobiaceae_ Synechococcus | 0.547 | 0.0263 |
| e09f2d58d292332880fe9e37607469e1 | *X. muta*: Aug. 2016 | Desulfobacterota_ Desulfovibrionia_ Desulfovibrionaceae_ Halodesulfovibrio | 0.573 | 0.0277 |
| e4a5985be454d1e8a3d50a2ab6ea7a8b | *X. muta*: Aug. 2016 | Desulfobacterota_ Desulfovibrionia_ Desulfovibrionaceae_ Halodesulfovibrio | 0.525 | 0.0027 |
| e51947121c608cf15031596f9d0c09eb | *X. muta*: Aug. 2016 | Desulfobacterota_ Desulfovibrionia_ Desulfovibrionaceae_NA | 0.461 | 0.0277 |
| 0261a50c7edb68abf5e65bb5094ce011 | *X. muta*: Aug. 2016 | Desulfobacterota_ Desulfovibrionia_ Desulfovibrionaceae_NA | 0.426 | 0.0277 |
| 385897234481e89b450a320371bb9c18 | *X. muta*: Aug. 2016 | Firmicutes_ Clostridia_ Fusibacteraceae_ Fusibacter | 0.571 | 0.0277 |
| a13fd587face977324024ea2f0c13e93 | *X. muta*: Aug. 2016 | Firmicutes_ Clostridia_ Peptostreptococcales-Tissierellales_ Anaeromicrobium | 0.565 | 0.0277 |
| 219ab3d6aaec11caf9e19b7652251c7b | *X. muta*: Aug. 2016 | Myxococcota_ bacteriap25_ bacteriap25_ bacteriap25 | 0.434 | 0.0362 |
| 7acd4f4ee4d657cf1cbff5b3f5873793 | *X. muta*: Aug. 2016 | Nanoarchaeota_ Nanoarchaeia_ Woesearchaeales_ Woesearchaeales | 0.565 | 0.0263 |
| 11e5775c3c621cd5e49485b9312c68e4 | *X. muta*: Aug. 2016 | Nitrospirota_ Nitrospiria_ Nitrospiraceae_ Nitrospira | 0.454 | 0.0388 |
| 35a8ee0f6c51051f654259a84b009c40 | *X. muta*: Aug. 2016 | Patescibacteria_ Saccharimonadia_ Saccharimonadales_ Saccharimonadales | 0.724 | 0.0015 |
| 989dc9271f6623ac4a4317ebe6fe3d96 | *X. muta*: Aug. 2016 | Patescibacteria_ Saccharimonadia_ Saccharimonadales_ Saccharimonadales | 0.526 | 0.0278 |

| | | | | |
|---|---|---|---|---|
| 60ccb7eac903709aae8dbc4e06870bf3 | *X. muta*: Aug. 2016 | Patescibacteria_ Saccharimonadia_ Saccharimonadales_ Saccharimonadales | 0.497 | 0.0017 |
| e03a0122cd5e09b64f02caa03c5d118b | *X. muta*: Aug. 2016 | PAUC34f_ PAUC34f_ PAUC34f_ PAUC34f | 0.507 | 0.0251 |
| 2a8c49a3dc619404d2cc309b796e44d4 | *X. muta*: Aug. 2016 | Poribacteria_ Poribacteria_ Poribacteria_ Poribacteria | 0.593 | 0.0067 |
| 74e8846e250814bf5418bb63ffed7adf | *X. muta*: Aug. 2016 | Poribacteria_ Poribacteria_ Poribacteria_ Poribacteria | 0.544 | 0.0047 |
| d246072d506010a5f472e70c7f8ac6c3 | *X. muta*: Aug. 2016 | Poribacteria_ Poribacteria_ Poribacteria_ Poribacteria | 0.517 | 0.0322 |
| f082fd9ffc0c39f7ffd27b093648407b | *X. muta*: Aug. 2016 | Poribacteria_ Poribacteria_ Poribacteria_ Poribacteria | 0.517 | 0.0328 |
| 6e08db9fa0d10596303c6f3a7564ee0c | *X. muta*: Aug. 2016 | Proteobacteria_ Alphaproteobacteria_ AEGEAN-169 marine group | 0.657 | 0.0026 |
| 5546361caf3e42e50fb56d85a11571f2 | *X. muta*: Aug. 2016 | Proteobacteria_ Alphaproteobacteria_ Clade_I_ Clade_Ia | 0.533 | 0.0248 |
| 5742262f41c8464b0518363de93692c8 | *X. muta*: Aug. 2016 | Proteobacteria_ Alphaproteobacteria_ Clade_IV_ Clade_IV | 0.682 | 0.0016 |
| 487f16e5da93cf159909d2d497fad3c5 | *X. muta*: Aug. 2016 | Proteobacteria_ Alphaproteobacteria_ Nisaeaceae_ OM75_clade | 0.491 | 0.0458 |
| 8b72c63a681aa6efff8e7e9fa0d4f16d | *X. muta*: Aug. 2016 | Proteobacteria_ Alphaproteobacteria_ Rhodobacteraceae_ Albidovulum | 0.567 | 0.0054 |
| 48367e547c1ba5167d58994df3c30cee | *X. muta*: Aug. 2016 | Proteobacteria_ Alphaproteobacteria_ Rhodobacteraceae_ uncultured | 0.442 | 0.0458 |
| cc31f1ecc1714e03edfa64901ffb3fe9 | *X. muta*: Aug. 2016 | Proteobacteria_ Alphaproteobacteria_ Rhodobacteraceae_NA | 0.577 | 0.0277 |
| 170f84b791738d735741ed6ee12af63d | *X. muta*: Aug. 2016 | Proteobacteria_ Alphaproteobacteria_ Rhodobacteraceae_NA | 0.457 | 0.0385 |
| e76098f481e1dba8781157c0b3586edf | *X. muta*: Aug. 2016 | Proteobacteria_ Alphaproteobacteria_ SAR116_clade_ SAR116_clade | 0.539 | 0.0238 |
| 8f54f51ec7e1706d4263604cfebfe98a | *X. muta*: Aug. 2016 | Proteobacteria_ Alphaproteobacteria_unclassified | 0.668 | 0.0018 |
| 2ef8937fcfef105310d9aae9f65c08ab | *X. muta*: Aug. 2016 | Proteobacteria_ Gammaproteobacteria_ Kangiellaceae_ Aliikangiella | 0.577 | 0.0277 |
| 5cf6cc6546c9dd87665e09c2702bafb8 | *X. muta*: Aug. 2016 | Proteobacteria_ Gammaproteobacteria_ KI89A_clade_ KI89A_clade | 0.505 | 0.0106 |
| 9442980515ce7f911bb4cc9c372db4e8 | *X. muta*: Aug. 2016 | Proteobacteria_ Gammaproteobacteria_ SAR86_clade_ SAR86_clade | 0.805 | 0.0001 |
| b47b6e5813cfd4b7690b522531f10aa4 | *X. muta*: Aug. 2016 | Proteobacteria_ Gammaproteobacteria_ Thioglobaceae_ SUP05_cluster | 0.826 | 0.0004 |
| 93fe6a3924426bdfeaa6517238bb4259 | *X. muta*: Aug. 2016 | Proteobacteria_ Gammaproteobacteria_ UBA10353 marine group | 0.468 | 0.0323 |
| badb932d6b9077f2d7b9f434f92fc9a3 | *X. muta*: Aug. 2016 | Proteobacteria_ Gammaproteobacteria_unclassified | 0.603 | 0.0071 |
| efa28eb526ec0c3caa54fd35809774e1 | *X. muta*: Aug. 2016 | SAR324 clade_SAR324 clade_SAR324 clade_SAR324 clade | 0.577 | 0.0274 |
| 784ab93ee765a6e776d5f8c92263f4a5 | *X. muta*: Aug. 2016 | SAR324 clade_SAR324 clade_SAR324 clade_SAR324 clade | 0.458 | 0.0284 |
| 014e875bd5c772fba46cece4f4b337f2 | *X. muta*: Sept. 2017 | Acidobacteriota_ Acidobacteriae_ PAUC26f_ PAUC26f | 0.533 | 0.0193 |
| 9d691231b57870cf57a310457ddd2ed6 | *X. muta*: Sept. 2017 | Acidobacteriota_ Acidobacteriae_ PAUC26f_ PAUC26f | 0.493 | 0.0226 |
| c43884aba0b5531ded76e219042cce72 | *X. muta*: Sept. 2017 | Acidobacteriota_ Acidobacteriae_ PAUC26f_ PAUC26f | 0.461 | 0.0383 |
| 81c525b33c2ce888506e4b77f0d9a41b | *X. muta*: Sept. 2017 | Acidobacteriota_ Acidobacteriae_NA_NA | 0.602 | 0.005 |
| 3363aef159cc6a36966b2e52418251e8 | *X. muta*: Sept. 2017 | Acidobacteriota_ Subgroup_21_ Subgroup_21_ Subgroup_21 | 0.491 | 0.0294 |
| 6cfe8609506fd1f5edf954903b55551d | *X. muta*: Sept. 2017 | Acidobacteriota_ Subgroup_21_ Subgroup_21_ Subgroup_21 | 0.48 | 0.0343 |
| e3c416584af267707c698bdbfe611c20 | *X. muta*: Sept. 2017 | Acidobacteriota_ Thermoanaerobaculia_ Thermoanaerobaculaceae_ Subgroup 10 | 0.594 | 0.0088 |
| 4097f89849f5fd6e69f89fd0467adb0e | *X. muta*: Sept. 2017 | Acidobacteriota_ Vicinamibacteria_ Subgroup_9_ Subgroup_9 | 0.541 | 0.0239 |
| 9d73d140a47350aae5f29e1fd20f98c4 | *X. muta*: Sept. 2017 | Acidobacteriota_ Vicinamibacteria_ uncultured_ uncultured | 0.574 | 0.0079 |
| 0ed938a1bb9f3ebce43a415afbf39ec1 | *X. muta*: Sept. 2017 | Acidobacteriota_ Vicinamibacteria_ uncultured_ uncultured | 0.502 | 0.0477 |
| aee4afb4738105b99eeae332b7824ceb | *X. muta*: Sept. 2017 | Acidobacteriota_ Vicinamibacteria_ uncultured_ uncultured | 0.471 | 0.0291 |
| 282fc2fabb1351ef2d4caa7047b8aa2c | *X. muta*: Sept. 2017 | Acidobacteriota_ Vicinamibacteria_ uncultured_ uncultured | 0.404 | 0.0304 |
| 5781ed23c9092dce09696c795c5db8a4 | *X. muta*: Sept. 2017 | Acidobacteriota_ Vicinamibacteria_ Vicinamibacteraceae_ Vicinamibacteraceae | 0.459 | 0.0416 |
| 160f9a311cbd59996a90505016861cf6 | *X. muta*: Sept. 2017 | Acidobacteriota_ Vicinamibacteria_NA_NA | 0.472 | 0.0373 |
| ae72fc3614c20c4d97cfc1e8c6bf6912 | *X. muta*: Sept. 2017 | Actinobacteriota_ Acidimicrobiia_ Microtrichaceae_ Sva0996 marine group | 0.555 | 0.0071 |
| 335500a6b88e7a2d97422865b1c74ddf | *X. muta*: Sept. 2017 | Actinobacteriota_ Acidimicrobiia_ Microtrichaceae_ Sva0996 marine group | 0.53 | 0.0227 |

| | | | | |
|---|---|---|---|---|
| 112c91d6d4715fc60b85254840c0f5e5 | *X. muta*: Sept. 2017 | Actinobacteriota_ Acidimicrobiia_ Microtrichaceae_ Sva0996 marine group | 0.517 | 0.0261 |
| 8dd2f62b5dc4df1f84d2db88e330f566 | *X. muta*: Sept. 2017 | Actinobacteriota_ Acidimicrobiia_ Microtrichaceae_ Sva0996 marine group | 0.501 | 0.032 |
| f582f3075ac4a13a1befa9bed4db3222 | *X. muta*: Sept. 2017 | Actinobacteriota_ Acidimicrobiia_ Microtrichaceae_ Sva0996 marine group | 0.484 | 0.0371 |
| 7b2db438be48f839007b9c6ef2985a2d | *X. muta*: Sept. 2017 | Actinobacteriota_ Acidimicrobiia_ Microtrichaceae_ Sva0996 marine group | 0.479 | 0.0297 |
| 9a6f6600708c4e75da1181232be0bd15 | *X. muta*: Sept. 2017 | Actinobacteriota_ Acidimicrobiia_ Microtrichaceae_ Sva0996 marine group | 0.45 | 0.0294 |
| 92b48363c4b512d5d5407751081fe712 | *X. muta*: Sept. 2017 | Actinobacteriota_ Thermoleophilia_ uncultured_ uncultured | 0.477 | 0.0415 |
| a97998beb91d5a43b2fc5aad67157beb | *X. muta*: Sept. 2017 | AncK6_ AncK6_ AncK6_ AncK6 | 0.646 | 0.0018 |
| 549691292795bebf5a0e8b551b9d7430 | *X. muta*: Sept. 2017 | Bacteroidota_ Bacteroidia_ Bacteroidaceae_ Bacteroides | 0.513 | 0.032 |
| 0fabb9c94ddc0c2e5376d2fd8a0354f1 | *X. muta*: Sept. 2017 | Bacteroidota_ Bacteroidia_ Muribaculaceae_ Muribaculaceae | 0.502 | 0.0338 |
| a980986880b89ac73d0fa0261cf6a7ab | *X. muta*: Sept. 2017 | Bacteroidota_ Rhodothermia_ Rhodothermaceae_ uncultured | 0.597 | 0.0056 |
| c928ed28c488be7cef122207df766c04 | *X. muta*: Sept. 2017 | Bacteroidota_ Rhodothermia_ Rhodothermaceae_ uncultured | 0.595 | 0.0078 |
| 5e4ead8ed2ecbcbd55eb5ab20a702439 | *X. muta*: Sept. 2017 | Bacteroidota_ Rhodothermia_ Rhodothermaceae_ uncultured | 0.578 | 0.0037 |
| 8a484cf27de3a6cfee14152aa2fa3204 | *X. muta*: Sept. 2017 | Bdellovibrionota_ Bdellovibrionia_ Bacteriovoracaceae_ Peredibacter | 0.512 | 0.0442 |
| 28f1b98223a43c7a4cdaa1c7f0bce9db | *X. muta*: Sept. 2017 | Bdellovibrionota_ Bdellovibrionia_ Bdellovibrionaceae_ Bdellovibrio | 0.515 | 0.0261 |
| 69b0ba35942d7d4e8a3348f33bf0eced | *X. muta*: Sept. 2017 | Bdellovibrionota_ Bdellovibrionia_ Bdellovibrionaceae_ Bdellovibrio | 0.501 | 0.0383 |
| fa62df4d7a111de5cd87a7df28587585 | *X. muta*: Sept. 2017 | Bdellovibrionota_ Bdellovibrionia_ Bdellovibrionaceae_ Bdellovibrio | 0.471 | 0.0378 |
| f3d2cd15a3f4767c6df280a6539806c4 | *X. muta*: Sept. 2017 | Bdellovibrionota_ Bdellovibrionia_ Bdellovibrionaceae_ Bdellovibrio | 0.47 | 0.024 |
| 09979f3762e724ed368b41986c64b99e | *X. muta*: Sept. 2017 | Chloroflexi_ Anaerolineae_ A4b_ A4b | 0.6 | 0.0057 |
| 13d7365b2e0161a1d143f3b67072209c | *X. muta*: Sept. 2017 | Chloroflexi_ Anaerolineae_ A4b_ A4b | 0.546 | 0.0073 |
| 3067441789a9beb27957c892af50ede1 | *X. muta*: Sept. 2017 | Chloroflexi_ Anaerolineae_ A4b_ A4b | 0.482 | 0.0483 |
| c75f5969ae2cee2a733fe9eb8da21044 | *X. muta*: Sept. 2017 | Chloroflexi_ Anaerolineae_ A4b_ A4b | 0.476 | 0.0461 |
| 2a095b99a34cecad701279d2dc7fe266 | *X. muta*: Sept. 2017 | Chloroflexi_ Anaerolineae_ A4b_ A4b | 0.462 | 0.0323 |
| 08f73ed0f932b8dd481b518f32b2443d | *X. muta*: Sept. 2017 | Chloroflexi_ Anaerolineae_ A4b_ A4b | 0.424 | 0.0323 |
| 52c36fddbe9cf744fd60c6738fdee72b | *X. muta*: Sept. 2017 | Chloroflexi_ Anaerolineae_ Caldilineaceae_ uncultured | 0.514 | 0.0332 |
| 4878aefab68febb56cdd72b7bd0f8fc8 | *X. muta*: Sept. 2017 | Chloroflexi_ Anaerolineae_ Caldilineaceae_ uncultured | 0.452 | 0.0372 |
| 397fcbf74ed9fcc0a4475a1815f7f0ff | *X. muta*: Sept. 2017 | Chloroflexi_ Dehalococcoidia_ SAR202_clade_ SAR202_clade | 0.67 | 0.0012 |
| deb8db91864fabc97c364f50fba6fe3f | *X. muta*: Sept. 2017 | Chloroflexi_ Dehalococcoidia_ SAR202_clade_ SAR202_clade | 0.53 | 0.0197 |
| d1ff1b0729d649fd01c2669326206c8f | *X. muta*: Sept. 2017 | Chloroflexi_ Dehalococcoidia_ SAR202_clade_ SAR202_clade | 0.497 | 0.0279 |
| 805f69cd81c86031ba5931dd5c9a0779 | *X. muta*: Sept. 2017 | Chloroflexi_ SHA-26_ SHA-26_ SHA-26 | 0.48 | 0.0383 |
| 0014ff58293c37e28d24c0abaf779545 | *X. muta*: Sept. 2017 | Chloroflexi_ TK10_ TK10_ TK10 | 0.523 | 0.0189 |
| a601e7967d2298b7edb7bf39df6664b0 | *X. muta*: Sept. 2017 | Chloroflexi_ TK17_ TK17_ TK17 | 0.523 | 0.0366 |
| 90315e56ff9e8960cc6689d28297e89e | *X. muta*: Sept. 2017 | Chloroflexi_ TK30_ TK30_ TK30 | 0.435 | 0.0323 |
| 6e80c62e7ce870b5d18063130b33907f | *X. muta*: Sept. 2017 | Crenarchaeota_ Nitrososphaeria_ Nitrosopumilaceae_ Nitrosopumilaceae | 0.608 | 0.0051 |
| 88c6d77438b8cad4bfc5ff7df88cc954 | *X. muta*: Sept. 2017 | Crenarchaeota_ Nitrososphaeria_ Nitrosopumilaceae_ Nitrosopumilaceae | 0.54 | 0.0072 |
| 534912e2e65dcb144d132b25b02c7d31 | *X. muta*: Sept. 2017 | Entotheonellaeota_ Entotheonellia_ Entotheonellaceae_ Entotheonellaceae | 0.434 | 0.0368 |
| abc672abd615e98ba7a3a594d1f607ec | *X. muta*: Sept. 2017 | Gemmatimonadota_ BD2-11_terrestrial_group_ NA_NA | 0.617 | 0.0052 |
| 455a0e4095e02b4a46470c532dd9a636 | *X. muta*: Sept. 2017 | Gemmatimonadota_ BD2-11_terrestrial_group_ NA_NA | 0.567 | 0.0055 |
| a69e13f88fc1a6e38bff836884dfd77c | *X. muta*: Sept. 2017 | Gemmatimonadota_ BD2-11_terrestrial_group_ NA_NA | 0.452 | 0.0383 |
| a55bbd8e378775030b1a3a74a01f09c3 | *X. muta*: Sept. 2017 | Gemmatimonadota_ PAUC43f marine benthic group_ NA_NA | 0.489 | 0.0301 |
| 6b935d522f98c7f7eb22fb2f78f1db63 | *X. muta*: Sept. 2017 | Myxococcota_ bacteriap25_ bacteriap25_ bacteriap25 | 0.59 | 0.005 |

| | | | | |
|---|---|---|---|---|
| 1725502fe3b545b5f4dfb2d2b580e0b5 | *X. muta*: Sept. 2017 | Myxococcota_ bacteriap25_ bacteriap25_ bacteriap25 | 0.553 | 0.0049 |
| 98dd20a3d10796d3ea25cf9aa8f9504f | *X. muta*: Sept. 2017 | Myxococcota_ bacteriap25_ bacteriap25_ bacteriap25 | 0.535 | 0.0192 |
| 98cc654e1256a4f046d4a0efb31769fa | *X. muta*: Sept. 2017 | Myxococcota_ bacteriap25_ bacteriap25_ bacteriap25 | 0.52 | 0.0426 |
| ef933f30eb8a4d88d760ff5faaf19140 | *X. muta*: Sept. 2017 | NB1-j_ NB1-j_ NB1-j_ NB1-j | 0.567 | 0.0076 |
| 73d1c97be269e6b73c0ed3fe07337609 | *X. muta*: Sept. 2017 | Nitrospinota_ P9X2b3D02_ P9X2b3D02_ P9X2b3D02 | 0.505 | 0.0252 |
| dd6695e07c0b29f8ee0749562385f394 | *X. muta*: Sept. 2017 | Nitrospirota_ Nitrospiria_ Nitrospiraceae_ Nitrospira | 0.541 | 0.0106 |
| 57697d5405329c1e93dedd819bd9d35f | *X. muta*: Sept. 2017 | PAUC34f_ PAUC34f_ PAUC34f_ PAUC34f | 0.617 | 0.003 |
| da6455ff3227f875191c964275e31478 | *X. muta*: Sept. 2017 | PAUC34f_ PAUC34f_ PAUC34f_ PAUC34f | 0.511 | 0.0325 |
| bd53fdd10e333cc6ed226f6fce2c0bb3 | *X. muta*: Sept. 2017 | PAUC34f_ PAUC34f_ PAUC34f_ PAUC34f | 0.481 | 0.0383 |
| eeea35d4946915cb48c1b25b9d748027 | *X. muta*: Sept. 2017 | PAUC34f_ PAUC34f_ PAUC34f_ PAUC34f | 0.453 | 0.0359 |
| fe018e40b18920445ff698ba7e827063 | *X. muta*: Sept. 2017 | PAUC34f_ PAUC34f_ PAUC34f_ PAUC34f | 0.447 | 0.0287 |
| 4baf314d7c6eecd85968c399a9d6ef5c | *X. muta*: Sept. 2017 | Poribacteria_ Poribacteria_ Poribacteria_ Poribacteria | 0.541 | 0.0199 |
| eecf15cb1bee1053177198b95189052a | *X. muta*: Sept. 2017 | Poribacteria_ Poribacteria_ Poribacteria_ Poribacteria | 0.465 | 0.0314 |
| e1866d63b3d587d463698c9bf80b301a | *X. muta*: Sept. 2017 | Proteobacteria_ Alphaproteobacteria_ Puniceispirillales_ Constrictibacter | 0.581 | 0.0075 |
| 9c534dcb0d57008a3b2c7204aa314a1a | *X. muta*: Sept. 2017 | Proteobacteria_ Alphaproteobacteria_ Rhodobacteraceae_ Ruegeria | 0.419 | 0.0267 |
| f07100fd9407b754b61f4fd661ad0d2d | *X. muta*: Sept. 2017 | Proteobacteria_ Alphaproteobacteria_ Rhodobacteraceae_NA | 0.43 | 0.0323 |
| 32c14ecda29c69ffebf9cbd191592ac0 | *X. muta*: Sept. 2017 | Proteobacteria_ Alphaproteobacteria_ uncultured_ uncultured | 0.625 | 0.0052 |
| c29bc77895d4d52c1fdac0b9045810d8 | *X. muta*: Sept. 2017 | Proteobacteria_ Alphaproteobacteria_ uncultured_ uncultured | 0.437 | 0.042 |
| 9b6d2e5b422b1c31e0a14df9be59aea1 | *X. muta*: Sept. 2017 | Proteobacteria_ Alphaproteobacteria_NA_NA | 0.468 | 0.0297 |
| 6d472a1d79dae690d28607258767704e | *X. muta*: Sept. 2017 | Proteobacteria_ Gammaproteobacteria_ Endozoicomonadaceae_ Endozoicomonas | 0.502 | 0.0281 |
| 3b6cd5238b57b8bb2bfd6215432281b8 | *X. muta*: Sept. 2017 | Proteobacteria_ Gammaproteobacteria_ EPR3968-O8a-Bc78_ EPR3968-O8a-Bc78 | 0.566 | 0.0079 |
| fd26e07a719f37602568f1daf30c3075 | *X. muta*: Sept. 2017 | Proteobacteria_ Gammaproteobacteria_ EPR3968-O8a-Bc78_ EPR3968-O8a-Bc78 | 0.469 | 0.0422 |
| 61dff9e414148a0a0d6438339a6246f6 | *X. muta*: Sept. 2017 | Proteobacteria_ Gammaproteobacteria_ KI89A_clade_ KI89A_clade | 0.608 | 0.0059 |
| 0eb1244f2354bbef303c1b8894de91c1 | *X. muta*: Sept. 2017 | Proteobacteria_ Gammaproteobacteria_ KI89A_clade_ KI89A_clade | 0.604 | 0.0054 |
| a67082f4d3cd06bddc90bae1f9cbee6f | *X. muta*: Sept. 2017 | Proteobacteria_ Gammaproteobacteria_ KI89A_clade_ KI89A_clade | 0.579 | 0.0071 |
| 4654e47a8ad997042e1e2f078ac2f55c | *X. muta*: Sept. 2017 | Proteobacteria_ Gammaproteobacteria_ KI89A_clade_ KI89A_clade | 0.577 | 0.0071 |
| 2f174f9e0dddbf72400f191a24fb63b1 | *X. muta*: Sept. 2017 | Proteobacteria_ Gammaproteobacteria_ KI89A_clade_ KI89A_clade | 0.537 | 0.0104 |
| d5cda0d773b7cf74d504cd602571acdd | *X. muta*: Sept. 2017 | Proteobacteria_ Gammaproteobacteria_ KI89A_clade_ KI89A_clade | 0.518 | 0.0271 |
| e555b814a6265849d108b04a398c5a5d | *X. muta*: Sept. 2017 | Proteobacteria_ Gammaproteobacteria_ KI89A_clade_ KI89A_clade | 0.501 | 0.0343 |
| 2165c5f164dbc4b43d252bc69b8adf67 | *X. muta*: Sept. 2017 | Proteobacteria_ Gammaproteobacteria_ KI89A_clade_ KI89A_clade | 0.491 | 0.0305 |
| f17418555b73d3138ddfd2cd744b21ac | *X. muta*: Sept. 2017 | Proteobacteria_ Gammaproteobacteria_ KI89A_clade_ KI89A_clade | 0.438 | 0.0383 |
| d2a25ec67179a0b6235d806a752deac8 | *X. muta*: Sept. 2017 | Proteobacteria_ Gammaproteobacteria_ Nitrosococcaceae_ AqS1 | 0.566 | 0.0072 |
| 4bf34e939f3102402abec9deab0319b3 | *X. muta*: Sept. 2017 | Proteobacteria_ Gammaproteobacteria_ Nitrosococcaceae_ AqS1 | 0.489 | 0.0359 |
| 413e612426ce719987419494fa632884 | *X. muta*: Sept. 2017 | Proteobacteria_ Gammaproteobacteria_ Nitrosococcaceae_ FS142-36B-02 | 0.508 | 0.0154 |
| d23d37bbf6dc8be13e10a2714adb5d27 | *X. muta*: Sept. 2017 | Proteobacteria_ Gammaproteobacteria_ pItb-vmat-80_ pItb-vmat-80 | 0.49 | 0.0383 |
| f7b4fa51e3a082c099807cb73250d628 | *X. muta*: Sept. 2017 | Proteobacteria_ Gammaproteobacteria_NA_NA | 0.644 | 0.0028 |
| 95f8ff42ab06a32b618f4113f6c759d4 | *X. muta*: Sept. 2017 | Proteobacteria_ Gammaproteobacteria_NA_NA | 0.559 | 0.0045 |
| 6da97eb3f6956e0620d1d085d3785d8c | *X. muta*: Sept. 2017 | Proteobacteria_ Gammaproteobacteria_NA_NA | 0.521 | 0.0422 |
| 7170f084336b39b16cd789d8b5441d64 | *X. muta*: Sept. 2017 | Proteobacteria_ Gammaproteobacteria_NA_NA | 0.504 | 0.0485 |
| 5b2924c62be2729f16eb5cdc55963981 | *X. muta*: Sept. 2017 | Schekmanbacteria_ Schekmanbacteria_ Schekmanbacteria_ Schekmanbacteria | 0.432 | 0.0397 |

| | | | | |
|---|---|---|---|---|
| aa6ee3cb1efe3069e1e2f36f804bfc4d | *X. muta*: Sept. 2017 | Spirochaetota_ Leptospirae_ Leptospiraceae_ uncultured | 0.48 | 0.042 |
| 5bc774c658da095cda68b735f062ee38 | *X. muta*: Sept. 2017 | Verrucomicrobiota_ Lentisphaeria_ Lentisphaeraceae_ Lentisphaera | 0.448 | 0.031 |
| efc9895ab1b9737085a8096ee160159f | *X. muta*: Sept. 2017 | Verrucomicrobiota_ Verrucomicrobiae_ Akkermansiaceae_ Akkermansia | 0.424 | 0.0479 |
| 6c13ba5f0d87c53e32dd3a4ef07bf962 | *X. muta*: Sept. 2017 | Verrucomicrobiota_ Verrucomicrobiae_ Puniceicoccaceae_ Cerasicoccus | 0.464 | 0.0154 |
| 7f32f1ecd63c5ddcd155763f9ee6b0d0 | *X. muta*: Oct. 2017 | Acidobacteriota_ Thermoanaerobaculia_ Thermoanaerobaculaceae_ Subgroup 10 | 0.546 | 0.0112 |
| d1ce2103d1dd82c1bde3b9517bec9c26 | *X. muta*: Oct. 2017 | Acidobacteriota_ Vicinamibacteria_ uncultured_ uncultured | 0.749 | 0.0002 |
| d7c7d42275a3b107017867c1841ba73e | *X. muta*: Oct. 2017 | Actinobacteriota_ Acidimicrobiia_ Microtrichaceae_ Sva0996 marine group | 0.67 | 0.0018 |
| d6c0e1d529b5fe9c68db5bd52ea56a4a | *X. muta*: Oct. 2017 | Actinobacteriota_ Acidimicrobiia_ Microtrichaceae_ Sva0996 marine group | 0.569 | 0.0263 |
| 927ea3701a1f926737fb51fe2b6d1508 | *X. muta*: Oct. 2017 | Actinobacteriota_ Acidimicrobiia_ Microtrichaceae_ Sva0996 marine group | 0.537 | 0.008 |
| 6c47cfd718e103fa990a752a2051ba8f | *X. muta*: Oct. 2017 | Chloroflexi_ Anaerolineae_ A4b_ A4b | 0.741 | 0.0001 |
| 7c08cad33533a031b243d96562084e57 | *X. muta*: Oct. 2017 | Chloroflexi_ Anaerolineae_ A4b_ A4b | 0.66 | 0.0025 |
| 40bfbfddd8735d95fc9ea8f6dc1892c7 | *X. muta*: Oct. 2017 | Chloroflexi_ Anaerolineae_ A4b_ A4b | 0.563 | 0.0236 |
| 8b828f9ec22b228578a34b2cefb99f69 | *X. muta*: Oct. 2017 | Chloroflexi_ Anaerolineae_ A4b_ A4b | 0.555 | 0.0018 |
| 2487ce3e9deb4bce09aa17554dc03844 | *X. muta*: Oct. 2017 | Chloroflexi_ Anaerolineae_ A4b_ A4b | 0.53 | 0.0171 |
| 9712f0938ecd7fa06530dadf52fe8ee2 | *X. muta*: Oct. 2017 | Chloroflexi_ Anaerolineae_ A4b_ A4b | 0.509 | 0.0265 |
| d68e0573864219ae1dda6c41e5f8f867 | *X. muta*: Oct. 2017 | Chloroflexi_ Anaerolineae_ A4b_ A4b | 0.466 | 0.024 |
| 01049762a54a77c2bb32bb354684a993 | *X. muta*: Oct. 2017 | Chloroflexi_ Anaerolineae_ Caldilineaceae_ uncultured | 0.657 | 0.0025 |
| ccf3f8f80dddec76558390bd84b357a8 | *X. muta*: Oct. 2017 | Chloroflexi_ Anaerolineae_ Caldilineaceae_ uncultured | 0.552 | 0.015 |
| c7373a2fd11b273094dcf3c80b93cf9e | *X. muta*: Oct. 2017 | Chloroflexi_ Dehalococcoidia_ SAR202_clade_ SAR202_clade | 0.759 | 0.0005 |
| 7ae614eb9fbeb2f45a04d7c7ad4de204 | *X. muta*: Oct. 2017 | Chloroflexi_ Dehalococcoidia_ SAR202_clade_ SAR202_clade | 0.606 | 0.0049 |
| 75db51d9f14ca50de3ba1d068b1d0330 | *X. muta*: Oct. 2017 | Chloroflexi_ Dehalococcoidia_ SAR202_clade_ SAR202_clade | 0.586 | 0.0042 |
| 550752ae02ff565b040f8a21ad71f1f2 | *X. muta*: Oct. 2017 | Chloroflexi_ Dehalococcoidia_ SAR202_clade_ SAR202_clade | 0.545 | 0.0257 |
| 4693cb685c09730128d5dfc9a33c98cc | *X. muta*: Oct. 2017 | Chloroflexi_ Dehalococcoidia_ SAR202_clade_ SAR202_clade | 0.536 | 0.0269 |
| f0944ca98f8f123c25908125f2ba4f19 | *X. muta*: Oct. 2017 | Chloroflexi_ Dehalococcoidia_ SAR202_clade_ SAR202_clade | 0.532 | 0.0206 |
| 08b6e4c68faa7baaec83bf05aa8cfbab | *X. muta*: Oct. 2017 | Chloroflexi_ Dehalococcoidia_ SAR202_clade_ SAR202_clade | 0.532 | 0.015 |
| d5399753c60df34478ee1900db66ee08 | *X. muta*: Oct. 2017 | Chloroflexi_ Dehalococcoidia_ SAR202_clade_ SAR202_clade | 0.504 | 0.0391 |
| aaecb7e4aeba785205fc66fc08a4b6c6 | *X. muta*: Oct. 2017 | Chloroflexi_ Dehalococcoidia_ SAR202_clade_ SAR202_clade | 0.491 | 0.0311 |
| b08d9b9d8dbd928e15573d0a2e0ae212 | *X. muta*: Oct. 2017 | Chloroflexi_ Dehalococcoidia_ SAR202_clade_ SAR202_clade | 0.489 | 0.0353 |
| 32f17c74bc413deedf842c4f7b700692 | *X. muta*: Oct. 2017 | Chloroflexi_ Dehalococcoidia_ SAR202_clade_ SAR202_clade | 0.459 | 0.045 |
| 9593f16e0987c113ec18e7cd27dc560f | *X. muta*: Oct. 2017 | Chloroflexi_ JG30-KF-CM66_ JG30-KF-CM66_ JG30-KF-CM66 | 0.609 | 0.0054 |
| b108ae293c699912b2c7751e7dc2bbc8 | *X. muta*: Oct. 2017 | Chloroflexi_ JG30-KF-CM66_ JG30-KF-CM66_ JG30-KF-CM66 | 0.534 | 0.0171 |
| ed9c06eb0a8196aebb2f09a82fc0533d | *X. muta*: Oct. 2017 | Chloroflexi_ TK10_ TK10_ TK10 | 0.717 | 0.0001 |
| 8da1b458ed9f71100182739bb9629ab5 | *X. muta*: Oct. 2017 | Chloroflexi_ TK10_ TK10_ TK10 | 0.711 | 0.0009 |
| 5b7f33df401ec536fd38379dd484ee1a | *X. muta*: Oct. 2017 | Chloroflexi_ TK10_ TK10_ TK10 | 0.661 | 0.0012 |
| 6362fb3cb74cab96642ff4f33b51681c | *X. muta*: Oct. 2017 | Chloroflexi_ TK17_ TK17_ TK17 | 0.627 | 0.0018 |
| ea206c94e913c3092fbb68ce26a3a934 | *X. muta*: Oct. 2017 | Chloroflexi_ TK17_ TK17_ TK17 | 0.544 | 0.028 |
| cdfcdee49a590bdafa5cc71e2b9f27e8 | *X. muta*: Oct. 2017 | Chloroflexi_ TK17_ TK17_ TK17 | 0.527 | 0.0213 |
| 68f7d94efff7be569754c938841b681b | *X. muta*: Oct. 2017 | Chloroflexi_ TK17_ TK17_ TK17 | 0.515 | 0.0236 |
| cc5c9e131fd620c3ea9db0fc1ffd96b5 | *X. muta*: Oct. 2017 | Chloroflexi_ TK30_ TK30_ TK30 | 0.49 | 0.0434 |
| 2c740e298871de196cb2f1b9a62ef681 | *X. muta*: Oct. 2017 | Firmicutes_ Clostridia_ Lachnospiraceae_ Lachnoclostridium | 0.545 | 0.0269 |

| | | | | |
|---|---|---|---|---|
| 4bc44f8decfa3119cb937d4d7c5676c3 | *X. muta*: Oct. 2017 | Nanoarchaeota_ Nanoarchaeia_ Woesearchaeales_ Woesearchaeales | 0.615 | 0.0021 |
| 9880cdd53cf868c39180bb277460bb00 | *X. muta*: Oct. 2017 | Nanoarchaeota_ Nanoarchaeia_ Woesearchaeales_ Woesearchaeales | 0.491 | 0.0238 |
| fdf048306903181c90b7e09d24c7561f | *X. muta*: Oct. 2017 | Poribacteria_ Poribacteria_ Poribacteria_ Poribacteria | 0.649 | 0.0025 |
| 2a7e09460e5cbe9f0d670b20c61262be | *X. muta*: Oct. 2017 | Proteobacteria_ Alphaproteobacteria_NA_NA | 0.56 | 0.0067 |
| 7aa11d38ed74061d0b07432f2574d29b | *A. clathrodes*: Aug. 2016 | Acidobacteriota_ Vicinamibacteria_ Vicinamibacteraceae_ Vicinamibacteraceae | 0.507 | 0.0002 |
| 8b180e80765379a5ecf1276e10f85163 | *A. clathrodes*: Aug. 2016 | Actinobacteriota_ Acidimicrobiia_ Microtrichaceae_ Sva0996 marine group | 0.312 | 0.0165 |
| fe2715780b37f5b29948878e4eac8022 | *A. clathrodes*: Aug. 2016 | Actinobacteriota_ Acidimicrobiia_ uncultured_ uncultured | 0.736 | 0.0001 |
| 46096113c55d29fb63a97c7ccb30da3d | *A. clathrodes*: Aug. 2016 | Actinobacteriota_ Acidimicrobiia_ uncultured_ uncultured | 0.721 | 0.0001 |
| a8e47b14fa1ec278af41bc7d78e8fb48 | *A. clathrodes*: Aug. 2016 | Actinobacteriota_ Acidimicrobiia_ uncultured_ uncultured | 0.697 | 0.0001 |
| c6322bf243c8c6fcbac46304d1e2e6af | *A. clathrodes*: Aug. 2016 | Bacteroidota_ Bacteroidia_ Amoebophilaceae_ uncultured | 0.376 | 0.0032 |
| 30fa3d09c35c0728473cef60534efa81 | *A. clathrodes*: Aug. 2016 | Bacteroidota_ Bacteroidia_ Flavobacteriaceae_ uncultured | 0.376 | 0.0134 |
| ccd8d5e31ad9bfa58066251ea1138225 | *A. clathrodes*: Aug. 2016 | Bacteroidota_ Bacteroidia_ Flavobacteriaceae_NA | 0.343 | 0.0023 |
| b4d6690e31369213642734660361a5ca | *A. clathrodes*: Aug. 2016 | Bacteroidota_ Bacteroidia_ Lentimicrobiaceae_ Lentimicrobium | 0.429 | 0.0027 |
| d2ad6d6a1bc1cb69a430066ca841b332 | *A. clathrodes*: Aug. 2016 | Bacteroidota_ Bacteroidia_ Lentimicrobiaceae_ Lentimicrobium | 0.355 | 0.0025 |
| 64805be33440e427c18e31e0d5e6094b | *A. clathrodes*: Aug. 2016 | Bacteroidota_ Bacteroidia_ Saprospiraceae_ Phaeodactylibacter | 0.379 | 0.004 |
| f3a2c825531c54cfc115b7b818561953 | *A. clathrodes*: Aug. 2016 | Bacteroidota_ Bacteroidia_ Saprospiraceae_ uncultured | 0.402 | 0.0034 |
| 408492a1941643c50c8a9c5e3614b48c | *A. clathrodes*: Aug. 2016 | Bacteroidota_ Bacteroidia_NA_NA | 0.541 | 0.0001 |
| 04243c3adce2a4c14c7b9d8dbd652f0c | *A. clathrodes*: Aug. 2016 | Bacteroidota_ Bacteroidia_NA_NA | 0.427 | 0.0001 |
| 90d5ff29c4c5c55af81629ecee606369 | *A. clathrodes*: Aug. 2016 | Bdellovibrionota_ Bdellovibrionia_ Bdellovibrionaceae_ Bdellovibrio | 0.521 | 0.0003 |
| 3000dd01eeaaf3321f23432476fee64c | *A. clathrodes*: Aug. 2016 | Bdellovibrionota_ Bdellovibrionia_ Bdellovibrionaceae_ Bdellovibrio | 0.431 | 0.0049 |
| c81067170b6c2eb0e8d61e2e749ec900 | *A. clathrodes*: Aug. 2016 | Bdellovibrionota_ Bdellovibrionia_ Bdellovibrionaceae_ Bdellovibrio | 0.421 | 0.0071 |
| 233e1e5b95ea054309238177bde19a48 | *A. clathrodes*: Aug. 2016 | Bdellovibrionota_ Bdellovibrionia_ Bdellovibrionaceae_ Bdellovibrio | 0.368 | 0.0138 |
| 8412293eee50ae30da499075bf79bcc8 | *A. clathrodes*: Aug. 2016 | Bdellovibrionota_ Bdellovibrionia_ Bdellovibrionaceae_ Bdellovibrio | 0.352 | 0.0024 |
| 9ad70c69d09459eeb13b025b516f4b03 | *A. clathrodes*: Aug. 2016 | Bdellovibrionota_ Bdellovibrionia_ Bdellovibrionaceae_ Bdellovibrio | 0.332 | 0.0351 |
| 1fffdac4862eb560f3094dfc329f0aa6 | *A. clathrodes*: Aug. 2016 | Bdellovibrionota_ Bdellovibrionia_ Bdellovibrionaceae_ Bdellovibrio | 0.323 | 0.0026 |
| 18eefd9b84d6e3415f914c8f3142db0d | *A. clathrodes*: Aug. 2016 | Bdellovibrionota_ Oligoflexia_ 0319-6G20_ 0319-6G20 | 0.277 | 0.0025 |
| b7740c2a02d4af9589023217ef8a7342 | *A. clathrodes*: Aug. 2016 | Campilobacterota_ Campylobacteria_ Arcobacteraceae_NA | 0.264 | 0.0154 |
| eb16874a5a59c61984a11437e522e063 | *A. clathrodes*: Aug. 2016 | Campilobacterota_ Campylobacteria_ Campylobacteraceae_ Campylobacter | 0.352 | 0.0141 |
| 9d5f7010907617ff13fa874b18b06b5d | *A. clathrodes*: Aug. 2016 | Chloroflexi_ Dehalococcoidia_ SAR202_clade_ SAR202_clade | 0.311 | 0.0069 |
| b3704fd959e1a374efa75d979a13e509 | *A. clathrodes*: Aug. 2016 | Crenarchaeota_ Nitrososphaeria_ Nitrosopumilaceae_ Candidatus_Nitrosopelagicus | 0.357 | 0.0177 |
| 6a1e4b41f741b4fd1a7adac461779ab7 | *A. clathrodes*: Aug. 2016 | Cyanobacteria_ Cyanobacteriia_ Cyanobiaceae_ Prochlorococcus | 0.477 | 0.0015 |
| 53f00a14f02bb0de5a41df9c35b92b65 | *A. clathrodes*: Aug. 2016 | Cyanobacteria_ Vampirivibrionia_ Gastranaerophilales_ Gastranaerophilales | 0.36 | 0.0001 |
| 822c0e34cd1c98c61479cbc852191360 | *A. clathrodes*: Aug. 2016 | Desulfobacterota_ Desulfobacteria_ Desulfolunaceae_ Desulfoluna | 0.489 | 0.0001 |
| 19d1f310473cc39014864b2178a400c8 | *A. clathrodes*: Aug. 2016 | Desulfobacterota_ Desulfovibrionia_ Desulfovibrionaceae_ Desulfovibrio | 0.451 | 0.0007 |
| ed0789829790bd0ab4a46e3f53fe0a59 | *A. clathrodes*: Aug. 2016 | Desulfobacterota_ Desulfovibrionia_ Desulfovibrionaceae_ Halodesulfovibrio | 0.469 | 0.0003 |
| 08b458863412f315b69aff049a88435b | *A. clathrodes*: Aug. 2016 | Desulfobacterota_ Desulfovibrionia_ Desulfovibrionaceae_ Halodesulfovibrio | 0.439 | 0.0001 |
| 6088e2f047a9fd144b4310ad15148a2e | *A. clathrodes*: Aug. 2016 | Desulfobacterota_ Desulfovibrionia_ Desulfovibrionaceae_ Halodesulfovibrio | 0.393 | 0.0024 |
| e4a5985be454d1e8a3d50a2ab6ea7a8b | *A. clathrodes*: Aug. 2016 | Desulfobacterota_ Desulfovibrionia_ Desulfovibrionaceae_ Halodesulfovibrio | 0.386 | 0.0001 |
| e09f2d58d292332880fe9e37607469e1 | *A. clathrodes*: Aug. 2016 | Desulfobacterota_ Desulfovibrionia_ Desulfovibrionaceae_ Halodesulfovibrio | 0.385 | 0.0001 |
| c34517a788b2dc8691c666621db1ac15 | *A. clathrodes*: Aug. 2016 | Desulfobacterota_ Desulfovibrionia_ Desulfovibrionaceae_ Halodesulfovibrio | 0.317 | 0.0001 |

| | | | | |
|---|---|---|---|---|
| 0261a50c7edb68abf5e65bb5094ce011 | *A. clathrodes*: Aug. 2016 | Desulfobacterota_ Desulfovibrionia_ Desulfovibrionaceae_NA | 0.524 | 0.0001 |
| e51947121c608cf15031596f9d0c09eb | *A. clathrodes*: Aug. 2016 | Desulfobacterota_ Desulfovibrionia_ Desulfovibrionaceae_NA | 0.397 | 0.0001 |
| 5948f84e364747615718552f4bfded36 | *A. clathrodes*: Aug. 2016 | Desulfobacterota_ Desulfovibrionia_ Desulfovibrionaceae_NA | 0.385 | 0.0004 |
| ca80e91cdaef1032d981b2ce7313ec15 | *A. clathrodes*: Aug. 2016 | Firmicutes_ Clostridia_ Clostridiaceae_ uncultured | 0.352 | 0.0029 |
| 216ac98287765ffa4ac3e18f96930f49 | *A. clathrodes*: Aug. 2016 | Firmicutes_ Clostridia_ Clostridiaceae_ uncultured | 0.34 | 0.0155 |
| 7b924ed3d8413fcfa05dff0632ac6c1d | *A. clathrodes*: Aug. 2016 | Firmicutes_ Clostridia_ Clostridiaceae_ uncultured | 0.325 | 0.0004 |
| eb46361ce52904349aabf94a3ad30512 | *A. clathrodes*: Aug. 2016 | Firmicutes_ Clostridia_ Clostridiaceae_ uncultured | 0.306 | 0.0002 |
| 91b3f60afa0840334f83e6ae6d5604ef | *A. clathrodes*: Aug. 2016 | Firmicutes_ Clostridia_ Lachnospiraceae_ Epulopiscium | 0.336 | 0.0145 |
| 978ee4c0178d38509290b408643ec5bb | *A. clathrodes*: Aug. 2016 | Firmicutes_ Clostridia_ Peptostreptococcaceae_ Romboutsia | 0.412 | 0.0074 |
| 23bd1417575bff64af0e2fdae2bd472e | *A. clathrodes*: Aug. 2016 | Firmicutes_ Clostridia_ Peptostreptococcales-Tissierellales_ uncultured | 0.421 | 0.0031 |
| 4edd3f6a29d963455be4d87f34391e1a | *A. clathrodes*: Aug. 2016 | Gemmatimonadota_ BD2-11_terrestrial_group_ NA_NA | 0.444 | 0.0041 |
| 852ac7b098065c718eedec0a99826ab5 | *A. clathrodes*: Aug. 2016 | Myxococcota_ bacteriap25_ bacteriap25_ bacteriap25 | 0.3 | 0.0137 |
| 6d0aeefe127607d7462cef7588d2fd91 | *A. clathrodes*: Aug. 2016 | Planctomycetota_ Phycisphaerae_ AKAU3564 sediment group_NA | 0.446 | 0.0033 |
| f7c9e95888f381208f289c617f6a7a2b | *A. clathrodes*: Aug. 2016 | Planctomycetota_ Phycisphaerae_ AKAU3564_sediment_group_NA | 0.376 | 0.0137 |
| d0495034caec14b095ecbf874e9d17a1 | *A. clathrodes*: Aug. 2016 | Planctomycetota_ Phycisphaerae_ Phycisphaeraceae_NA | 0.515 | 0.0001 |
| 41be6c83e42c6ce36f33303352e31e91 | *A. clathrodes*: Aug. 2016 | Poribacteria_ Poribacteria_ Poribacteria_ Poribacteria | 0.575 | 0.0001 |
| fa5832d66e27b928e9306252ce53005a | *A. clathrodes*: Aug. 2016 | Porifera_ Demospongiae_ Demospongiae_ Demospongiae | 0.715 | 0.0001 |
| b14b1e29552a7a4a6b7e4af37d28a31e | *A. clathrodes*: Aug. 2016 | Proteobacteria_ Alphaproteobacteria_ AEGEAN-169 marine group | 0.401 | 0.0095 |
| 08700e14c442225a1f498e75977809d0 | *A. clathrodes*: Aug. 2016 | Proteobacteria_ Alphaproteobacteria_ AEGEAN-169 marine group | 0.371 | 0.0134 |
| 85a534028047f4407f95e6422a8bdb8c | *A. clathrodes*: Aug. 2016 | Proteobacteria_ Alphaproteobacteria_ AT-s3-44_ AT-s3-44 | 0.346 | 0.0136 |
| b23273610f0956889dd291a10db0b1eb | *A. clathrodes*: Aug. 2016 | Proteobacteria_ Alphaproteobacteria_ Beijerinckiaceae_ Methylobacterium | 0.436 | 0.0003 |
| 027e868b683e709d600ee448681cebcb | *A. clathrodes*: Aug. 2016 | Proteobacteria_ Alphaproteobacteria_ Beijerinckiaceae_ Methylobacterium | 0.392 | 0.0133 |
| 1947fd8de1d6f07a76f44a64e8facee6 | *A. clathrodes*: Aug. 2016 | Proteobacteria_ Alphaproteobacteria_ Beijerinckiaceae_ Methylobacterium | 0.376 | 0.0156 |
| 5546361caf3e42e50fb56d85a11571f2 | *A. clathrodes*: Aug. 2016 | Proteobacteria_ Alphaproteobacteria_ Clade_I_ Clade_Ia | 0.511 | 0.001 |
| 9ef7f4be295f211b054ddfcd091f22bd | *A. clathrodes*: Aug. 2016 | Proteobacteria_ Alphaproteobacteria_ Clade_I_ Clade_Ia | 0.386 | 0.0148 |
| 05c843221801cb3b08e96de54d2e6ac0 | *A. clathrodes*: Aug. 2016 | Proteobacteria_ Alphaproteobacteria_ Clade_I_ Clade_Ib | 0.482 | 0.0009 |
| ca1483f56851807381b6bf55e37cb7c4 | *A. clathrodes*: Aug. 2016 | Proteobacteria_ Alphaproteobacteria_ Clade_I_ Clade_Ib | 0.463 | 0.0034 |
| 2aa8c06efa876d080ee031bfe42420c9 | *A. clathrodes*: Aug. 2016 | Proteobacteria_ Alphaproteobacteria_ Clade_II_ Clade_II | 0.642 | 0.0001 |
| 6de40fcdea2aa22c95bc4f5b856c9b03 | *A. clathrodes*: Aug. 2016 | Proteobacteria_ Alphaproteobacteria_ Clade_III_ Clade_III | 0.423 | 0.003 |
| 3fa8df16cb367c8e2d5e0ab3f2c4871e | *A. clathrodes*: Aug. 2016 | Proteobacteria_ Alphaproteobacteria_ Rhodobacteraceae_ Albidovulum | 0.393 | 0.0051 |
| 606952d29ba7c7ec952786841bc53515 | *A. clathrodes*: Aug. 2016 | Proteobacteria_ Alphaproteobacteria_ Rhodobacteraceae_NA | 0.441 | 0.0033 |
| bbfed89cd414e1e7c217c58494857f73 | *A. clathrodes*: Aug. 2016 | Proteobacteria_ Alphaproteobacteria_ Rhodobacteraceae_NA | 0.435 | 0.0031 |
| 3d870f602b139ffda8f915b9705040a2 | *A. clathrodes*: Aug. 2016 | Proteobacteria_ Alphaproteobacteria_ Rhodobacteraceae_NA | 0.387 | 0.0164 |
| b6d06f5862175ad99eb22b4c1e919b72 | *A. clathrodes*: Aug. 2016 | Proteobacteria_ Alphaproteobacteria_ Rhodobacteraceae_NA | 0.338 | 0.0136 |
| be5c0927fe342c83c48ecc80193b422e | *A. clathrodes*: Aug. 2016 | Proteobacteria_ Alphaproteobacteria_ Sphingomonadaceae_ Sphingobium | 0.406 | 0.0028 |
| 8cb4eec725fe8e690c06103916bba72e | *A. clathrodes*: Aug. 2016 | Proteobacteria_ Alphaproteobacteria_ Sphingomonadaceae_ Sphingomonas | 0.382 | 0.0033 |
| cd760f81691f5a5d0ea7714f2b347f93 | *A. clathrodes*: Aug. 2016 | Proteobacteria_ Alphaproteobacteria_ Stappiaceae_ Pseudovibrio | 0.443 | 0.0003 |
| 95ba0218fab70b8c4807eaac1c451619 | *A. clathrodes*: Aug. 2016 | Proteobacteria_ Alphaproteobacteria_ Stappiaceae_ Pseudovibrio | 0.387 | 0.0035 |
| 88b2cb86c0c9dee735e12d55a0880daf | *A. clathrodes*: Aug. 2016 | Proteobacteria_ Alphaproteobacteria_ Stappiaceae_ Pseudovibrio | 0.34 | 0.0032 |
| 2a9193fd0156adc0f559818b6fed16bb | *A. clathrodes*: Aug. 2016 | Proteobacteria_ Alphaproteobacteria_ uncultured_ uncultured | 0.376 | 0.0037 |

| | | | | |
|---|---|---|---|---|
| 0f1768344ceaef637f6bc94b7eb20259 | *A. clathrodes*: Aug. 2016 | Proteobacteria_ Alphaproteobacteria_NA_NA | 0.327 | 0.0005 |
| de00632a6754273bdef12fed7bfd4eb6 | *A. clathrodes*: Aug. 2016 | Proteobacteria_ Gammaproteobacteria_ Coxiellaceae_ Coxiella | 0.707 | 0.0001 |
| aa404544885ff0999f721532694fdcc3 | *A. clathrodes*: Aug. 2016 | Proteobacteria_ Gammaproteobacteria_ Coxiellaceae_ Coxiella | 0.582 | 0.0001 |
| 3169c77e9620902b938490dc39b454d1 | *A. clathrodes*: Aug. 2016 | Proteobacteria_ Gammaproteobacteria_ Coxiellaceae_ Coxiella | 0.393 | 0.004 |
| 8c4c76bea3b0db83df3593f73509f166 | *A. clathrodes*: Aug. 2016 | Proteobacteria_ Gammaproteobacteria_ Ectothiorhodospiraceae_ uncultured | 0.346 | 0.0316 |
| 35d3daf29fa0b8028e1845576666c3e5 | *A. clathrodes*: Aug. 2016 | Proteobacteria_ Gammaproteobacteria_ Endozoicomonadaceae_ Endozoicomonas | 0.416 | 0.0001 |
| 6d472a1d79dae690d28607258767704e | *A. clathrodes*: Aug. 2016 | Proteobacteria_ Gammaproteobacteria_ Endozoicomonadaceae_ Endozoicomonas | 0.381 | 0.0001 |
| f24139a25fa20687f6933c1988c06d23 | *A. clathrodes*: Aug. 2016 | Proteobacteria_ Gammaproteobacteria_ Endozoicomonadaceae_ Endozoicomonas | 0.323 | 0.0002 |
| f2335497138a785fb4288dfcf64a56c0 | *A. clathrodes*: Aug. 2016 | Proteobacteria_ Gammaproteobacteria_ Halieaceae_ OM60(NOR5)_clade | 0.352 | 0.0151 |
| 2ef8937fcfef105310d9aae9f65c08ab | *A. clathrodes*: Aug. 2016 | Proteobacteria_ Gammaproteobacteria_ Kangiellaceae_ Aliikangiella | 0.341 | 0.0036 |
| 03099841ef74f32270c525d9d5492dac | *A. clathrodes*: Aug. 2016 | Proteobacteria_ Gammaproteobacteria_ KI89A_clade_ KI89A_clade | 0.396 | 0.0088 |
| 50a48811db3c95400741562125e619df | *A. clathrodes*: Aug. 2016 | Proteobacteria_ Gammaproteobacteria_ KI89A_clade_ KI89A_clade | 0.372 | 0.0162 |
| 13c8bd0c6fb64d5b49440dffa4ed29f4 | *A. clathrodes*: Aug. 2016 | Proteobacteria_ Gammaproteobacteria_ MBMPE27_ MBMPE27 | 0.539 | 0.0001 |
| ce4587b6e4b0dd00976083f9756c8119 | *A. clathrodes*: Aug. 2016 | Proteobacteria_ Gammaproteobacteria_ Nitrosococcaceae_ AqS1 | 0.575 | 0.0001 |
| a099cc5242221500cb80a03ec6bc6305 | *A. clathrodes*: Aug. 2016 | Proteobacteria_ Gammaproteobacteria_ Nitrosococcaceae_ AqS1 | 0.451 | 0.0031 |
| 4bf34e939f3102402abec9deab0319b3 | *A. clathrodes*: Aug. 2016 | Proteobacteria_ Gammaproteobacteria_ Nitrosococcaceae_ AqS1 | 0.382 | 0.0158 |
| f7f304f78d06f1666249749605294643 | *A. clathrodes*: Aug. 2016 | Proteobacteria_ Gammaproteobacteria_ Piscirickettsiaceae_ Endoecteinascidia | 0.478 | 0.0038 |
| 9442980515ce7f911bb4cc9c372db4e8 | *A. clathrodes*: Aug. 2016 | Proteobacteria_ Gammaproteobacteria_ SAR86_clade_ SAR86_clade | 0.62 | 0.0001 |
| 9ec997c700d09a502a482a43b707b85c | *A. clathrodes*: Aug. 2016 | Proteobacteria_ Gammaproteobacteria_ SAR86_clade_ SAR86_clade | 0.567 | 0.0001 |
| 7715944abdcc3856acebb58ede6b09c8 | *A. clathrodes*: Aug. 2016 | Proteobacteria_ Gammaproteobacteria_ SAR86_clade_ SAR86_clade | 0.379 | 0.0202 |
| c254b9aefd8b2a221071777fd30660ff | *A. clathrodes*: Aug. 2016 | Proteobacteria_ Gammaproteobacteria_ Spongiibacteraceae_ BD1-7_clade | 0.304 | 0.0065 |
| 59119808c8e414460016157eca84739b | *A. clathrodes*: Aug. 2016 | Proteobacteria_ Gammaproteobacteria_ Thioglobaceae_ SUP05_cluster | 0.476 | 0.0007 |
| b47b6e5813cfd4b7690b522531f10aa4 | *A. clathrodes*: Aug. 2016 | Proteobacteria_ Gammaproteobacteria_ Thioglobaceae_ SUP05_cluster | 0.393 | 0.0146 |
| e99730501d8a2d0639cedc50419fecb7 | *A. clathrodes*: Aug. 2016 | Proteobacteria_ Gammaproteobacteria_ Vibrionaceae_ uncultured | 0.336 | 0.0371 |
| 4ade1c6fac0b565acb1f78742cf01456 | *A. clathrodes*: Aug. 2016 | Proteobacteria_ Gammaproteobacteria_ Vibrionaceae_ Vibrio | 0.529 | 0.0003 |
| d7d7b0e69416842f55a6c6ebd78cd03e | *A. clathrodes*: Aug. 2016 | Proteobacteria_ Gammaproteobacteria_ Vibrionaceae_ Vibrio | 0.408 | 0.014 |
| 28c5ce89edc28a7b4eed336644b30acc | *A. clathrodes*: Aug. 2016 | Proteobacteria_ Gammaproteobacteria_NA_NA | 0.373 | 0.0204 |
| efa28eb526ec0c3caa54fd35809774e1 | *A. clathrodes*: Aug. 2016 | SAR324 clade_Marine group B_ NA_NA | 0.393 | 0.0146 |
| e1f8a5ea0cc1011994812c99142f2b01 | *A. clathrodes*: Aug. 2016 | Spirochaetota_ Leptospirae_ Leptospiraceae_ Leptospiraceae | 0.345 | 0.02 |
| baee8e7e3efe158cb4fb37da60fab9d0 | *A. clathrodes*: Aug. 2016 | Spirochaetota_ Leptospirae_ Leptospiraceae_ Leptospiraceae | 0.341 | 0.0156 |
| 7c6a7aa3a98f8b0a39920e8de0db2521 | *A. clathrodes*: Aug. 2016 | Spirochaetota_ Spirochaetia_ Spirochaetaceae_ uncultured | 0.376 | 0.0142 |
| 7b803559a6a82a6ee058b239731b3b01 | *A. clathrodes*: Aug. 2016 | Verrucomicrobiota_ Lentisphaeria_ P.palmC41_ P.palmC41 | 0.365 | 0.0003 |
| 874359ce05108ae6d0eb15976b76f5cb | *A. clathrodes*: Aug. 2016 | Marinimicrobia_SAR406 clade_NA_NA | 0.448 | 0.0036 |
| 1c1bb254b8b270264a7a3bd48dde5116 | *A. clathrodes*: Oct. 2017 | Acidobacteriota_ Vicinamibacteria_ uncultured_ uncultured | 0.616 | 0.0001 |
| c5e220d55b91be9dcaa6217b64ad36fd | *A. clathrodes*: Oct. 2017 | Acidobacteriota_ Vicinamibacteria_ uncultured_ uncultured | 0.36 | 0.0217 |
| a0ded0719445c2312f5eab1b3a385fc7 | *A. clathrodes*: Oct. 2017 | Acidobacteriota_ Vicinamibacteria_ uncultured_ uncultured | 0.359 | 0.0202 |
| 69074d2252604b321238eff89c873a2b | *A. clathrodes*: Oct. 2017 | Acidobacteriota_ Vicinamibacteria_ Vicinamibacteraceae_ Vicinamibacteraceae | 0.6 | 0.0001 |
| e128c73af1ad25cd4d2e978532a76a9b | *A. clathrodes*: Oct. 2017 | Actinobacteriota_ Acidimicrobiia_ Microtrichaceae_ Sva0996 marine group | 0.418 | 0.0067 |
| 9515f5f55d2d2d059d774c50a6a72e3f | *A. clathrodes*: Oct. 2017 | Chloroflexi_ JG30-KF-CM66_ JG30-KF-CM66_ JG30-KF-CM66 | 0.482 | 0.0009 |
| 73ba47d29bd12c5103945a1607379fab | *A. clathrodes*: Oct. 2017 | Cyanobacteria_ Cyanobacteriia_ Cyanobiaceae_ Cyanobium | 0.344 | 0.0182 |

| | | | | |
|---|---|---|---|---|
| 5c4f0df5a646a74943e9f0818ff50cf6 | *A. clathrodes*: Oct. 2017 | Cyanobacteria_ Cyanobacteriia_ Cyanobiaceae_ Synechococcus | 0.419 | 0.005 |
| aa02f8dcbedb2d711bc22c8affde3b7c | *A. clathrodes*: Oct. 2017 | Cyanobacteria_ Cyanobacteriia_ Cyanobiaceae_ Synechococcus | 0.396 | 0.0028 |
| cd0283e6687d70df913b1bc1fd730133 | *A. clathrodes*: Oct. 2017 | Planctomycetota_ Planctomycetes_ Pirellulaceae_ Blastopirellula | 0.385 | 0.0012 |
| c12c6a15f208df5992ed2c7d14fd2a12 | *A. clathrodes*: Oct. 2017 | Planctomycetota_ Planctomycetes_ Pirellulaceae_ Rubripirellula | 0.397 | 0.0022 |
| b6a8bbc59e71239ab4a9e0303ba14d5a | *A. clathrodes*: Oct. 2017 | Planctomycetota_ Planctomycetes_ Pirellulaceae_ uncultured | 0.327 | 0.0254 |
| 30ed29e8b2a60a549e95d41a89e8a427 | *A. clathrodes*: Oct. 2017 | Planctomycetota_ Planctomycetes_ Rubinisphaeraceae_ uncultured | 0.339 | 0.0496 |
| 3738b275a37a3ea7197ca89533b3abba | *A. clathrodes*: Oct. 2017 | Proteobacteria_ Gammaproteobacteria_ Legionellaceae_ Legionella | 0.326 | 0.0496 |
| ce0640e9b5d1f535dcc8260e2338f4aa | Seawater: Aug. 2016 | Actinobacteriota_ Acidimicrobiia_ Actinomarinaceae_Actinomarina | 0.571 | 0.0043 |
| ea9c34dae90ecbf15f06deefb6e54dbb | Seawater: Aug. 2016 | Actinobacteriota_ Acidimicrobiia_ Actinomarinaceae_Actinomarina | 0.545 | 0.0069 |
| df15b4b2180c1412c93a0c471b5d9fca | Seawater: Aug. 2016 | Bacteroidota_ Bacteroidia_ Cryomorphaceae_ uncultured | 0.626 | 0.0003 |
| dd66766b2fe3a746a47e7d155ac28bef | Seawater: Aug. 2016 | Bacteroidota_ Bacteroidia_ Cryomorphaceae_ uncultured | 0.577 | 0.0067 |
| 711b27e1edc410ffac57c081e97c5a7c | Seawater: Aug. 2016 | Bacteroidota_ Bacteroidia_ Cyclobacteriaceae_ Fabibacter | 0.53 | 0.0149 |
| 25eda3a8deea4880061ba8b630c5e723 | Seawater: Aug. 2016 | Bacteroidota_ Bacteroidia_ Flavobacteriaceae_ NS2b_marine_group | 0.577 | 0.0069 |
| 2bc527ce1df7b1e131c5c9be089a7470 | Seawater: Aug. 2016 | Bacteroidota_ Bacteroidia_ Flavobacteriaceae_ NS4_marine_group | 0.767 | 0.0006 |
| 632921f144d05dc881ae3c80c9191efa | Seawater: Aug. 2016 | Bacteroidota_ Bacteroidia_ Flavobacteriaceae_ NS4_marine_group | 0.746 | 0.0004 |
| 427b763780d5250f242448ab5a46ac83 | Seawater: Aug. 2016 | Bacteroidota_ Bacteroidia_ Flavobacteriaceae_ NS4_marine_group | 0.695 | 0.0002 |
| dfd33a21cbd4de7266ee1ec8eeffc0c5 | Seawater: Aug. 2016 | Bacteroidota_ Bacteroidia_ Flavobacteriaceae_ NS4_marine_group | 0.632 | 0.0096 |
| ab848ca711def86b9a6b34751cd54961 | Seawater: Aug. 2016 | Bacteroidota_ Bacteroidia_ Flavobacteriaceae_ NS4_marine_group | 0.63 | 0.0006 |
| 01dfff5483186d1610dc263a6db41ffb | Seawater: Aug. 2016 | Bacteroidota_ Bacteroidia_ Flavobacteriaceae_ NS4_marine_group | 0.577 | 0.01 |
| 53b93714df5b9c838f8c5ec94c31b0b4 | Seawater: Aug. 2016 | Bacteroidota_ Bacteroidia_ Flavobacteriaceae_ NS4_marine_group | 0.556 | 0.0057 |
| 1b9e952f455565d835cb087ecb490dbf | Seawater: Aug. 2016 | Bacteroidota_ Bacteroidia_ Flavobacteriaceae_ NS4_marine_group | 0.545 | 0.0001 |
| 9d497f6d28fe7e3455e100596ec3bbe2 | Seawater: Aug. 2016 | Bacteroidota_ Bacteroidia_ Flavobacteriaceae_ NS4_marine_group | 0.529 | 0.0006 |
| db59f44cbf635f24cfe4a7192a4a2a88 | Seawater: Aug. 2016 | Bacteroidota_ Bacteroidia_ Flavobacteriaceae_ NS5_marine_group | 0.632 | 0.0069 |
| 730d48bd720ccdcb91016c8f8b9f269a | Seawater: Aug. 2016 | Bacteroidota_ Bacteroidia_ Flavobacteriaceae_ NS5_marine_group | 0.543 | 0.012 |
| a6481ea882bc30cc797d9641fb35b1a8 | Seawater: Aug. 2016 | Bacteroidota_ Bacteroidia_ Flavobacteriaceae_ NS5_marine_group | 0.535 | 0.0125 |
| 4da6179f32abb07b20d045d4f50d62ec | Seawater: Aug. 2016 | Bacteroidota_ Bacteroidia_ Flavobacteriaceae_ NS5_marine_group | 0.529 | 0.0069 |
| d8f1d6e909f880f987a28869da798b4b | Seawater: Aug. 2016 | Bacteroidota_ Bacteroidia_ Flavobacteriaceae_ NS5_marine_group | 0.498 | 0.0084 |
| 4572a73fea626ed66ebcafc52a9dacbf | Seawater: Aug. 2016 | Bacteroidota_ Bacteroidia_ Flavobacteriaceae_ NS5_marine_group | 0.459 | 0.0084 |
| 7a25f704cbab3504e659ed1a5c675ef9 | Seawater: Aug. 2016 | Bacteroidota_ Bacteroidia_ Flavobacteriaceae_ NS5_marine_group | 0.456 | 0.0458 |
| 67679fd25af82b2dcd1977585093a7dc | Seawater: Aug. 2016 | Bacteroidota_ Bacteroidia_ Flavobacteriaceae_ uncultured | 0.529 | 0.0079 |
| ce7c00f38cf40c99226d09ffe8dc18fd | Seawater: Aug. 2016 | Bacteroidota_ Bacteroidia_ NS7_marine_group_ NS7_marine_group | 0.587 | 0.0137 |
| 221a0cbbaffc326428d0eee22ccfdf0b | Seawater: Aug. 2016 | Bacteroidota_ Bacteroidia_ NS9_marine_group_ NS9_marine_group | 0.656 | 0.0007 |
| 2c2bc659f3e4b05a040e5a85db470189 | Seawater: Aug. 2016 | Bacteroidota_ Bacteroidia_ NS9_marine_group_ NS9_marine_group | 0.484 | 0.0347 |
| dbce81b5131df3e2bb42b36f1f198d6e | Seawater: Aug. 2016 | Bacteroidota_ Bacteroidia_ Saprospiraceae_ uncultured | 0.504 | 0.0095 |
| 8c1853b896032fa153d0d09e20985100 | Seawater: Aug. 2016 | Bacteroidota_ Bacteroidia_ uncultured_ uncultured | 0.829 | 0.0001 |
| ea91cf5f16019e6f11ee82c17b8895e2 | Seawater: Aug. 2016 | Bacteroidota_ Rhodothermia_ Balneolaceae_ Balneola | 0.619 | 0.0001 |
| 926b6c77c03122f9d321f6e431fbafb1 | Seawater: Aug. 2016 | Bdellovibrionota_ Bdellovibrionia_ Bacteriovoracaceae_ uncultured | 0.77 | 0.0005 |
| 227911db0c5e1780427d348f31541d01 | Seawater: Aug. 2016 | Bdellovibrionota_ Bdellovibrionia_ Bacteriovoracaceae_ uncultured | 0.425 | 0.0219 |
| daec6d612436dda037ec8831128b3c8a | Seawater: Aug. 2016 | Bdellovibrionota_ Bdellovibrionia_ Bdellovibrionaceae_ OM27 clade | 0.739 | 0.0002 |
| 203233cc6815855ac4f4e440be518398 | Seawater: Aug. 2016 | Bdellovibrionota_ Bdellovibrionia_ Bdellovibrionaceae_ OM27 clade | 0.723 | 0.0018 |

| | | | | |
|---|---|---|---|---|
| f60e44fd7dfef4ad1294858904d48c06 | Seawater: Aug. 2016 | Bdellovibrionota_ Bdellovibrionia_ Bdellovibrionaceae_ OM27_clade | 0.594 | 0.0069 |
| 3fee6f873beffcb66dffafeb932bcfb8 | Seawater: Aug. 2016 | Cyanobacteria_ Cyanobacteriia_ Cyanobiaceae_ Cyanobium | 0.673 | 0.0006 |
| 5edffc711a125e132bacefc7d1cebcdc | Seawater: Aug. 2016 | Cyanobacteria_ Cyanobacteriia_ Cyanobiaceae_ Prochlorococcus | 0.696 | 0.0002 |
| 6b552e93793448fc89916a5acbfe450c | Seawater: Aug. 2016 | Cyanobacteria_ Cyanobacteriia_ Cyanobiaceae_ Prochlorococcus | 0.508 | 0.0194 |
| f3e6d546c65a70fad100a063f5059453 | Seawater: Aug. 2016 | Cyanobacteria_ Cyanobacteriia_ Cyanobiaceae_ Prochlorococcus | 0.475 | 0.0145 |
| 5914f7175c14d32732e4b3bb91aef0ba | Seawater: Aug. 2016 | Cyanobacteria_ Cyanobacteriia_ Cyanobiaceae_ Prochlorococcus | 0.444 | 0.0312 |
| 6e087899941182e34b6278aa1719f73f | Seawater: Aug. 2016 | Cyanobacteria_ Cyanobacteriia_ Cyanobiaceae_ Prochlorococcus | 0.399 | 0.0378 |
| 7f46445bba0258b68f8b27cbe7ca8fe3 | Seawater: Aug. 2016 | Cyanobacteria_ Cyanobacteriia_ Cyanobiaceae_ Synechococcus | 0.923 | 0.0001 |
| 96faaf8c4eeb6edf11524c726e36b935 | Seawater: Aug. 2016 | Cyanobacteria_ Cyanobacteriia_ Cyanobiaceae_ Synechococcus | 0.757 | 0.0009 |
| 7410ff0ed5c0811ef3086f75d06a7309 | Seawater: Aug. 2016 | Cyanobacteria_ Cyanobacteriia_ Cyanobiaceae_ Synechococcus | 0.57 | 0.0042 |
| 161d69085f963c2394b49851d20a260c | Seawater: Aug. 2016 | Cyanobacteria_ Cyanobacteriia_ Phormidiaceae_ Trichodesmium | 0.548 | 0.0123 |
| cfe646e2046d11ae525b84f29c83227b | Seawater: Aug. 2016 | Desulfobacterota_ Desulfuromonadia_ Bradymonadales_ Bradymonadales | 0.632 | 0.0077 |
| ef639fc13fb36391eee63411c474fefc | Seawater: Aug. 2016 | Desulfobacterota_ Desulfuromonadia_ Bradymonadales_ Bradymonadales | 0.594 | 0.0077 |
| f3e6d9afd6faa5d47483d633758e22a8 | Seawater: Aug. 2016 | Desulfobacterota_ Desulfuromonadia_ PB19_ PB19 | 0.738 | 0.0002 |
| 202378eae0a463daffeb262db116b48e | Seawater: Aug. 2016 | Desulfobacterota_ Desulfuromonadia_ PB19_ PB19 | 0.625 | 0.0096 |
| 35b8663b925b92d83380844a13feaba0 | Seawater: Aug. 2016 | Margulisbacteria_ Margulisbacteria_ Margulisbacteria_ Margulisbacteria | 0.71 | 0.0006 |
| 16e9bbdf2cd1b2b99154c7ea4f7aa156 | Seawater: Aug. 2016 | Margulisbacteria_ Margulisbacteria_ Margulisbacteria_ Margulisbacteria | 0.667 | 0.0002 |
| 74e304a41a134b33146bd6644df374f9 | Seawater: Aug. 2016 | Patescibacteria_ Gracilibacteria_ Candidatus_Peregrinibacteria_Peregrinibacteria | 0.594 | 0.0084 |
| 5b15c060a0400be217f6c249fa13ad01 | Seawater: Aug. 2016 | Planctomycetota_ Planctomycetes_ Pirellulaceae_ uncultured | 0.501 | 0.0237 |
| b7dac74135c427ac12a409a48fc417c4 | Seawater: Aug. 2016 | Proteobacteria_ Alphaproteobacteria_ AEGEAN-169 marine group | 0.768 | 0.0006 |
| 29ca6614cc134e6a067dfc5151f9e098 | Seawater: Aug. 2016 | Proteobacteria_ Alphaproteobacteria_ AEGEAN-169 marine group | 0.731 | 0.0005 |
| 36a5812b371dfd6996703f8e33070c42 | Seawater: Aug. 2016 | Proteobacteria_ Alphaproteobacteria_ AEGEAN-169 marine group | 0.68 | 0.0018 |
| 77cb93a811acc96bee76d3f9e1090ecc | Seawater: Aug. 2016 | Proteobacteria_ Alphaproteobacteria_ AEGEAN-169 marine group | 0.671 | 0.0003 |
| b4d270282eb787c97eec8c5d34462cfa | Seawater: Aug. 2016 | Proteobacteria_ Alphaproteobacteria_ AEGEAN-169 marine group | 0.65 | 0.0004 |
| ecb7bec5f17246b33f5a2ad6a23a2f11 | Seawater: Aug. 2016 | Proteobacteria_ Alphaproteobacteria_ AEGEAN-169 marine group | 0.618 | 0.0069 |
| 112a22205015d4e0d7aa0be7b89a5678 | Seawater: Aug. 2016 | Proteobacteria_ Alphaproteobacteria_ AEGEAN-169 marine group | 0.616 | 0.0058 |
| 6e08db9fa0d10596303c6f3a7564ee0c | Seawater: Aug. 2016 | Proteobacteria_ Alphaproteobacteria_ AEGEAN-169 marine group | 0.443 | 0.0387 |
| 95aad3eab8694c19b34ac83d77e3a229 | Seawater: Aug. 2016 | Proteobacteria_ Alphaproteobacteria_ Clade II_ Clade II | 0.713 | 0.0022 |
| 676f2e73d996e5fb4f3bf7323a5b680d | Seawater: Aug. 2016 | Proteobacteria_ Alphaproteobacteria_ Clade_I_ Clade_Ia | 0.603 | 0.0069 |
| b8b5780c1ce726c1079482ca708a4c67 | Seawater: Aug. 2016 | Proteobacteria_ Alphaproteobacteria_ Clade_I_ Clade_Ia | 0.576 | 0.0116 |
| ad4a4a1ae048e63338a8578db0bd6067 | Seawater: Aug. 2016 | Proteobacteria_ Alphaproteobacteria_ Clade_I_ Clade_Ib | 0.618 | 0.0069 |
| 6018223b27150015283a87fa2c664ee7 | Seawater: Aug. 2016 | Proteobacteria_ Alphaproteobacteria_ Clade_I_ Clade_Ib | 0.594 | 0.0069 |
| 5e2c0141835a486186d1ec0bd92f553d | Seawater: Aug. 2016 | Proteobacteria_ Alphaproteobacteria_ Clade_I_ Clade_Ib | 0.573 | 0.0069 |
| 7d8d34ec1a6d9902eb23ace5ddd63a5e | Seawater: Aug. 2016 | Proteobacteria_ Alphaproteobacteria_ Clade_I_ Clade_Ib | 0.503 | 0.0225 |
| c8845f01025917fda00850ae3a695f99 | Seawater: Aug. 2016 | Proteobacteria_ Alphaproteobacteria_ Clade_I_ Clade_Ib | 0.475 | 0.0361 |
| 7c5b07a92a8830cecf821c75f4bc32c0 | Seawater: Aug. 2016 | Proteobacteria_ Alphaproteobacteria_ Clade_II_ Clade_II | 0.754 | 0.0006 |
| 3947e817bf6281b4e7fcd794c26b2769 | Seawater: Aug. 2016 | Proteobacteria_ Alphaproteobacteria_ Clade_II_ Clade_II | 0.696 | 0.001 |
| b8f356c93ef1ecb66fcc0d5cb25384ce | Seawater: Aug. 2016 | Proteobacteria_ Alphaproteobacteria_ Clade_II_ Clade_II | 0.64 | 0.0018 |
| 897876fbdc53a5f77c3a4fc1b44cc943 | Seawater: Aug. 2016 | Proteobacteria_ Alphaproteobacteria_ Clade_II_ Clade_II | 0.629 | 0.0006 |
| 4160a089ef938b7157fdb362d6194216 | Seawater: Aug. 2016 | Proteobacteria_ Alphaproteobacteria_ Clade_II_ Clade_II | 0.615 | 0.0006 |

| | | | | |
|---|---|---|---|---|
| 13b6779e2ea6f6c51c9a136c5848b37d | Seawater: Aug. 2016 | Proteobacteria_ Alphaproteobacteria_ Clade_II_ Clade_II | 0.614 | 0.0034 |
| c3ec53c187585b01b87e33ef59ace59b | Seawater: Aug. 2016 | Proteobacteria_ Alphaproteobacteria_ Clade_II_ Clade_II | 0.517 | 0.0006 |
| f62fdf744d68a68fc7b4b416f9b4ea1c | Seawater: Aug. 2016 | Proteobacteria_ Alphaproteobacteria_ Clade_II_ Clade_II | 0.505 | 0.0201 |
| cd82cffffa5a272dde88382feb1799ad | Seawater: Aug. 2016 | Proteobacteria_ Alphaproteobacteria_ Clade_II_ Clade_II | 0.45 | 0.0325 |
| 317a810c7803461cae39a687fe4283bf | Seawater: Aug. 2016 | Proteobacteria_ Alphaproteobacteria_ Clade_II_ Clade_II | 0.426 | 0.0455 |
| 6de40fcdea2aa22c95bc4f5b856c9b03 | Seawater: Aug. 2016 | Proteobacteria_ Alphaproteobacteria_ Clade_III_ Clade_III | 0.607 | 0.0003 |
| a2ca8b84937888088b211c6857de77bf | Seawater: Aug. 2016 | Proteobacteria_ Alphaproteobacteria_ Clade_III_ Clade_III | 0.541 | 0.0006 |
| c2174a26077a12e699443390bd1913ff | Seawater: Aug. 2016 | Proteobacteria_ Alphaproteobacteria_ Hyphomicrobiaceae_ Filomicrobium | 0.594 | 0.0079 |
| 25e397704f7154057ff2f242ed7428b9 | Seawater: Aug. 2016 | Proteobacteria_ Alphaproteobacteria_ Parvibaculaceae_ uncultured | 0.701 | 0.0003 |
| 001e7b36aee068d2f87b4b8b627352fb | Seawater: Aug. 2016 | Proteobacteria_ Alphaproteobacteria_ PS1_clade_ PS1_clade | 0.573 | 0.0003 |
| f8ee7243228c15925e677b5d588d6cac | Seawater: Aug. 2016 | Proteobacteria_ Alphaproteobacteria_ PS1_clade_ PS1_clade | 0.531 | 0.0138 |
| 2c1abae7a17234303c2e2e1547258a3e | Seawater: Aug. 2016 | Proteobacteria_ Alphaproteobacteria_ Rhodobacteraceae_ Ascidiaceihabitans | 0.531 | 0.0195 |
| 3418c640e8c753c7053bf04b152a3c9b | Seawater: Aug. 2016 | Proteobacteria_ Alphaproteobacteria_ Rhodobacteraceae_NA | 0.76 | 0.0006 |
| cc31f1ecc1714e03edfa64901ffb3fe9 | Seawater: Aug. 2016 | Proteobacteria_ Alphaproteobacteria_ Rhodobacteraceae_NA | 0.73 | 0.0006 |
| 3ff3e2b7713aafa598d5df3bb886f8ad | Seawater: Aug. 2016 | Proteobacteria_ Alphaproteobacteria_ Rhodobacteraceae_NA | 0.626 | 0.0069 |
| 22941735d1f5080614010e969a7949bd | Seawater: Aug. 2016 | Proteobacteria_ Alphaproteobacteria_ S25-593_ S25-593 | 0.754 | 0.0006 |
| 13a5ae4c5b1402ea51db6a6d88626b83 | Seawater: Aug. 2016 | Proteobacteria_ Alphaproteobacteria_ S25-593_ S25-593 | 0.678 | 0.0026 |
| d7a30d9fe61b0f29f3d657a1bd4b3118 | Seawater: Aug. 2016 | Proteobacteria_ Alphaproteobacteria_ S25-593_ S25-593 | 0.672 | 0.0006 |
| 35c8271539edb716f697ef9047b29fa5 | Seawater: Aug. 2016 | Proteobacteria_ Alphaproteobacteria_ S25-593_ S25-593 | 0.647 | 0.0012 |
| 9a7666ef2536b9aba55360de92737541 | Seawater: Aug. 2016 | Proteobacteria_ Alphaproteobacteria_ S25-593_ S25-593 | 0.594 | 0.0084 |
| 6e4adc38d605e6c1c2295b460abab9c5 | Seawater: Aug. 2016 | Proteobacteria_ Alphaproteobacteria_ SAR116_clade_ Candidatus_Puniceispirillum | 0.632 | 0.0084 |
| 2c9c95d43c4be9c1f9cc9c795e22fec9 | Seawater: Aug. 2016 | Proteobacteria_ Alphaproteobacteria_ SAR116_clade_ SAR116_clade | 0.82 | 0.0002 |
| 2ea223f5051d449015e533ea2091e896 | Seawater: Aug. 2016 | Proteobacteria_ Alphaproteobacteria_ SAR116_clade_ SAR116_clade | 0.816 | 0.0006 |
| ceaea439b55965816ae4eb9b308f0000 | Seawater: Aug. 2016 | Proteobacteria_ Alphaproteobacteria_ SAR116_clade_ SAR116_clade | 0.787 | 0.0004 |
| 4ee88997788e13e61782af0af8907edc | Seawater: Aug. 2016 | Proteobacteria_ Alphaproteobacteria_ SAR116_clade_ SAR116_clade | 0.78 | 0.0006 |
| f19c9351558d568b2e228f6a8d7dba2f | Seawater: Aug. 2016 | Proteobacteria_ Alphaproteobacteria_ SAR116_clade_ SAR116_clade | 0.612 | 0.0059 |
| 1e578e5b88a8176a26e797deb74bcebd | Seawater: Aug. 2016 | Proteobacteria_ Alphaproteobacteria_ SAR116_clade_ SAR116_clade | 0.569 | 0.0006 |
| cc0df71b999fbaff1cfe19cdf300dfaa | Seawater: Aug. 2016 | Proteobacteria_ Alphaproteobacteria_ SAR116_clade_ SAR116_clade | 0.54 | 0.0084 |
| b1eee9d661c00ba11054e790f10e9dd9 | Seawater: Aug. 2016 | Proteobacteria_ Alphaproteobacteria_ Sphingomonadaceae_ Sphingopyxis | 0.632 | 0.0069 |
| cd58184a123a4ee0ca31017fb9667b51 | Seawater: Aug. 2016 | Proteobacteria_ Alphaproteobacteria_ Stappiaceae_ Stappiaceae | 0.465 | 0.0279 |
| bc6903d0064c36e190010c8cae4b70ff | Seawater: Aug. 2016 | Proteobacteria_ Alphaproteobacteria_ Stappiaceae_ Stappiaceae | 0.46 | 0.0157 |
| abf1e9f186acd0a3d3f2a946e7c379c8 | Seawater: Aug. 2016 | Proteobacteria_ Alphaproteobacteria_ uncultured_ uncultured | 0.705 | 0.0019 |
| 8696e6b42645982923ff0e9cb25cfc79 | Seawater: Aug. 2016 | Proteobacteria_ Alphaproteobacteria_ uncultured_ uncultured | 0.613 | 0.0069 |
| 7c0df59f3ea7461669e41a69b9223ae7 | Seawater: Aug. 2016 | Proteobacteria_ Alphaproteobacteria_ uncultured_ uncultured | 0.505 | 0.0138 |
| f406b9bc61936a51699db1b8631b8369 | Seawater: Aug. 2016 | Proteobacteria_ Alphaproteobacteria_ uncultured_ uncultured | 0.477 | 0.0123 |
| 9f7df1b9ce535af7d8dd74499d7cf303 | Seawater: Aug. 2016 | Proteobacteria_ Gammaproteobacteria_ Alcanivoracaceae1_ Alcanivorax | 0.557 | 0.0006 |
| 7668a30438c5fdeb5b09f8e925470267 | Seawater: Aug. 2016 | Proteobacteria_ Gammaproteobacteria_ Coxiellaceae_ Coxiella | 0.718 | 0.0002 |
| e3e04d81fe275fcbee4980f42f215f99 | Seawater: Aug. 2016 | Proteobacteria_ Gammaproteobacteria_ Coxiellaceae_ Coxiella | 0.632 | 0.0096 |
| 942226eaad2f068ad85dd45d24cea154 | Seawater: Aug. 2016 | Proteobacteria_ Gammaproteobacteria_ Coxiellaceae_ Coxiella | 0.618 | 0.0069 |
| 1c8387132e76cf9185e079ba46d29746 | Seawater: Aug. 2016 | Proteobacteria_ Gammaproteobacteria_ Coxiellaceae_ Coxiella | 0.618 | 0.0069 |

| | | | | |
|---|---|---|---|---|
| 7e865b1e993e304ad06c2aa21cefa687 | Seawater: Aug. 2016 | Proteobacteria_ Gammaproteobacteria_ Coxiellaceae_ Coxiella | 0.561 | 0.0078 |
| aa9d7ab25ecda0fedb1d785a205af9b4 | Seawater: Aug. 2016 | Proteobacteria_ Gammaproteobacteria_ Coxiellaceae_ Coxiella | 0.529 | 0.0084 |
| 9b77b69418e26a604e6721ffb7ed26f0 | Seawater: Aug. 2016 | Proteobacteria_ Gammaproteobacteria_ Coxiellaceae_ Coxiella | 0.529 | 0.0069 |
| 991c3081ddba2a5e05920de4de4a63a7 | Seawater: Aug. 2016 | Proteobacteria_ Gammaproteobacteria_ Ectothiorhodospiraceae_ uncultured | 0.69 | 0.0001 |
| c90aa24db63ee378a69fd05a9c17c349 | Seawater: Aug. 2016 | Proteobacteria_ Gammaproteobacteria_ Halieaceae_ OM60(NOR5)_clade | 0.768 | 0.0005 |
| df6f5330b944baf3440591d76b313909 | Seawater: Aug. 2016 | Proteobacteria_ Gammaproteobacteria_ Halieaceae_ OM60(NOR5)_clade | 0.756 | 0.0006 |
| 1895163381cb08cb9849606686b8a0c3 | Seawater: Aug. 2016 | Proteobacteria_ Gammaproteobacteria_ Halieaceae_ OM60(NOR5)_clade | 0.635 | 0.0029 |
| 7987b494778fb929aebb3043bbbcaf07 | Seawater: Aug. 2016 | Proteobacteria_ Gammaproteobacteria_ Halieaceae_ OM60(NOR5)_clade | 0.594 | 0.0069 |
| a6b8f9f050228c36abe4492882a11f27 | Seawater: Aug. 2016 | Proteobacteria_ Gammaproteobacteria_ Halieaceae_ OM60(NOR5)_clade | 0.579 | 0.0062 |
| c92417979ba135060ce865a0d20e769a | Seawater: Aug. 2016 | Proteobacteria_ Gammaproteobacteria_ Halieaceae_ OM60(NOR5)_clade | 0.487 | 0.0006 |
| 443ad648580773ee6053790f3a2de0c3 | Seawater: Aug. 2016 | Proteobacteria_ Gammaproteobacteria_ KI89A_clade_ KI89A_clade | 0.511 | 0.0253 |
| 9d879b65b9db1ee1e912eb4191740f64 | Seawater: Aug. 2016 | Proteobacteria_ Gammaproteobacteria_ Litoricolaceae_ Litoricola | 0.745 | 0.0008 |
| a49f66e342e722cad630deb65ea82657 | Seawater: Aug. 2016 | Proteobacteria_ Gammaproteobacteria_ Litoricolaceae_ Litoricola | 0.735 | 0.0005 |
| 24bbc31940709f3fe2b5ceeb025487ea | Seawater: Aug. 2016 | Proteobacteria_ Gammaproteobacteria_ Litoricolaceae_ Litoricola | 0.661 | 0.0017 |
| 3bd5385cab1346abb973cbfe6d9f0bdb | Seawater: Aug. 2016 | Proteobacteria_ Gammaproteobacteria_ Marinomonadaceae_ Marinomonas | 0.541 | 0.0107 |
| 0c611f056e8c436c0b9ea766b03fa067 | Seawater: Aug. 2016 | Proteobacteria_ Gammaproteobacteria_ Methylophagaceae_ Methylophaga | 0.618 | 0.0084 |
| b74a6dea408fbfdc09e37b0bf5049fdc | Seawater: Aug. 2016 | Proteobacteria_ Gammaproteobacteria_ MWH-UniP1 aquatic group_NA | 0.524 | 0.0006 |
| 7c3f4427da7952dbb02fe3e00f7ce7aa | Seawater: Aug. 2016 | Proteobacteria_ Gammaproteobacteria_ MWH-UniP1_aquatic_group_NA | 0.593 | 0.0048 |
| e05e4f085b96829763ee7e0753b13774 | Seawater: Aug. 2016 | Proteobacteria_ Gammaproteobacteria_ Porticoccaceae_ SAR92 clade | 0.733 | 0.0004 |
| 7df2590093e2a8d482ccb4d73f49a29a | Seawater: Aug. 2016 | Proteobacteria_ Gammaproteobacteria_ Porticoccaceae_ SAR92_clade | 0.675 | 0.0005 |
| 4b0160f7ccc7c2b03767d35aeb6ebebf | Seawater: Aug. 2016 | Proteobacteria_ Gammaproteobacteria_ SAR86_clade_ SAR86 clade | 0.739 | 0.0004 |
| dcd60c368bddfc09116fca502d17acd0 | Seawater: Aug. 2016 | Proteobacteria_ Gammaproteobacteria_ SAR86_clade_ SAR86 clade | 0.727 | 0.0006 |
| c305ab9020c2c5135e3a10aa1c7f7c7b | Seawater: Aug. 2016 | Proteobacteria_ Gammaproteobacteria_ SAR86_clade_ SAR86_clade | 0.814 | 0.0002 |
| 9ec997c700d09a502a482a43b707b85c | Seawater: Aug. 2016 | Proteobacteria_ Gammaproteobacteria_ SAR86_clade_ SAR86_clade | 0.7 | 0.0002 |
| 9442980515ce7f911bb4cc9c372db4e8 | Seawater: Aug. 2016 | Proteobacteria_ Gammaproteobacteria_ SAR86_clade_ SAR86_clade | 0.669 | 0.0006 |
| 5ade5e6e170fc840b051e6db9b20f6b5 | Seawater: Aug. 2016 | Proteobacteria_ Gammaproteobacteria_ SAR86_clade_ SAR86_clade | 0.613 | 0.0061 |
| 8be065ce4c492fc4e455e39f8cb0df09 | Seawater: Aug. 2016 | Proteobacteria_ Gammaproteobacteria_ SAR86_clade_ SAR86_clade | 0.604 | 0.0069 |
| a901a446072ee210c63413d49a4aecce | Seawater: Aug. 2016 | Proteobacteria_ Gammaproteobacteria_ SAR86_clade_ SAR86_clade | 0.56 | 0.0001 |
| 1fb75bc033bf8be2ebe6ce8cc3ba0147 | Seawater: Aug. 2016 | Proteobacteria_ Gammaproteobacteria_ SAR86_clade_ SAR86_clade | 0.558 | 0.0141 |
| 703bba9790f220e543ef8a292ec80e20 | Seawater: Aug. 2016 | Proteobacteria_ Gammaproteobacteria_ SAR86_clade_ SAR86_clade | 0.556 | 0.0004 |
| edd413da17e10fa2488c6bf58fc8f714 | Seawater: Aug. 2016 | Proteobacteria_ Gammaproteobacteria_ SAR86_clade_ SAR86_clade | 0.545 | 0.0039 |
| c360ebf606bc22eacf074d3e28e0628d | Seawater: Aug. 2016 | Proteobacteria_ Gammaproteobacteria_ SAR86_clade_ SAR86_clade | 0.533 | 0.0129 |
| a78991c0e76742c13f02d877ae85ac39 | Seawater: Aug. 2016 | Proteobacteria_ Gammaproteobacteria_ Spongiibacteraceae_ BD1-7_clade | 0.498 | 0.0077 |
| e99730501d8a2d0639cedc50419fecb7 | Seawater: Aug. 2016 | Proteobacteria_ Gammaproteobacteria_ Vibrionaceae_ uncultured | 0.51 | 0.0327 |
| 86845709330d300cda73dc215dd6cc7f | Seawater: Aug. 2016 | Proteobacteria_ Gammaproteobacteria_NA_NA | 0.617 | 0.0058 |
| 2c90976a6d73ec89c850b6b8771ee571 | Seawater: Aug. 2016 | SAR324 clade_Marine group B_ NA_NA | 0.574 | 0.0013 |
| cfd23b4c97ae6f15e8b9717f7ff95405 | Seawater: Aug. 2016 | SAR324 clade_Marine group B_ NA_NA | 0.555 | 0.0077 |
| bd6edf25c3a3b570f9f471f9b658ea77 | Seawater: Aug. 2016 | Verrucomicrobiota_ Chlamydiae_ Chlamydiaceae_ uncultured | 0.632 | 0.0079 |
| 0dcdc65fc96d866ebec6d3c29a2ed307 | Seawater: Aug. 2016 | Verrucomicrobiota_ Kiritimatiellae_ Kiritimatiellaceae_ R76-B128 | 0.447 | 0.0079 |
| 9a57903995bfe6cd19c94eba09d7c966 | Seawater: Aug. 2016 | Verrucomicrobiota_ Verrucomicrobiae_ Pedosphaeraceae_ SCGC_AAA164-E04 | 0.764 | 0.0006 |

| | | | | |
|---|---|---|---|---|
| 3d35e642d296196619f455c5f3694260 | Seawater: Aug. 2016 | Verrucomicrobiota_ Verrucomicrobiae_ Puniceicoccaceae_ Coraliomargarita | 0.849 | 0.0002 |
| 5cdfaf51ed18b88b6f3f2aca0fe6a206 | Seawater: Aug. 2016 | Verrucomicrobiota_ Verrucomicrobiae_ Puniceicoccaceae_ Coraliomargarita | 0.7 | 0.001 |
| 3c053526c5b09f880a975cec099fef25 | Seawater: Aug. 2016 | Verrucomicrobiota_ Verrucomicrobiae_ Puniceicoccaceae_ Lentimonas | 0.763 | 0.0001 |
| 01b8c125e5158fa67f3b6c9e6d263431 | Seawater: Aug. 2016 | WPS-2_ WPS-2_ WPS-2_ WPS-2 | 0.618 | 0.0079 |
| 9a4abf5ed8f0ff8eec580004a028dfea | Seawater: Aug. 2016 | Marinimicrobia_SAR406 clade_NA_NA | 0.447 | 0.0227 |
| 742574ffa44ab2b37a00cff90020e045 | Seawater: Oct. 2017 | Actinobacteriota_ Acidimicrobiia_ Microtrichaceae_ Sva0996 marine group | 0.586 | 0.0114 |
| 163fd46ec09dfe3bd19184f624dab6af | Seawater: Oct. 2017 | Bacteroidota_ Bacteroidia_ Flavobacteriaceae_ NS2b_marine_group | 0.556 | 0.0167 |
| 6509395ba6d1619a35b3d815623fe234 | Seawater: Oct. 2017 | Bacteroidota_ Bacteroidia_ Flavobacteriaceae_ NS4_marine_group | 0.477 | 0.0379 |
| 3ef18d3db2ad87853d22a74048de8af0 | Seawater: Oct. 2017 | Bacteroidota_ Bacteroidia_ NS9_marine_group_ NS9_marine_group | 0.509 | 0.0273 |
| f3bdcc5040fcac34c18f947ea085debf | Seawater: Oct. 2017 | Bdellovibrionota_ Bdellovibrionia_ Bacteriovoracaceae_ uncultured | 0.578 | 0.0097 |
| 1c492390008157183ff827649a8530e4 | Seawater: Oct. 2017 | Bdellovibrionota_ Bdellovibrionia_ Bdellovibrionaceae_ OM27_clade | 0.477 | 0.042 |
| 88310351fcc1f3ce61d05c4cedfc777b | Seawater: Oct. 2017 | Bdellovibrionota_ Oligoflexia_ uncultured_ uncultured | 0.536 | 0.0155 |
| 99b84be2fb41895e181a82f1f769d601 | Seawater: Oct. 2017 | Bdellovibrionota_ Oligoflexia_ uncultured_ uncultured | 0.473 | 0.0355 |
| 94adb88b4dc27697115db4fb98e4a4d3 | Seawater: Oct. 2017 | Chloroflexi_ Dehalococcoidia_ SAR202_clade_ SAR202_clade | 0.651 | 0.0039 |
| 8212b33fb820f18cb107b5cc5b05e93c | Seawater: Oct. 2017 | Chloroflexi_ Dehalococcoidia_ SAR202_clade_ SAR202_clade | 0.546 | 0.0286 |
| 9217ee2f86049e987b872de6e596448a | Seawater: Oct. 2017 | Chloroflexi_ Dehalococcoidia_ SAR202_clade_ SAR202_clade | 0.489 | 0.032 |
| 897a323aabb5494fe95da8745bf2f149 | Seawater: Oct. 2017 | Dadabacteria_ Dadabacteriia_ Dadabacteriales_ Dadabacteriales | 0.526 | 0.0229 |
| 8d46a1cbd43a7e900d6651eb70d44b38 | Seawater: Oct. 2017 | Desulfobacterota_ Desulfuromonadia_ Bradymonadales_ Bradymonadales | 0.533 | 0.023 |
| b721d5e2d9de745875a9c24bcc8316f1 | Seawater: Oct. 2017 | Gemmatimonadota_ BD2-11 terrestrial group_NA_NA | 0.488 | 0.026 |
| d21767ad8591e72b1d006cf3ada7d6f5 | Seawater: Oct. 2017 | Margulisbacteria_ Margulisbacteria_ Margulisbacteria_ Margulisbacteria | 0.506 | 0.0341 |
| 186a55f35bf39b44454380339c950209 | Seawater: Oct. 2017 | Nanoarchaeota_ Nanoarchaeia_ Woesearchaeales_ Woesearchaeales | 0.65 | 0.0037 |
| cc08b0d91a99730c1f59b56b959af16c | Seawater: Oct. 2017 | NB1-j_ NB1-j_ NB1-j_ NB1-j | 0.511 | 0.0288 |
| f87820ab1694993e261b7f30de7d6e99 | Seawater: Oct. 2017 | Planctomycetota_ OM190_ OM190_ OM190 | 0.524 | 0.0215 |
| 26bbb4a8022d6374d37f9d3147a6d513 | Seawater: Oct. 2017 | Planctomycetota_ Planctomycetes_ Pirellulaceae_ Blastopirellula | 0.583 | 0.0071 |
| cd0283e6687d70df913b1bc1fd730133 | Seawater: Oct. 2017 | Planctomycetota_ Planctomycetes_ Pirellulaceae_ Blastopirellula | 0.526 | 0.025 |
| c12c6a15f208df5992ed2c7d14fd2a12 | Seawater: Oct. 2017 | Planctomycetota_ Planctomycetes_ Pirellulaceae_ Rubripirellula | 0.601 | 0.009 |
| 401eac421946eea8324228f54fc7b490 | Seawater: Oct. 2017 | Planctomycetota_ Planctomycetes_ Pirellulaceae_ uncultured | 0.518 | 0.0299 |
| e62236eccfc9e91dcd9f682bb3b8a604 | Seawater: Oct. 2017 | Planctomycetota_ Planctomycetes_ Rubinisphaeraceae_ uncultured | 0.638 | 0.0041 |
| 30ed29e8b2a60a549e95d41a89e8a427 | Seawater: Oct. 2017 | Planctomycetota_ Planctomycetes_ Rubinisphaeraceae_ uncultured | 0.549 | 0.0113 |
| cba09c45336067692e1698ed5545cfbe | Seawater: Oct. 2017 | Proteobacteria_ Alphaproteobacteria_ AEGEAN-169 marine group | 0.637 | 0.0037 |
| c76587bb614bd683b14c821f1d1d3773 | Seawater: Oct. 2017 | Proteobacteria_ Alphaproteobacteria_ Clade_II_ Clade_II | 0.503 | 0.0296 |
| 3e500d5204cb228f285b60352d83f92f | Seawater: Oct. 2017 | Proteobacteria_ Alphaproteobacteria_ Clade_II_ Clade_II | 0.49 | 0.0307 |
| 5b29fd2172d3c580d5ccb27241e094c9 | Seawater: Oct. 2017 | Proteobacteria_ Alphaproteobacteria_ S25-593_ S25-593 | 0.631 | 0.0051 |
| 958c2f1d614d0b18d88fe13a25c69e91 | Seawater: Oct. 2017 | Proteobacteria_ Alphaproteobacteria_ SAR116_clade_ SAR116_clade | 0.562 | 0.0185 |
| 669073dbfc6f067119aa86d2c286a8a1 | Seawater: Oct. 2017 | Proteobacteria_ Alphaproteobacteria_ Sphingomonadaceae_ Erythrobacter | 0.443 | 0.0343 |
| 95bf6daa606070aa9ffeb37e2cfad46f | Seawater: Oct. 2017 | Proteobacteria_ Alphaproteobacteria_ uncultured_ uncultured | 0.834 | 0.0001 |
| 9617fd0f7534adca733d0f21c03166a8 | Seawater: Oct. 2017 | Proteobacteria_ Alphaproteobacteria_ uncultured_ uncultured | 0.615 | 0.0086 |
| 9df77a8e00ca159c7f1ec80a0cf1179d | Seawater: Oct. 2017 | Proteobacteria_ Alphaproteobacteria_ uncultured_ uncultured | 0.517 | 0.0271 |
| 25bd90a5d60556457264c1e1c1f7de50 | Seawater: Oct. 2017 | Proteobacteria_ Alphaproteobacteria_ uncultured_ uncultured | 0.49 | 0.0318 |
| 2b3de226c378f7cdcede54adb6a803aa | Seawater: Oct. 2017 | Proteobacteria_ Gammaproteobacteria_ Alteromonadaceae_ Alteromonas | 0.475 | 0.0243 |

| | | | | |
|---|---|---|---|---|
| f1dcb832d2bc6a8299d3eff2fad37027 | Seawater: Oct. 2017 | Proteobacteria_ Gammaproteobacteria_ KI89A_clade_ KI89A clade | 0.584 | 0.0114 |
| 0cbf4cea7d202c84e5fed0170a826e07 | Seawater: Oct. 2017 | Proteobacteria_ Gammaproteobacteria_ KI89A_clade_ KI89A_clade | 0.697 | 0.0017 |
| 42052d6ad85f2a53162430dddbe7da88 | Seawater: Oct. 2017 | Proteobacteria_ Gammaproteobacteria_ OM182_clade_ OM182_clade | 0.668 | 0.0042 |
| b5f9962e1f4bd60efe96664457d4782e | Seawater: Oct. 2017 | Proteobacteria_ Gammaproteobacteria_ Vibrionaceae_ Photobacterium | 0.586 | 0.0079 |
| 03a6db6b45fe0f9efdfd1558e468ae04 | Seawater: Oct. 2017 | Proteobacteria_ Gammaproteobacteria_ Woeseiaceae_ Woeseia | 0.579 | 0.0098 |
| c67fa05a1bf5423688103136ebddec57 | Seawater: Oct. 2017 | Proteobacteria_ Gammaproteobacteria_NA_NA | 0.532 | 0.0182 |
| d6a0558df2186bb75bcf8c0d935d00f7 | Seawater: Oct. 2017 | Thermoplasmatota_ Thermoplasmata_ Marine_Group_II_ Marine_Group_II | 0.725 | 0.001 |
| a3995cc050a34658922227ab8656d369 | Seawater: Oct. 2017 | Thermoplasmatota_ Thermoplasmata_ Marine_Group_II_ Marine_Group_II | 0.531 | 0.0209 |
| 8b3bd16f0719e160836c70fc3e39590e | Seawater: Oct. 2017 | Thermoplasmatota_ Thermoplasmata_ Marine_Group_II_ Marine_Group_II | 0.498 | 0.0311 |
| 7e6bcaaf0b6fb03520f9d1d0f01aa7d7 | Seawater: Oct. 2017 | Thermoplasmatota_ Thermoplasmata_ Marine_Group_II_ Marine_Group_II | 0.498 | 0.029 |
| c4c8ab11109af1f18c2f95dcc1209c03 | Seawater: Oct. 2017 | Thermoplasmatota_ Thermoplasmata_ Marine_Group_II_ Marine_Group_II | 0.471 | 0.0472 |
| 99d5f77e5994d3f043b756b8f5dbc815 | Seawater: Oct. 2017 | Thermoplasmatota_ Thermoplasmata_ Thermoplasmata_ Marine_Group_III | 0.677 | 0.0024 |
| 572f6a2959c808a72d0b76c6dad9e7c5 | Seawater: Oct. 2017 | Verrucomicrobiota_ Verrucomicrobiae_ Arctic97B-4 marine group_NA | 0.519 | 0.024 |
| 91c5f061cb2fb58c4e13122af2749ae1 | Seawater: Oct. 2017 | Verrucomicrobiota_ Verrucomicrobiae_ DEV007_ DEV007 | 0.781 | 0.0007 |
| a391966037962ad7a30e92b4cc19a488 | Seawater: Oct. 2017 | Verrucomicrobiota_ Verrucomicrobiae_ Pedosphaeraceae_ SCGC_AAA164-E04 | 0.513 | 0.028 |
| 9af00ae4ff45984e4885f22c4b501dd7 | Seawater: Oct. 2017 | Verrucomicrobiota_ Verrucomicrobiae_ Puniceicoccaceae_ A714019 | 0.577 | 0.0099 |
| 7f3c98ae531cffd14e75e76a05760a50 | Seawater: Oct. 2017 | Verrucomicrobiota_ Verrucomicrobiae_ Rubritaleaceae_ Roseibacillus | 0.478 | 0.0324 |
| 874359ce05108ae6d0eb15976b76f5cb | Seawater: Oct. 2017 | Marinimicrobia_SAR406 clade_NA_NA | 0.724 | 0.0004 |
| 585ca34ed5218b522df0705c0b1e12ac | Seawater: Oct. 2017 | Marinimicrobia_SAR406 clade_NA_NA | 0.536 | 0.0214 |

**Supplemental Table S7**. Bacterial strains, primers, and annealing conditions used for qPCR analysis.

| Target bacteria | Target Gene | Primer sequences | Annealing Temp (°C) | Amplicon Size (bp) | Standards | Reference |
|---|---|---|---|---|---|---|
| *Escherichia coli* | ybbW | 5'-TGA TTG GCA AAA TCT GGC CG-3'<br>5'-GAA ATC GCC CAA ATC GCC AT-3' | 60 | 210 | *E. coli* K12 ATCC 47076 | (1) |
| *Enterococcus* spp. | 23S rRNA | 5'-GAG AAA TTC CAA ACG AAC TTG-3'<br>5'-CAG TGC TCT ACC TCC ATC ATT-3' | 55 | 112 | *E. faecalis* ATCC 29212 | (2) |
| *Klebsiella pneumoniae* | phoE | 5'-TGC CCA GAC CGA TAA CTT TA-3'<br>5'-CTG TTT CTT CGC TTC ACG C-3' | 52 | 142 | *K. pneumoniae* ATCCBAA-1705 | (3) |
| *Pseudomonas aeruginosa* | oprI | 5'-ATG AAC AAC GTT CTG AAA TTC TCT GCT-3'<br>5'-CTT GCG GCT GGC TTT TTC CAG-3' | 57 | 249 | *P. aeruginosa* ATCCBAA-1744 | (4) |
| *Salmonella enterica* | hilA | 5'- GTG AAA TTA TCG CCA CGT TCG GGC AA-3'<br>5'-TCA TCG CAC CGT CAA AGG AAC C-3' | 60 | 284 | *S. enterica* ATCC 14028 | (5) |
| *Serratia marcescens* | luxS | 5'-TGC CTG GAA AGC GGC GAT GG-3'<br>5'-CGC CAG CTC GTC GTT GTG GT -3' | 62 | 174 | *S. marcescens* Db11 | (6) |
| *Staphylococcus aureus* | eap | 5'-TAC TAA CGA AGC ATC TGC C-3'<br>5'-TTA AAT CGA TAT CAC TAA TAC CTC-3' | 50 | 230 | *S. aureus* ATCC 29213 | (7) |

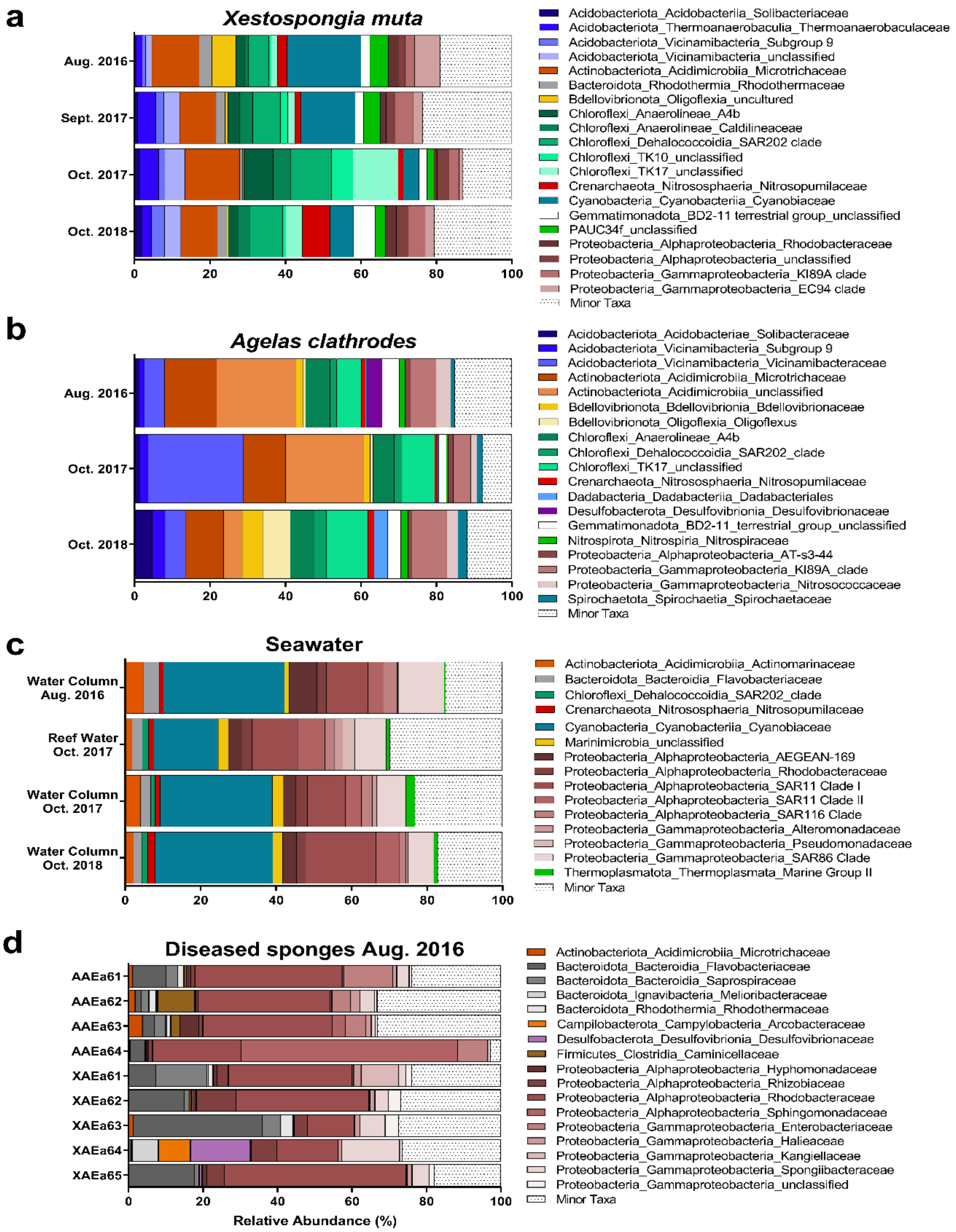


**Supplemental Figure S1. Percent abundance of bacterial sequences for a) heathy *Xestospongia muta*, b) healthy *Agelas clathrodes*, c) Seawater, and d) diseased sponge samples.** Bacterial taxonomy was assigned to Family using the Silva v. 138 database. All taxa < 1% of the total community were grouped under the category ‘Minor Taxa’.

**REFERENCES**


1. Walker DI, McQuillan J, Taiwo M, Parks R, Stenton CA, Morgan H, Mowlem MC, Lees DN. 2017. A highly specific *Escherichia coli* qPCR and its comparison with existing methods for environmental waters. Water Res 126:101–110.
2. Hou A, Laws EA, Gambrell RP, Bae HS, Tan M, Delaune RD, Li Y, Roberts H. 2006. Pathogen indicator microbes and heavy metals in lake pontchartrain following Hurricane Katrina. Environ Sci Technol 40:5904–5910.
3. Sun F, Wu D, Qiu Z, Jin M, Wang X, Li J. 2010. Development of real-time PCR systems based on SYBR Green for the specific detection and quantification of *Klebsiella pneumoniae* in infant formula. Food Control 21:487–491.
4. De Vos D, Lim A, Pirnay JP, Struelens M, Vandenvelde C, Duinslaeger L, Vanderkelen A, Cornelis P. 1997. Direct detection and identification of *Pseudomonas aeruginosa* in clinical samples such as skin biopsy specimens and expectorations by multiplex PCR based on two outer membrane lipoprotein genes, oprI and oprL. J Clin Microbiol 35:1295–1299.
5. Kumar A, Balachandran Y, Gupta S, Khare S, Suman. 2010. Quick PCR based diagnosis of typhoid using specific genetic markers. Biotechnol Lett 32:707–712.
6. Joyner J, Wanless D, Sinigalliano CD, Lipp EK. 2014. Use of quantitative real-time PCR for direct detection of *Serratia marcescens* in marine and other aquatic environments. Appl Environ Microbiol 80:1679–1683.
7. Hussain M, Von Eiff C, Sinha B, Joost I, Herrmann M, Peters G, Becker K. 2008. eap gene as novel target for specific identification of *Staphylococcus aureus*. J Clin Microbiol 46:470–476.
